# Functional community assembly and turnover along elevation and latitude

**DOI:** 10.1101/706523

**Authors:** Marta A. Jarzyna, Ignacio Quintero, Walter Jetz

## Abstract

The drivers of community coexistence are known to vary with environment, but their consistency across latitudes and scales, and resulting conservation implications, remain little understood. Here, we combine functional and phylogenetic evidence along elevations to document strong biotic constraints on coexistence in avian communities in both benign (tropical low elevations) and severely harsh (temperate/polar highlands) environments. Assemblages in both are marked by high assemblage functional uniqueness, whereas in tropical highlands and temperate/polar low elevations there is strong functionally redundancy and pronounced environmental constraints. Only in harsh environments is phylogeny an effective surrogate for functional assemblage structure, reflecting nuanced shifts in the position, shape, and composition of measured multivariate trait space along gradients. Independent of scale and latitude, high elevation assemblages emerge as exceptionally susceptible to functional change.

## Introduction

Species, and the communities they form, convey a range of functions to ecosystems and humans that are now under increasing threat from global change (1). Drivers, presumed mechanisms, and evidence of community change are non-uniform among latitudes and elevations (2–4). This raises fundamental questions about the processes and relative roles of biotic and abiotic drivers underpinning community coexistence along large-scale spatial and environmental gradients and their implications for the maintenance of community functions (5–8). While biotic constraints such as competitive constraints on trait equivalencies, i.e., limiting similarity (9), or certain facilitative intercations involving unique species (10–12) are expected to increase divergence (i.e., overdispersion) of trait characteristics among community members, particularly for closely related species, environmental constraints (or filters) that select for common phenotypes might decrease trait divergence and enhance trait similarity (i.e., clustering) (13). The latter are expected to dominate toward harsh and less stable environments (13, 14), whereas biotic constraints, particularly competitive exclusion, should be more prominent in productive and stable settings (15, 16). Despite their key role for gauging the functional consequences of climate change, whether and how these processes and resulting patterns hold from local to regional/global scales, where macroevolutionary constraints and contingencies on clade functional space emerge, remains largely unclear (17–19) or reliant on phylogenetic proxies of uncertain surrogacy value (20–22).

Here, we use the natural experiment provided by elevational gradients of the world’s main mountain regions replicated along latitude (23, 24), a time-calibrated phylogeny (25), comprehensive trait data (26), and elevational distributions (24) addressing nearly all extant bird species, to test how functional structure varies from benign to harsh conditions across local, regional, and global scales. For 8,410 assemblages along elevations from sea level to 7,340 meters, we estimate prevalence of assembly mechanisms using dendrogram-based functional diversity (FD), hypervolume-based functional diversity (FD_H_; Supplementary Material), and species’ local functional distinctness (FDI) which captures the distinct contribution species make to the total functional diversity of a local assemblage (27). We use their species richness-controlled values, cFD and cFDI (standardized effect sizes), supported by quantile scores and associated p-values, to distinguish overdispersion from clustering. To investigate the evolutionary underpinnings of species coexistence and perform a widely called for test of phylogenetic and functional structure surrogacy (20, 22, 28) we use comprehensive phylogenetic information for all assemblages. Finally, we assess the shape, position, and turnover in assemblage’s multivariate trait volume and decompose it into its constituent elements (single traits) to gain a more mechanistic understanding of assemblage functional structure (29, 30).

## Results

### Functional structure

Globally, absolute FD and FD_H_ peak at ca. 1,500-2,000m (Fig. 1A, Figs. S1-S2, Table S1). Both low and high elevations exhibit functional overdispersion with higher levels of FD and mean FDI (FDI_avg_) than their species richness would suggest (cFD and cFDI patterns, Fig. 1B,E). In contrast, mid elevations are generally more functionally clustered (Fig. 1B,E). But a strong latitudinal variation in this pattern is evident (Fig. 1, Figs. S1-S3, Table S1). cFD, cFD_H_, and cFDI_avg_ decline along elevation in the tropics (<23.5°) and sub-tropics (23.5-35°), but increase toward high elevations in temperate and polar regions (>35°; Fig. 1B,E, Figs. S1-S2, Table S1). Accordingly, cFD, cFD_H_, and cFDI_avg_ decrease with latitude in low to mid elevations, but this trend reverses for high elevations (Fig. 1C,F, Figs. S1-S2, Table S1). Species’ local functional role is highly uneven along elevation and latitude, with a small number of species contributing much more functional uniqueness than the rest in regions with strong environmental constraints (Fig. 1G-I, Figs. S1-S2, Table S1). While the species richness-controlled skewness of species FDI values, cFDI_skew_, increases toward higher elevations in the tropics and sub-tropics, the opposite applies to temperate and polar regions (Fig. 1H, Figs. S1-S2, Table S1). Consequently, cFDI_skew_ increases from the tropics to the arctic at low to mid elevations (Fig. 1I, Figs. S1-S2, Table S1), but declines at high elevations.

**Fig. 1.**
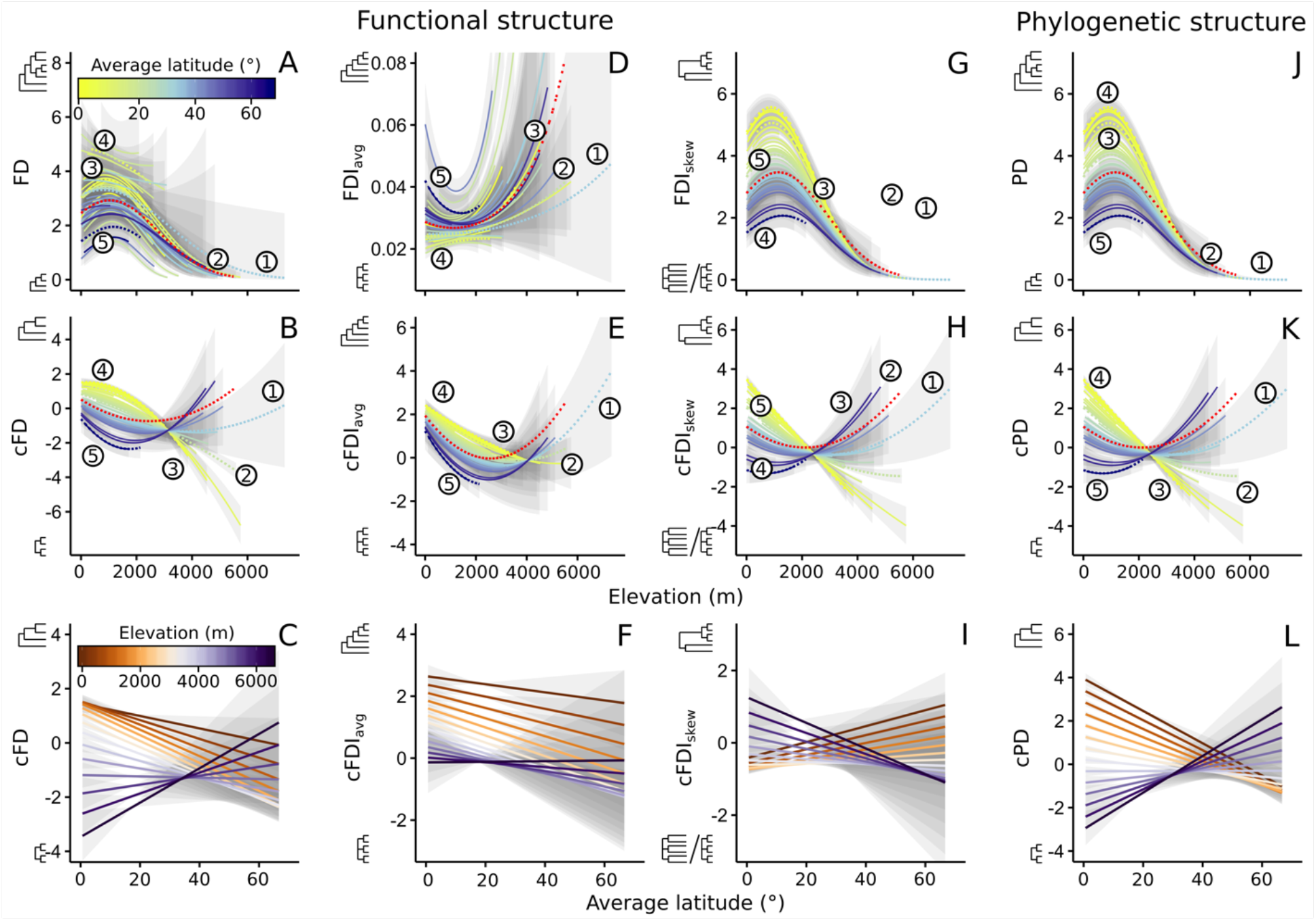
The interaction of central latitude of mountain regions with elevational gradients of avian functional and phylogenetic assemblage structure. Top: fitted elevational trends for dendrogram-based assemblage functional diversity (FD; A), mean (FDI_avg_; D) and skewness (FDI_skew_; G) of species’ local functional distinctness, and dendrogram-based assemblage phylogenetic diversity (PD; J). Middle: fitted elevational trends for the species richness-controlled FD (cFD; B), FDI_avg_ (cFDI_avg_; E), FDI_skew_ (cFDI_skew_; H), and PD (cPD; K), measured using standardized effect sizes. Bottom: latitudinal trends for cFD (C), cFDI_avg_ (F), cFDI_skew_ (I), and cPD (L), given by predictions from models in (B, E, H, and K). Fitted global patterns, with mountain ranges included as random effects in the model, are shown in dashed red line. cFD, cFDI_avg_, and cPD values >0 and <0 indicate overdispersion and clustering, respectively; cFDI_skew_ values >0 and <0 indicate more and less even distribution of species FDI than expected by chance. Values of PD (in MYA) were scaled by 1/1000. Five highlighted in dashed line regions are the Hindukush-Himalaya (1), the Andes (2), the Sumatran Islands (3), the Cameroon Mountains (4), and the Scandinavian Mountains (5) ranges. Grey areas are 95% credible intervals. For details on multi-level models see Materials and Methods.

### Phylogenetic and functional structure surrogacy

Absolute (PD) and richness-controlled (cPD) phylogenetic diversity follow elevational and latitudinal trends broadly similar to those of function (Fig. 1J,K,L, Fig. S1, Table S1). The phylogenetic structure of an assemblage provides a reasonable proxy for the functional structure, but in a strongly latitude-dependent way (Fig. 2C,D, Table S2). In the tropics, the phylogenetic structure offers reasonable, but not strong, surrogacy for functionally overdispersed (frequency, Fr, with which cPD>0 sites are also cFD>0; Fr=0.63) and clustered (frequency with which cPD<0 sites are also cFD<0; Fr=0.62) assemblages (Fig. 2D, Table S2). Toward higher latitudes, the predictive power of phylogenetic structure diverges strongly (Fig. 2D, Table S2), reaching respectively Fr=0.21 and Fr=0.81 for functionally overdispersed and clustered assemblages in temperate and polar regions. This general latitudinal pattern of the predictive ability of the phylogeny is upheld by quantile scores and associated estimated p-values (two-tailed test, *α*=0.05; Fig S5, Table S2).

**Fig. 2.**
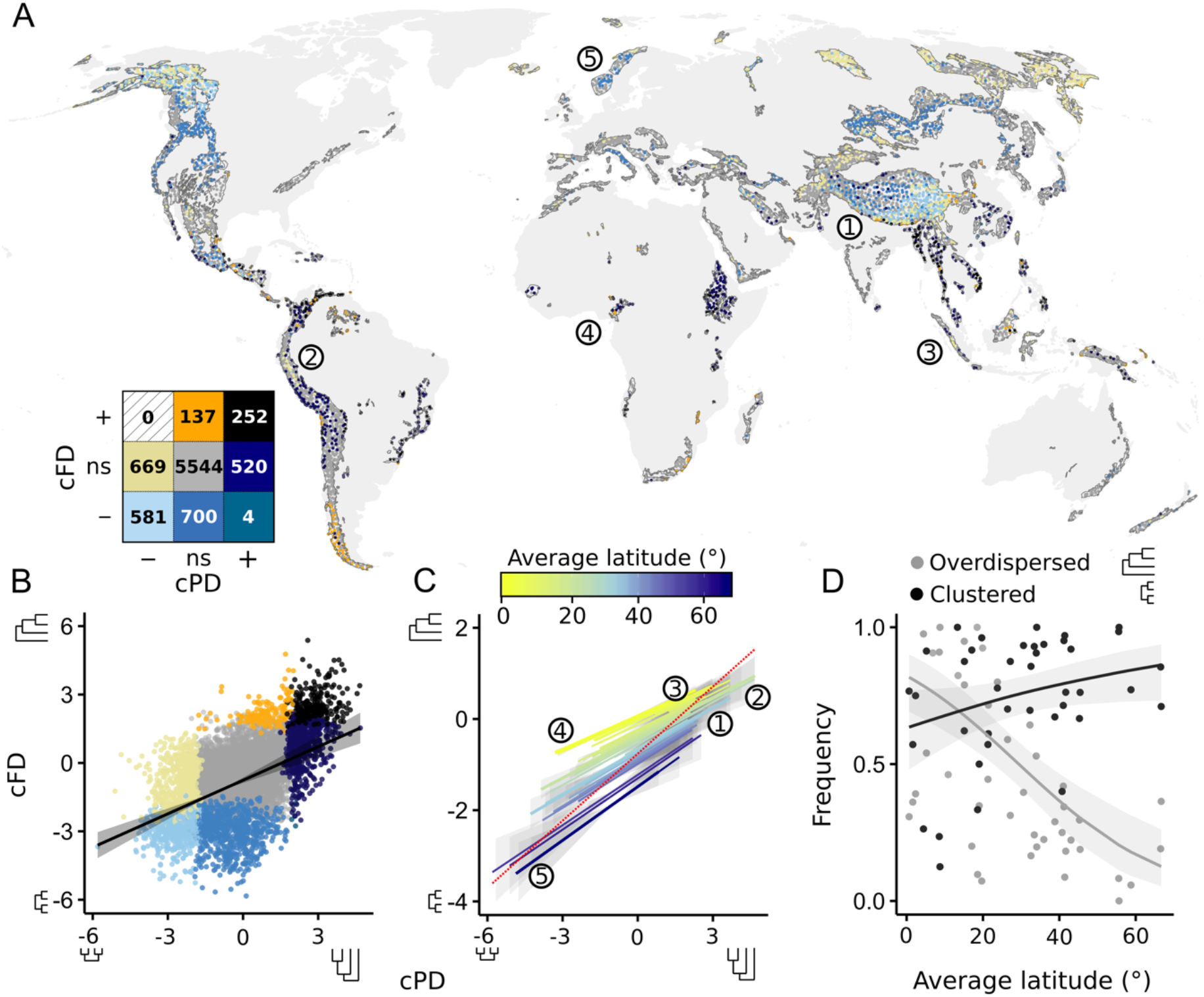
Interaction of phylogenetic and functional structure of all 8,410 analyzed assemblages in mountain regions worldwide across the latitude. Structure is given as species richness-controlled phylogenetic (cPD) and functional (cFD) diversity (standardized effect sizes). Statistically significant deviations in (A) and (B) are denoted by + (overdispersion), – (clustering); ns denotes non-significance; numbers in the contingency table show the number of sites with a statistically significant deviation. (C) illustrates the interaction of central latitude of mountain regions with the relationship between cPD and cFD, with fitted aggregate global pattern shown in dashed red line. (D) illustrates the latitudinal gradient of the frequency with which functional structure is accurately predicted from phylogenetic structure alone. For highlighted regions see Fig. 1.

### Turnover and decomposition of the multivariate trait volume

Globally, the volume of assemblage traits and its relative (centroid) location vary little up to ca. 3,500m elevation, but both shift drastically above that (Fig. 3A,B), suggesting strong across elevation assemblage trait volume redundancy from low to mid elevations and increasing uniqueness above. This is confirmed by the global pattern of overlap in assemblage trait volume (as Sørensen similarity), FD_O_, whose rapid decline toward high elevations (Fig. 3C) coincides with the global peak in functional clustering (at ca. 2,000m; Fig. 1). This global pattern mirrors those in tropical mountain systems, but not those in temperate and polar regions where FD_O_ slightly increases toward higher elevations (Fig. 3C). Simpson’s similarity, FD_T_, remains high and relatively constant along elevation both globally and in the tropics, but increases slightly in higher latitude regions (Fig. 3D). Fraction of trait volume unique to lower elevation assemblages (FD_Ul_) increases strongly toward higher elevations globally and in the tropics, but this increase is less pronounced for temperate and polar regions (Fig. 3E). Fraction of trait volume unique to higher elevation assemblages (FD_Uh_) declines slightly along the elevational gradient globally and for high latitude regions, but shows a slight humped pattern in the tropics (Fig. 3F).

**Fig. 3.**
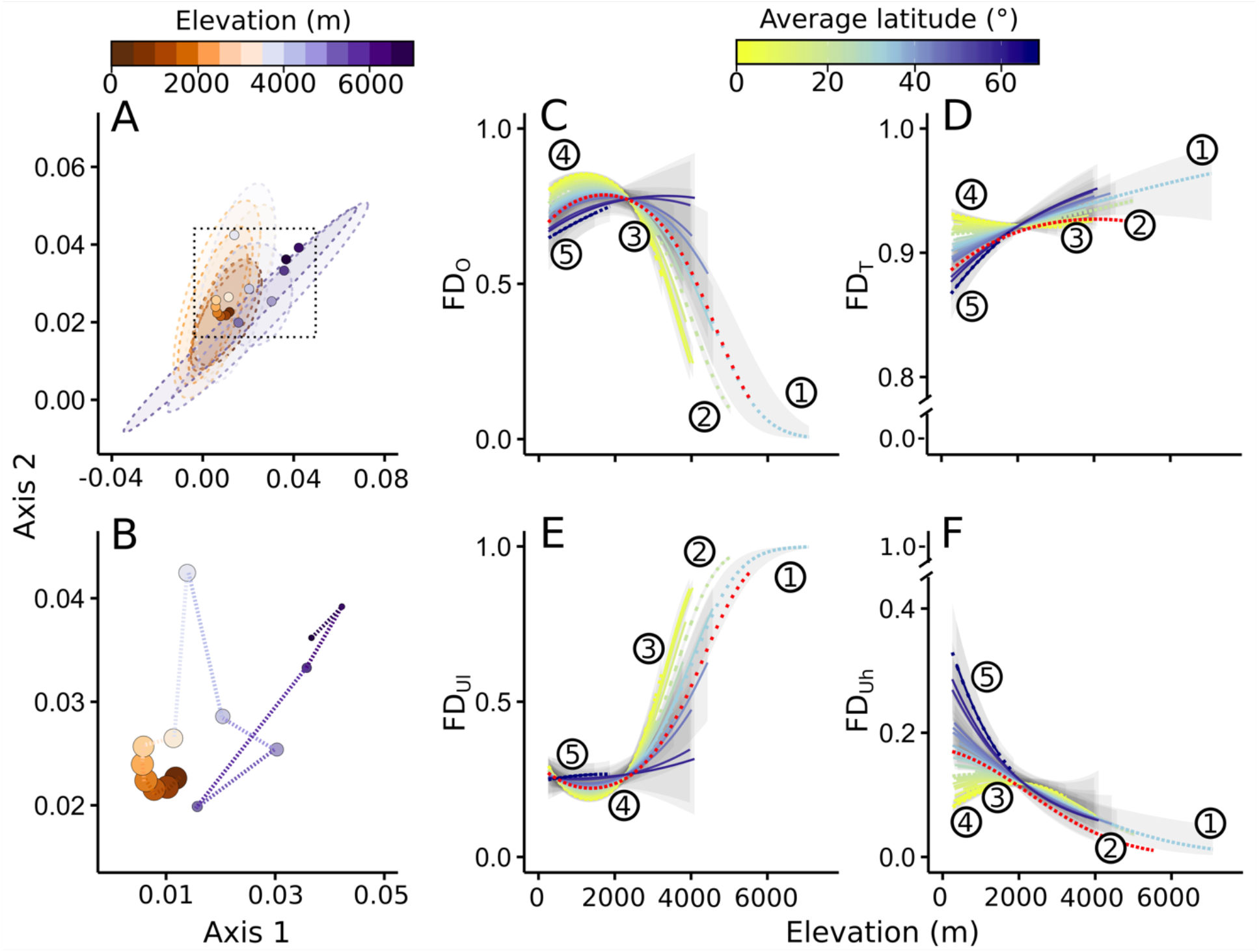
Elevational gradients of assemblage trait space. (A,B) provide the two-dimensional global trait space resulting from Principal Coordinate Analysis (PCoA); points are the centroids of the trait space specific to each consecutive elevational band of 500m intervals averaged over all mountain regions, with 75% confidence ellipses encompassing assemblages at each elevational band (A) and their trajectory toward higher elevations (B). Size of points in (B) illustrates the number of mountain regions informing each elevational band, decreasing with elevation from 42 at 500m to 19 at 4,000m to 1 at 6,000m. (C-F) show the interaction of central latitude of mountain regions with elevational gradients of the overlap in trait volume (FD_O_, Sørensen similarity; C), turnover (FD_T_, Simpson’s similarity; D), and fractions of trait volume unique to assemblages located at lower (FD_Ul_; E) and higher (FD_Uh_; F) elevations. In (C-F), fitted aggregate global patterns are shown in dashed red line. For highlighted regions see Fig. 1.

The variation in avian trait volume along elevation and latitude we uncover is underpinned by a strongly expected (Fig. S6) change in the prevalence of select traits (Fig. 4, Figs. S7,S8). Certain functional components—e.g., mid- and upper-canopy and open-water foragers—generally decline drastically with increasing elevation (Fig. 4B, Figs. S7,S8). Nocturnal species also disappear toward higher elevations (Fig. 4E), which we attribute to harsher night-time temperatures and decreased daytime competition. The prevalence of birds of prey, granivorous, and ground foraging birds increases steeply linearly with elevation in temperate and arctic regions, but closer to the equator shows a mid-elevation peak and a steep decline thereafter (Fig. 4D, Figs. S7,S8). The proportion of insectivorous species increases with elevation in the tropics, but not in temperate and arctic regions, suggestive of lower insect abundance at high elevations there (Fig. 4C). Average assemblage body mass declines from lower to mid elevations and then, at least in higher latitudes, strongly increases toward high elevations (Fig. 4F), consistent with Bergmann’s rule type pattern (31).

**Fig. 4.**
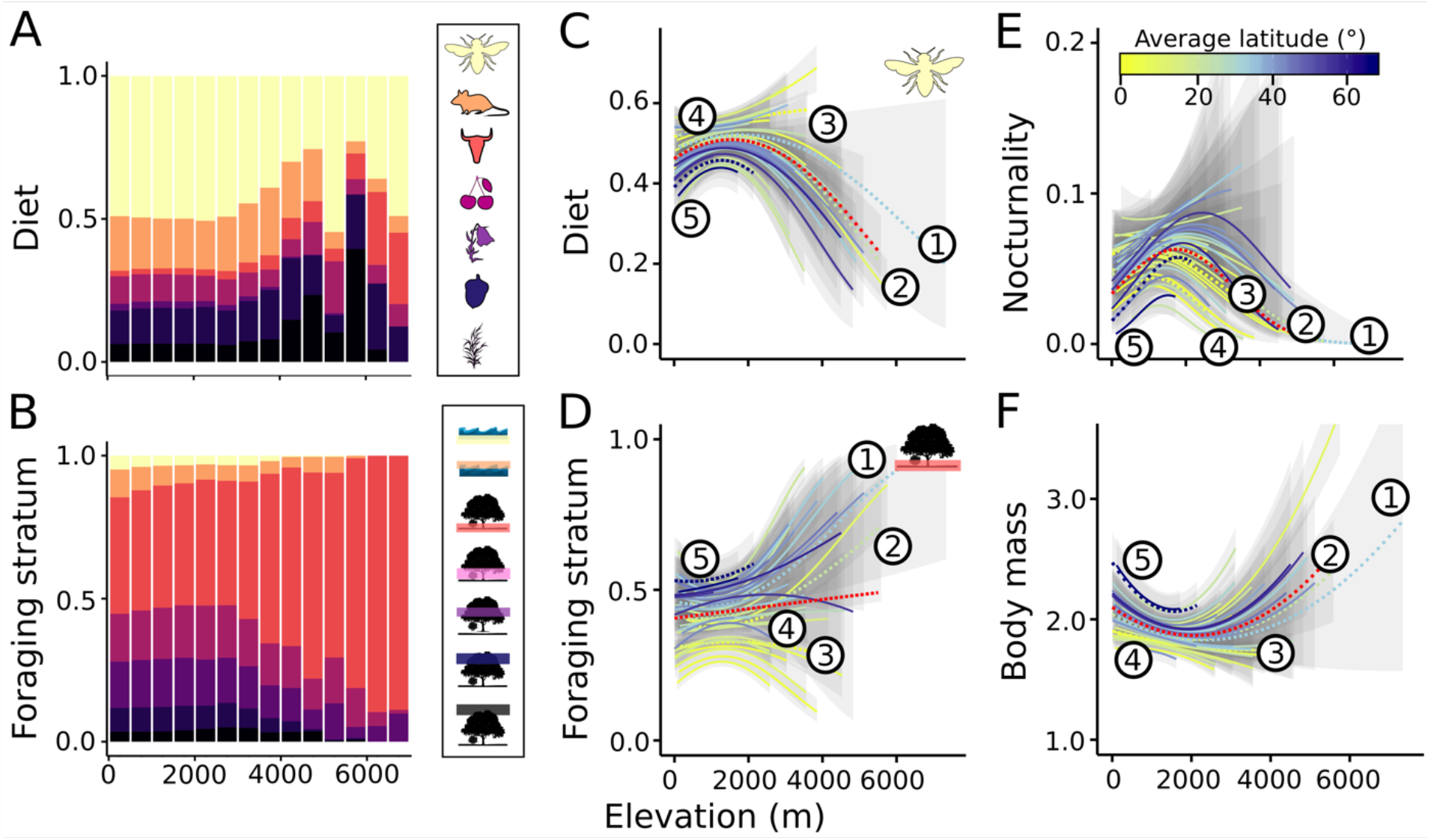
Elevational gradients in the specific aspects of trait space. Shown are all components of the (A) dietary and (B) foraging stratum axes, (C-F) selected individual traits—proportions of insectivorous diet (C), ground foragers (D), nocturnality (E), and average body mass (F). (C-F) illustrate the interaction of central latitude of mountain regions with elevational gradients of the given functional attribute. In (C-F), fitted aggregate global patterns are shown in dashed red line. For highlighted regions see Fig. 1.

## Discussion

Our results confirm our expectation around a prominence of biotic interactions (as suggested by overdispersion) in more productive and stable environments (i.e., tropical low elevations). Correspondingly, environmental filtering dominates in less benign tropical highlands as well as across most of the elevational gradient in temperate and polar regions, characterized by the strong selective force of their harsher winters. This mirrors earlier findings on the increasing role of abiotic contraints leading to phylogenetic clustering (32–34) and faster rates of speciation (24) at high elevations. Contrary to our predictions, however, we find increasing functional overdispersion toward the highlands in temperate and arctic regions. This is consistent with findings of functional overdispersion at both ends of the resource availability spectrum (35). We suggest that in the already harsh conditions of higher latitudes inhabited by a restricted pool of clades, higher elevations impose severe limitations where facilitative interactions (35), potentially paired with competitive exclusion, becomes predominant. We elucidate more of the mechanisms of community assembly by showing that local functional redundancy drives the centers of functional clustering (tropical highlands and temperate low to mid elevations), while overdispersion (the tropical low to mid elevations and temperate highlands) is driven predominantly by functionally unique species. The predominance of functionally unique species in temperate and polar highlands is indeed consistent with increased facilitation as facilitative interactions in stressful environments tend to involve primarily functionally distinct species (5, 10, 11, 36).

A time-calibrated species-level phylogeny allowed us to investigate the evolutionary underpinnings of species coexistence. High joint phylogenetic and functional overdispersion, i.e., when assemblages are comprised of distantly related species with significant trait differences, suggests that only distantly related lineages can evolve sufficient niche differences to overcome competitive dynamics and attain coexistence. In contrast, functional overdispersion coupled with phylogenetic clustering indicates that closely related species readily evolve trait dissimilarities to avoid competition. Functional and phylogenetic clustering, on the other hand, means that closely related species share adaptations that allow them to persist in local environmental conditions. Finally, functional clustering in hand with phylogenetic overdispersion implies that distantly related species adapt convergently to the local environment. Our results suggest that the interplay of these ecological and evolutionary processes of community assembly shows strong elevational and latitudinal variation.

Specifically, in tropical low to mid elevations, where phylogenetic overdispersion offers reasonable surrogacy for functional overdispersion, distantly related species might often coexist thanks to having evolved significant trait dissimilarities. As elevation in the tropics increases, phylogeny remains a reasonable predictor of the functional structure, suggesting that close relatives often share trait combinations conferring the ability to tolerate increasingly harsh environmental conditions of the highlands. With increasing latitude, phylogeny provides a reliable proxy only for clustered assemblages, again suggesting that closely related species that occur in low and mid elevations share phylogenetically conserved adaptations allowing them to persist in stressful environmental conditions of temperate and polar regions. In contrast, functional overdispersion that predominantly characterizes temperate and polar highlands cannot be inferred from the phylogenetic structure, implying that close relatives in the highlands of high latitude regions evolve niche differences to avoid competition. These results suggest that phylogeny might be a reasonable substitute for assemblage functional composition in places where harsh environmental conditions constrain assemblages to a few, functionally similar species. However, the relationship is inconsistent or weak for assemblages where highly functionally unique species prevail. In such settings, it is ill-suited for identifying biotic constraints as a dominant assembly mechanism and for recognizing regions with particularly distinct ecosystem-relevant functions.

The inconsistent predictive power of phylogeny highlights how the mechanistic interpretation of assemblage functional structure across scales benefits from an assessement of its composition and turnover, down to the level of single traits and their respective drivers (29, 30). Globally, we find a strong among-assemblage trait volume redundancy from low to mid elevations and increasing uniqueness above. We attribute the escalating dissimilarity in trait volumes along elevation to decreases in species richness and its associated functional diversity (i.e., nestedness *sensu* (37)) rather than turnover in actual species identities or their functions. As elevation increases, species holding functions reflecting elevational, treeline or terrain, dependencies of associated habitats drop out from assemblages. However, strong latitudinal variation in functional dissimilarity patterns and select underlying traits exists. In the tropics, increasingly depauperate assemblages are functionally nested along elevation as environmental filtering becomes predominant toward higher elevations. In temperate and arctic regions, ubiquitous strong environmental filtering results in relatively stable among-assemblage functional similarity along elevation.

Inferring mechanisms of community assembly from observational data is admittedly challenging in the absence of additional evidence from experimental manipulation (14, 38). Nonetheless, afforded by the concurrent examination of the functional and phylogenetic structure, our analysis provides a powerful test of assembly processes across scales. The additional assessement of the compositional turnover in trait space paired with detailed examination of change in multiple traits along a demonstrable environmental gradient (14) further allows disentangling the mechanisms that might otherwise be masked by multivariate patterns (35).

The uncovered patterns of functional community assembly and turnover have important conservation implications. At local scales, high assemblage functional uniqueness paired with low species richness makes temperate and polar highlands disproportionately susceptible to the loss of critical ecological functions. At larger scales, sustaining the regional multi-functionality of ecosystems requires high dissimilarity among assemblages (39, 40) because no single, local assemblage can support all ecosystem processes (41). We find such high among-assemblage functional uniqueness for tropical and sub-tropical highland ecosystems, causing them to be exceedingly susceptible to disruption of their large-scale functioning. Globally, different processes and scales combine to make high elevation ecosystems exceptionally prone to future ecological perturbations and loss of functions.

## Materials and Methods

### Mountain systems

To delineate mountain regions, we used the inventory of the World’s mountain regions based on expert delineation and terrain ruggedness provided by the Global Mountain Biodiversity Assessment (GMBA, http://www.mountainbiodiversity.org). The GMBA inventory identified 46 broad-scale mountain regions across the five continents. (24) evaluated the biological independence of the GMBA delineation and found that only a small proportion of species is shared among the 46 regions (median similarity of 0.47% +− 10.37% (sd)).

### Distributions and elevational ranges and biodiversity sampling

Data on breeding distributions were compiled from the best available sources for a given broad geographical region or taxonomic group and totaled 9993 species (for individual maps, see https://mol.org). The database of bird elevational ranges was compiled by (24) and is available at https://mol.org. Using these distributional and elevational ranges, we then followed the protocol established by (24) to sample biodiversity data. The sampling procedure returned a total of 21,655 assemblages across the world and 8,410 assemblages within our delimited mountain systems with a median of 71 per mountain region (minimum 3 in the Central Australian mountains, maximum 1,816 in the Himalayas). While admittedly coarse-scale and solely presence-absence, the distributional data used here comprise the best current understanding of bird distributions at the global scale.

### Avian phylogenetic and functional diversity, and trait space

For each assemblage, we calculated avian dendrogram-based functional diversity (FD) (42). We based estimates of FD on a compilation of function-relevant traits in (26). Four trait categories were included: body mass, nocturnality, diet, and foraging niche. The diet and foraging niche categories included seven axes each: proportions of invertebrates, vertebrates, carrion, fresh fruits, nectar and pollen, seeds, and other plant materials in species’ diet (diet category); proportional use of water below surface, water around surface, terrestrial ground level, understory, mid canopy, upper canopy and aerial (foraging niche category). Following existing practice (2, 43) we calculated multivariate trait dissimilarity using Gower’s distance for each pairwise combination of all 9,993 species in the dataset. Equal weights were given to each of the trait categories and to each axis within the trait categories (i.e., each diet and foraging niche variable was given a 1/7 weight, whereas the weights of body mass and nocturnality was 1). We used Gower’s distance as distance metric because this index can handle quantitative, semi-quantitative, and qualitative variables and assign different weights to individual traits (43).

The functional dendrogram was built using UPGMA (Unweighted Pair Group Method with Arithmetic Mean) clustering. UPGMA clustering has the highest cophenetic correlation coefficient among most popular clustering methods (Ward, Single, Complete, WPGMA, WPGMC, and UPGMC clustering methods) and the lowest 2-norm index (44), ensuring most faithful preservation of the original distances in the dissimilarity matrix. At each of the point locations, the functional dendrogram was pruned of the branches for species that did not occur at that location to reflect that location’s species composition. FD_D_ was calculated as a sum of branch lengths of such local functional dendrogram.

The calculation of dendrogram-based phylogenetic diversity (PD) followed the same procedure (45), but instead of a functional dendrogram, we used 20 dendrograms sampled from full pseudo-posterior distribution of phylogenetic trees assembled by (25) (http://birdtree.org/). PD was calculated as the total branch length of tree branches averaged over the 20 phylogenetic trees, with 20 trees thought to provide a sufficiently strong initial estimate (46).

We used the same local functional dendrogram to quantify the assemblage mean (FDI_avg_) and skewness (FDI_skew_) of species’ local functional distinctness (FDI). FDI was calculated as the fair proportion branch length of the branches in a local functional dendrogram leading to a species tip and measures the uniqueness of a species functional characteristics in the context of all other species in the same assemblage. As such, FDI decomposes overall FD into individual species contributions. When averaged across the entire assemblage, FDI_avg_ informs on how the average distinct functional position of single species changes along gradients (FDI_avg_). High FDI_avg_ indicate on average a given assemblage is comprised of functionally unique species. Skewness of the distribution of FDI values further informs on whether individual species contribute uniformly or unevenly to overall FD (low and high FDI_skew_, respectively).

The clustering algorithm can affect the shape of the functional dendrogram and, consequently, the estimate of the dendrogram-based functional diversity (47). To ensure that our choice of the clustering algorithm did not significantly alter the inferences, we further quantified functional diversity directly with the multidimensional functional space. Specifically, we calculated functional diversity as the assemblage trait volume (FD_H_), quantified using the hypervolume metric (48). To obtain FD_H_, we first conducted the Principal Coordinate Analysis (PCoA) using the multivariate trait dissimilarity matrix based on Gower’s distance for each pairwise combination of all species in the dataset. We then used the first two principal coordinates to quantify the hypervolume of each point location using the minimal convex hull method in the package hypervolume (48). Using more than two principal coordinates was computationally unfeasible because the algorithm becomes exponentially inefficient in high dimensionalities. Hypervolumes quantified in this manner reflected positions of all species present at a given assemblage. Results for FD_H_ are provided in the Supplementary Material (Fig. S2).

To further evaluate how assemblage trait volume changes with elevation, we quantified an overlap in trait volume (FD_O_) among the point locations falling along the elevational gradient using package hypervolume (48). For each point at an elevation *a*, we selected the geographically closest point that lies within (*a*+500m, *a*+1000m) elevational region. Because we wanted to evaluate changes in trait volume along elevation rather than changes that might arise from geographic distance, we excluded all pairs of points that were geographically farther apart than 500km. For each pair of points we quantified niche overlap using four metrics: Sørensen (FD_O_) and Simpson’s (FD_T_) similarity and unique fractions of trait volume for points at lower (FD_Ul_) and higher (FD_Uh_) elevation (Blonder et al., 2014). To obtain these indices, we first conducted the Principal Coordinate Analysis (PCoA) using the multivariate trait dissimilarity matrix based on Gower’s distance for each pairwise combination of all species in the dataset. We then used the first two principal coordinates to quantify the hypervolume of each point location using the minimal convex hull method in the package hypervolume (48). Sørensen similarity was then given as twice the intersection of trait volumes of the assemblage at lower elevation and that at upper elevation divided by volume of both assemblages. Unique fractions were calculated as the unique component of the trait volume of, respectively, lower and higher elevation assemblage divided by the entire trait volume of that assemblage. Simpson’s similarity (not provided by the hypervolume package) was derived from unique fractions indices and equaled to the intersection of trait volume of assemblage at lower elevation and that at higher elevation divided by trait volume of the assemblages with a smaller unique fraction (37). We conducted sensitivity analysis by repeating the above steps for points falling within (*a*+250m, *a*+750m) and (*a*+750m, *a*+1250m), and for pairs of points located geographically not farther than 100km and 250km, and present these results in the Supplementary Material (Fig. S9).

We also assessed individual components of avian trait volume. To separate trait axes, or guilds, we used dietary niche (i.e., proportions of different types of food in species’ diet—invertebrates, vertebrates, carrion, fresh fruits, nectar and pollen, seeds, and other plant materials), foraging niche (i.e., the proportional use of each of seven niches—water below surface, water around surface, terrestrial ground level, understory, mid canopy, upper canopy, and aerial), nocturnality, and log-transformed body mass. For dietary and foraging niche, we quantified the relative proportion (i.e., prevalence) of each diet and foraging niche for each assemblage. To account for the effect of separate mountain ranges, we first assessed individual components of avian trait volume for each mountain range and then averaged those values to obtain a global estimate.

### Richness-controlled avian phylogenetic and functional diversity, and trait space

Multivariate components of phylogenetic and functional diversity are often closely associated with species richness (SR), and interpretations of assemblage structure benefit from statistically controlling for this association. To evaluate whether elevational gradients in phylogenetic, functional, and trait diversity deviate from the expectation given SR, we generated null model for PD, FD, FDI_avg_, FDI_skew_, FD_H_, and individual components of avian assemblage trait volume. Even though FDI values provide estimates of individual species contributions to FD, their assemblage-level means (FDI_avg_) and skewness (FDI_skew_) are often closely associated with SR (27). Consequently, species richness-controlled equivalents of FDI_avg_ and FDI_skew_ allow more in-depth interpretations of the mechanisms behind the patterns of assemblage structure.

First, we developed the expected values of all metrics by randomly selecting species from a regional species pool (i.e., species that occur in a given mountain region), keeping point-level SR constant (49). The random selection of species for null models was performed 100 times. We calculated the standardized effect sizes as the difference between observed values and those expected from the null model, expressed in units standard deviation. The standardized effect sizes were considered the species richness-controlled equivalents of biodiversity metrics and were denoted as cPD, cFD, cFDI_avg_, cFDI_skew_, and cFD_H_. Negative and positive values of standardized effect sizes indicated that an observed value was lower and higher, respectively, than the average expected value. Negative and positive values of cPD, cFD, cFDI_avg_, and cFD_H_ suggested clustering and overdispersion, respectively. Negative and positive values of cFDI_skew_ suggested less and more, respectively, even distribution of species functional distinctness values.

Because standardized effect sizes might be affected by the shape of the null distribution, it is recommended to additionally calculate quantile values (49). We thus futher calculated quantile scores for the observed values of PD, FD, FDI_avg_, FDI_skew_, and FD_H_, and estimated their associated p-values (i.e., two-tailed test, *α*=0.05). The observed values falling outside the 2.5% and 97.5% quantiles of the null distribution were considered statistically significantly lower (clustering) and higher (overdispersion), respectively, than expected given SR. While the calculation of quantile scores and the associated p-values removes some of the biases potentially resulting from the the shape of the null distribution, they also remove information regarding the size of the effect itself (49). We thus report both the standardized effect sizes (i.e., cPD, cFD, cFDI_avg_, cFDI_skew_, and cFD_H_) and the quantile scores with associated p-values.

We then quantified frequency (Fr) with which phylogenetic structure predicted functional structure of an assemblage. Fr of successful predictions was calculated separately for overdispersed and clustered assemblages and was given by the proportion of phylogenetically overdispersed (clustered) assemblages that were also functionally overdispersed (clustered). We quantified Fr for both designations of overdispersion and clustering: (a) all positive and negative values of standardized effect sizes were considered to be overdispersed and clustered and (b) observed values outside the 2.5% and 97.5% quantiles of the null distribution were considered to be overdispersed and clustered, respectively. To account for the effect of separate mountain ranges, Fr was averaged per mountain region, and mountain region’s central latitude was used to evaluate the latitudinal gradient in Fr.

### Global elevational gradients of avian functional, phylogenetic, and trait diversity

To explore how PD, cPD, FD, cFD, FDI_avg_, cFDI_avg_, FDI_skew_, cFDI_skew_, FD_H_, and cFD_H_, FD_O_, FD_T_, FD_Ul_, FD_Uh_, and individual components of assemblage trait volume vary with elevation, we used two different models: following a linear relationship,

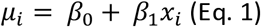

and a quadratic relationship,

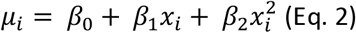

To estimate a global pattern, we used two-level generalized hierarchical model, in which the parameters for each mountain system came from a multivariate normal distribution (for model details, see (24)). Log-transformed PD, FD, FDist_avg_, and FDist_skew_, FD_H_, as well as cPD, cFD, cFDI_avg_, and cFDI_skew_, cFD_H_ followed a Gaussian distribution 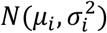. FD_O_, FD_T_, FD_Ul_, FD_Uh_, and each component of avian trait space followed a beta distribution *beta*(*α*_*i*_, *β*_*i*_); with exception of log-transformed body mass, which followed a Gaussian distribution. All predictors were standardized by rescaling the distribution to have mean of 0 and standard deviation of 1. Each model was fitted in a Bayesian framework using INLA (50) through the R-INLA package for R v.3.4.3 (https://www.r-project.org/). To select the best fitting model (quadratic or linear), we estimated the Watanabe–Akaike information criterion (WAIC), which is preferable to similar alternatives because it averages over the posterior distribution rather than using only point estimates (51). We did not use the coefficient of determination, R^2^, to evaluate goodness-of-fit of each model because R^2^ does not integrate across the full posterior probability and thus does not account for uncertainty in parameter estimates.

### Latitudinal influences on elevational gradients of avian functional, phylogenetic, and trait diversity

In order to explore influences of latitudinal position on the elevational gradients in PD, FD, FDist_avg_, FDist_skew_, FD_H_, cPD, cFD, cFDI_avg_, cFDI_skew_, cFD_H_, FD_O_, FD_T_, FD_Ul_, FD_Uu_, and individual components of assemblage trait volume, we expanded the global hierarchical model to include average absolute latitude of the mountain system following (24). Average latitude was fitted as interaction terms with linear and quadratic effects of elevation. All models were run using R-INLA. WAIC was used to select best fitting model.

Lastly, we used the expanded best fitting global model (i.e., one including average latitude as the interaction term) to predict the latitudinal gradients in PD, cPD, FD, cFD, FDI_avg_, cFDI_avg_, FDI_skew_, cFDI_skew_, FD_H_, and cFD_H_ at different elevational bands.

## Acknowledgments

The authors are grateful for support from the Translational Data Analytics Institute at the Ohio State University and Yale Center for Biodiversity and Global Change. The authors also acknowledge support from NSF DEB 1441737, DBI 1262600, DEB 1558568, NASA NNX11AP72G to W.J.

