## Supplementary Material for "Functional community assembly and turnover along elevation and latitude"

Marta A. Jarzyna[1,2], Ignacio Quintero[2,3], and Walter Jetz[2]

[1]Department of Evolution, Ecology and Organismal Biology, The Ohio State University
318 W. 12th Avenue, 300 Aronoff Laboratory, Columbus OH, 43210
[2]Department of Ecology and Evolutionary Biology, Yale University
165 Prospect Street, New Haven, CT, 06520, USA
[3]Institut de Biologie de l'ENS (IBENS), Département de biologie, École normale supérieure, CNRS, INSERM, Université PSL, 75005 Paris, France

Ignacio Quintero:
Walter Jetz:

**Keywords:** biodiversity, birds, community assembly, elevational gradient, functional diversity, latitudinal gradient, traits

**Corresponding author:** Marta A. Jarzyna, Department of Evolution, Ecology and Organismal Biology, The Ohio State University, 318 W. 12th Avenue, 300 Aronoff Laboratory, Columbus OH, 43210.

**Supplementary Material**

**TABLES**

Table S1. Interaction of central absolute latitude of mountain regions with the intercept (*$\beta_0$*), linear (*$\beta_1$*), and quadratic (*$\beta_2$*) effect of the elevation on avian functional and phylogenetic diversity and structure. The results are based on global multilevel models with mountain system as a random effect and absolute latitude of the centroid of the mountain system as a covariate (see text for details). Functional and phylogenetic diversity and assemblage structure are measured with the dendrogram-based avian assemblage functional diversity (FD), the assemblage mean ($FDI_{avg}$) and skewness ($FDI_{skew}$) of species' local functional distinctness underpinning it, the hypervolume-based avian assemblage functional diversity ($FD_H$), and phylogenetic diversity (PD), as well as their species richness-controlled equivalents: cFD, $cFDI_{avg}$, $cFDI_{skew}$, $FD_H$, and cPD (measured with standardized effect sizes). Orange and blue fields indicate statistically significantly, respectively, negative and positive effects (i.e., whose 95% CI did not overlap 0).

| Dependent variable | $\beta_0$ | $\beta_1$ | $\beta_2$ |
|---|---|---|---|
| FD | -0.15 | 0.03 | -0.01 |
| $FDI_{avg}$ | 0.04 | -0.01 | 0.05 |
| $FDI_{skew}$ | 0.00 | -0.14 | 0.00 |
| $FD_H$ | -0.03 | 0.07 | 0.10 |
| PD | -0.21 | 0.07 | 0.01 |
| cFD | -0.75 | 0.47 | 0.45 |
| $cFDI_{avg}$ | -0.56 | 0.08 | 0.19 |
| $cFDI_{skew}$ | 0.00 | -0.31 | -0.05 |
| $cFD_H$ | -0.26 | 0.23 | 0.35 |
| cPD | -0.33 | 0.86 | 0.17 |

Table S2. The frequency with which assemblage phylogenetic structure accurately predicts functional structure globally and across the latitudinal gradient, from tropics to the temperate and polar regions. Clustering and overdispersion are given by (i) species richness-controlled functional (cFD) and phylogenetic (cPD) diversity, measured using standardized effect sizes (SES), of <0 and >0, respectively and (ii) p-values of 0.025 and 0.975, respectively, estimated from the quantile scores for the observed values of FD and PD (see text for details).

| Structure | Model | Frequency | | |
|---|---|---|---|---|
| | | Global | Tropics | Temperate and polar |
| Overdispersed | SES-based | 0.45 | 0.63 | 0.21 |
| | Quantile score-based | 0.26 | 0.40 | 0.00 |
| Clustered | SES-based | 0.74 | 0.62 | 0.81 |
| | Quantile score-based | 0.26 | 0.25 | 0.20 |

**FIGURES**

Fig. S1. The interaction of central latitude of mountain regions with elevational gradients of avian functional and phylogenetic assemblage structure. Top: fitted elevational trends for dendrogram-based assemblage functional diversity (FD; A), mean ($FDI_{avg}$; C) and skewness ($FDI_{skew}$; E) of species' local functional distinctness, and dendrogram-based assemblage phylogenetic diversity (PD; G). Bottom: fitted elevational trends for the species richness-controlled FD (cFD; B), $FDI_{avg}$ ($cFDI_{avg}$; D), $FDI_{skew}$ ($cFDI_{skew}$; F), and PD (cPD; H), measured using standardized effect sizes. Fitted global patterns, with mountain ranges included as random effects in the model, are shown in red dashed line. cFD, $cFDI_{avg}$, and cPD values >0 and <0 indicate overdispersion and clustering, respectively; $cFDI_{skew}$ values >0 and <0 indicate more and less even distribution of species FDI than expected by chance. Values of PD (in MYA) were scaled by 1/1000. Five highlighted in dashed line regions are the Hindukush-Himalaya (1), the Andes (2), the Sumatran Islands (3), the Cameroon Mountains (4), and the Scandinavian Mountains (5) ranges. Grey areas are 95% credible intervals. For details on multi-level models see Materials and Methods.

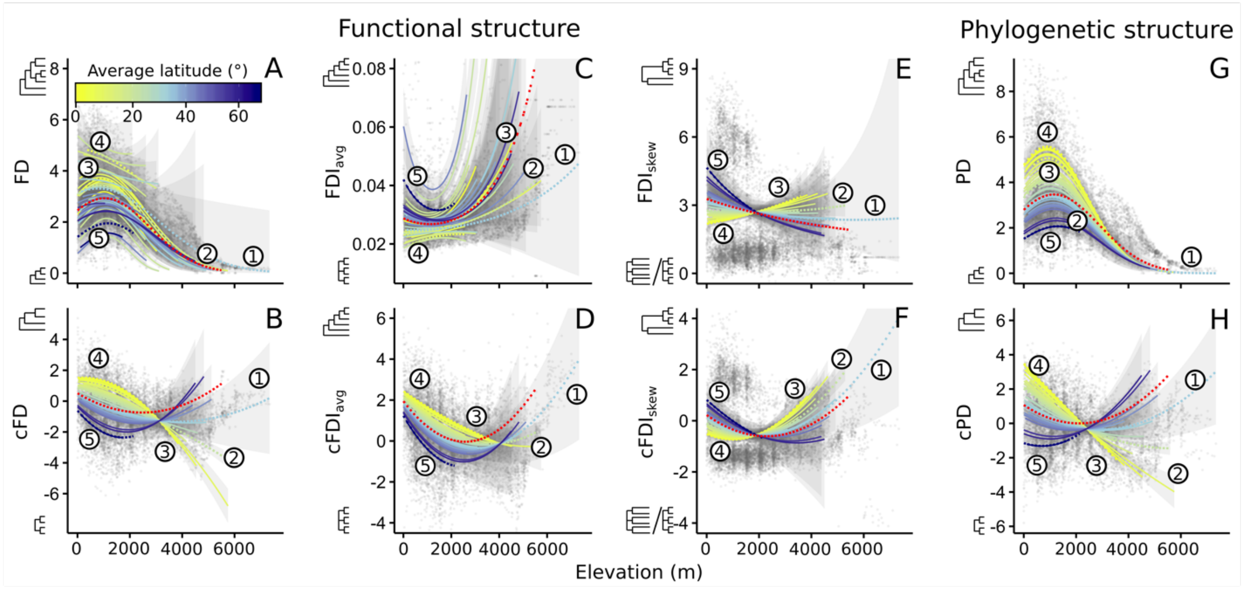

Fig. S2. The interaction of central latitude of mountain regions with elevational gradients of avian functional assemblage structure: (A) fitted elevational trends for hypervolume-based assemblage functional diversity ($FD_H$), (B) fitted elevational trends for the species richness-controlled $FD_H$ ($cFD_H$), measured using standardized effect sizes, and (C) latitudinal trends for $cFD_H$, given by predictions from models in (B). Fitted global patterns, with mountain ranges included as random effects in the model, are shown in red dashed line. $cFD_H$ values >0 and <0 indicate overdispersion and clustering, respectively. Five highlighted in dashed line regions are the Hindukush-Himalaya (1), the Andes (2), the Sumatran Islands (3), the Cameroon Mountains (4), and the Scandinavian Mountains (5) ranges. Grey areas are 95% credible intervals. For details on multi-level models see Materials and Methods.

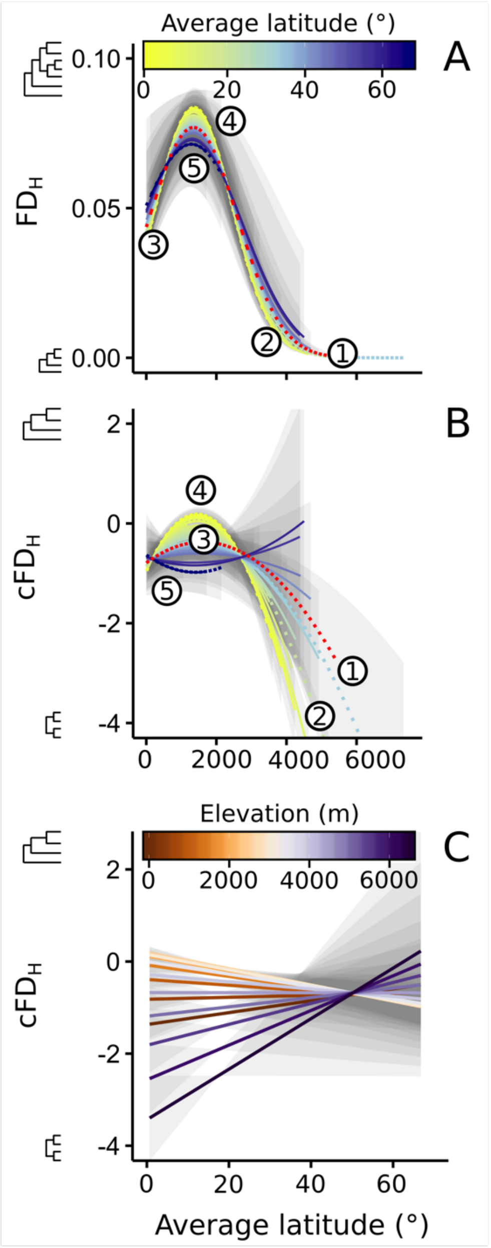

Fig. S3. Mountain-region elevational gradients of (A) dendrogram-based functional diversity (FD), (B) assemblage mean local functional distinctness ($FDI_{avg}$), (C) skewness of species local functional distinctness ($FDI_{skew}$), (A) hypervolume-based functional diversity ($FD_H$), (E) overlap in trait space measured as Sørensen similarity ($FD_O$), true turnover measured as Simpson's similarity ($FD_T$), and unique components of trait space ($FD_U$) for assemblages located at lower ($FD_{Ul}$) and higher ($FD_{Uh}$) elevations, (F) two-dimensional trait space, (G) components of the dietary, (H) foraging, and (I) nocturnality and body mass axes. Trait space in (F) results from Principal Coordinate Analysis (PCoA) and points show the centroids of the trait space specific to each consecutive elevational band of 500m, with 75% confidence ellipses encompassing species at each elevational band.

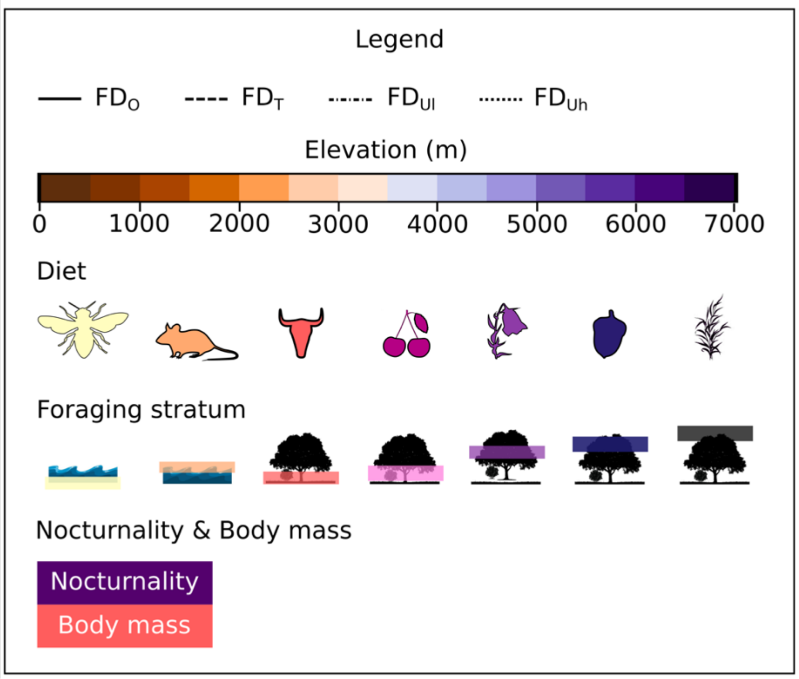

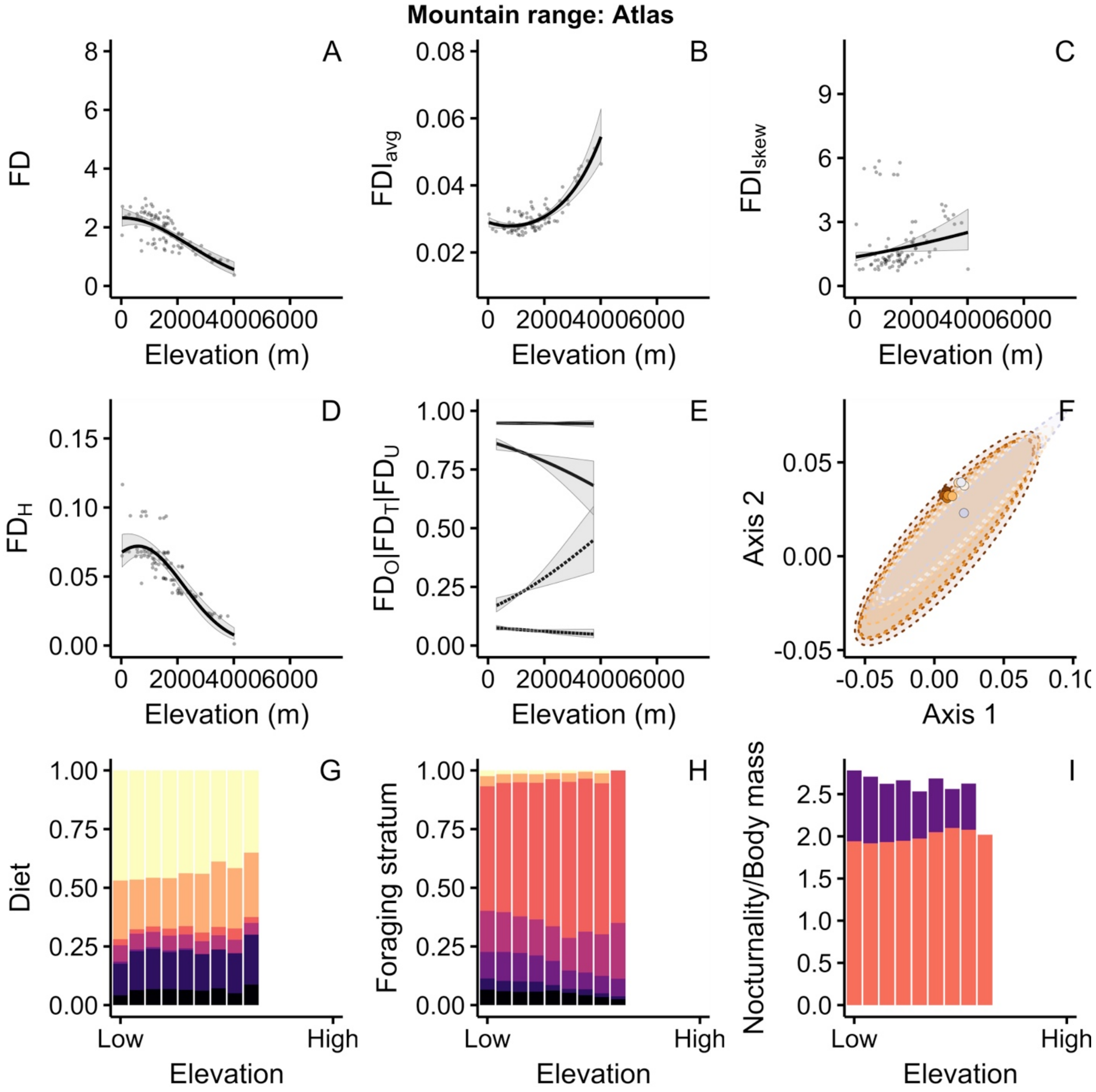
Mountain range: Atlas
A
B
C
D
E
F
G
H
I
FD
$FDI_{avg}$
$FDI_{skew}$
$FD_H$
$FD_O|FD_T|FD_U$
Axis 2
Axis 1
Diet
Foraging stratum
Nocturnality/Body mass
Elevation (m)
Elevation
Low
High

**Mountain range: Sahara**

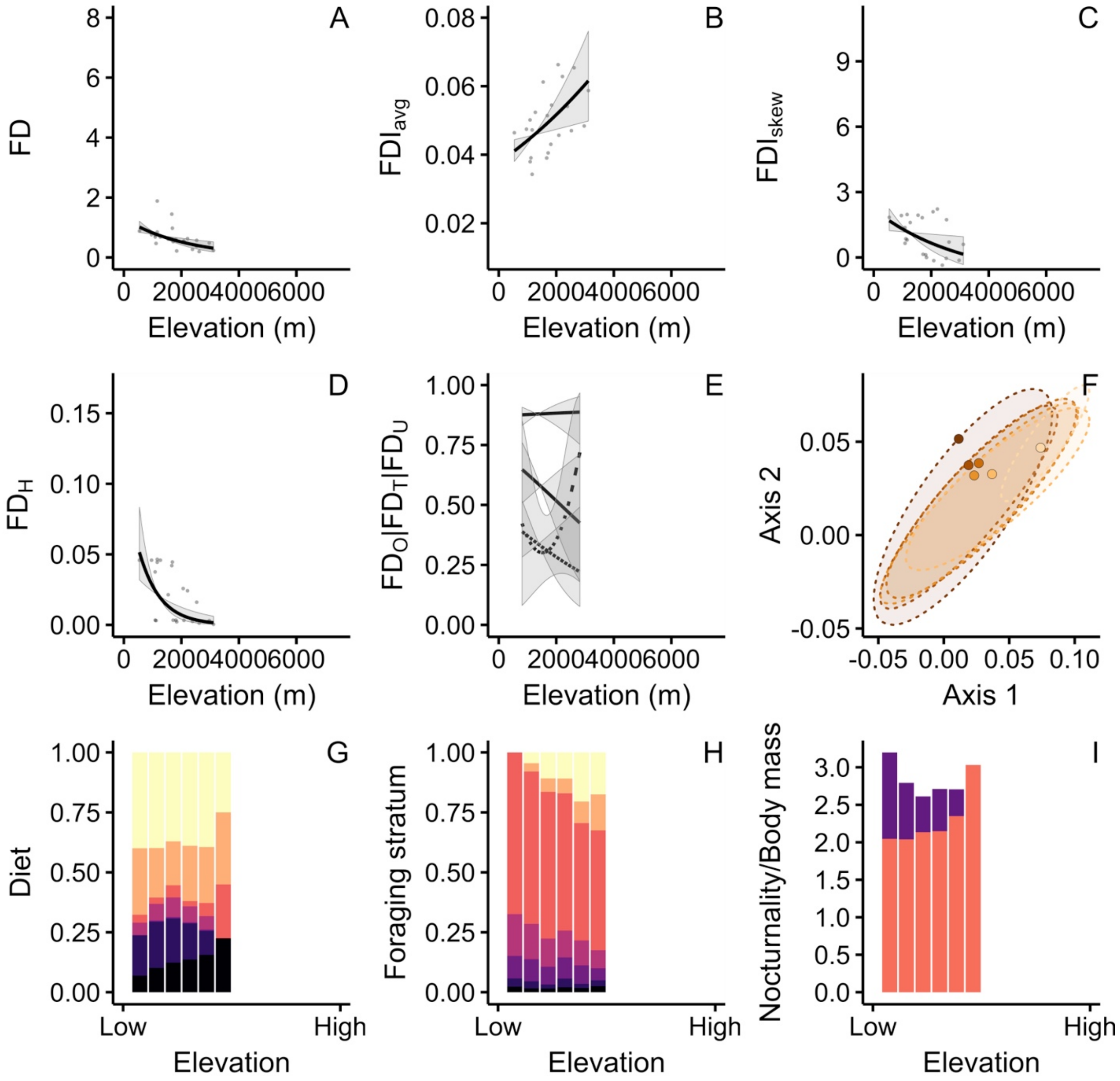

**Mountain range: Great Rift Valley**

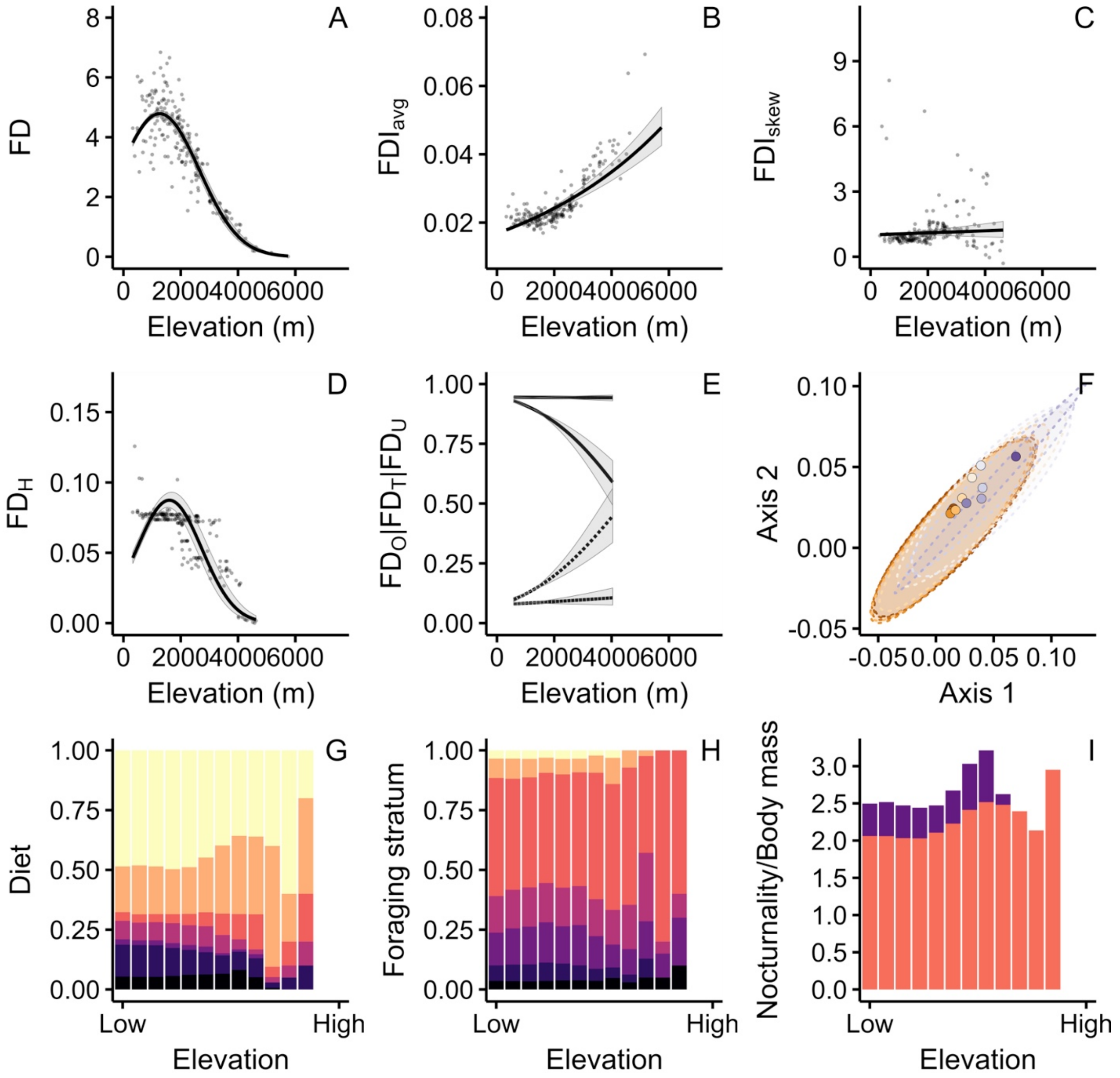

**Mountain range: Cameroon Mountains**

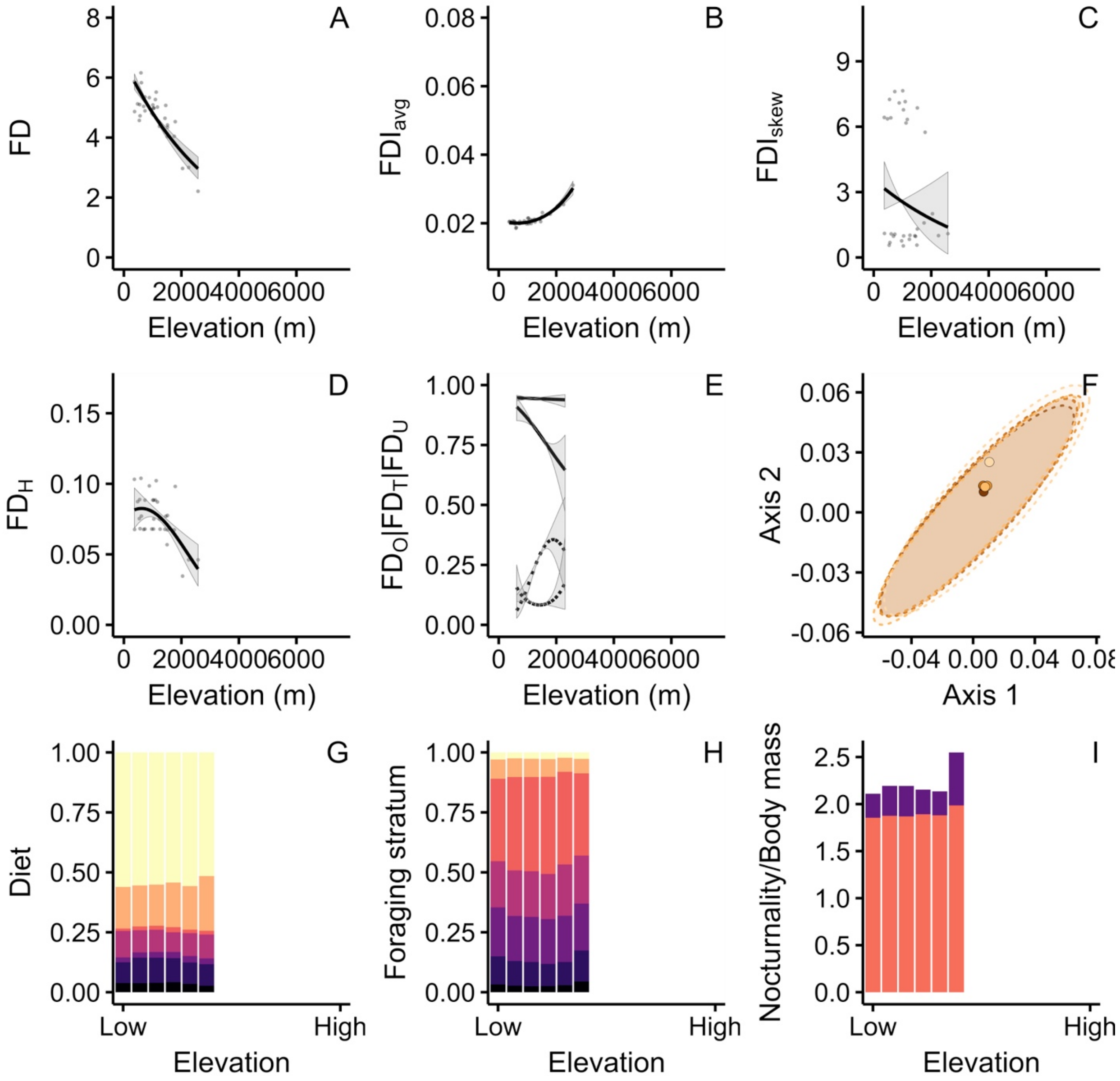

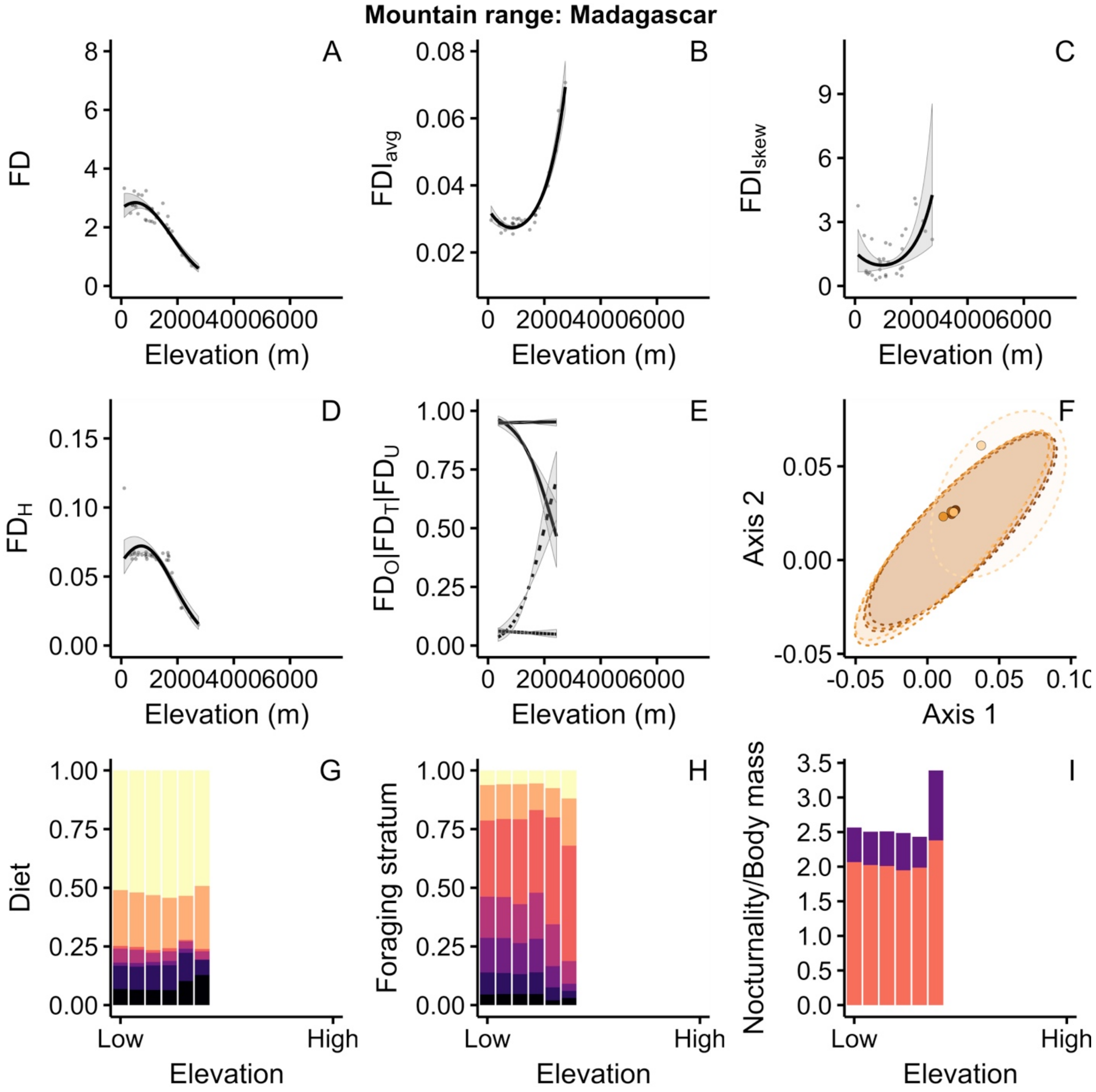

Mountain range: Madagascar
A
B
C
D
E
F
G
H
I
FD
$FDI_{avg}$
$FDI_{skew}$
$FD_H$
$FD_O|FD_T|FD_U$
Axis 2
Axis 1
Diet
Foraging stratum
Nocturnality/Body mass
Elevation (m)
Elevation
Low
High

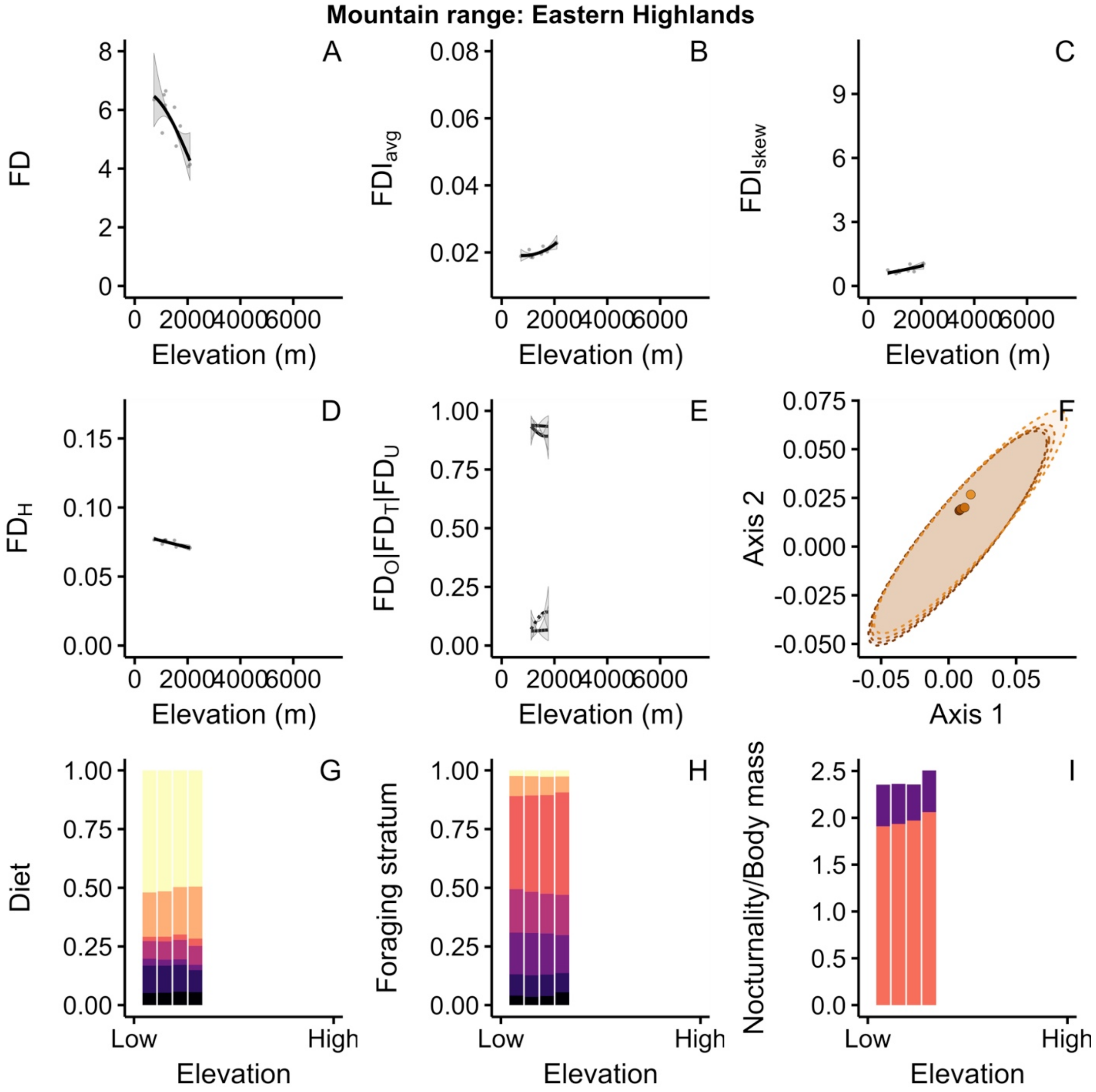

Mountain range: Eastern Highlands
A
B
C
D
E
F
G
H
I
FD
FDI$_{avg}$
FDI$_{skew}$
FD$_H$
FD$_O$|FD$_T$|FD$_U$
Axis 1
Axis 2
Elevation (m)
0
2000
4000
6000
Diet
Foraging stratum
Nocturnality/Body mass
Elevation
Low
High

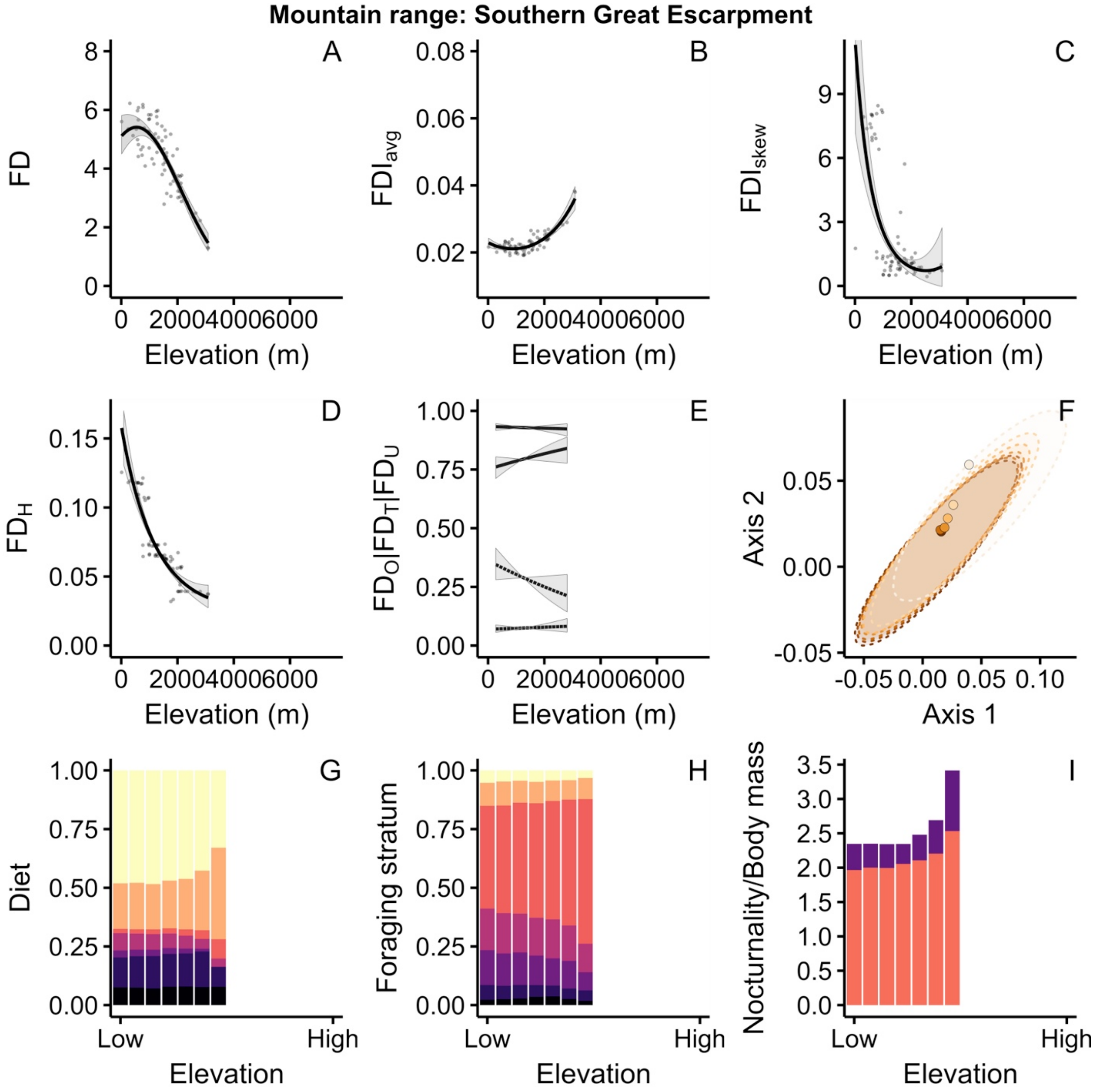

Mountain range: Southern Great Escarpment
A
B
C
D
E
F
G
H
I
FD
FDI_avg
FDI_skew
FD_H
FD_O|FD_T|FD_U
Axis 2
Axis 1
Diet
Foraging stratum
Nocturnality/Body mass
Elevation (m)
Elevation
Low
High

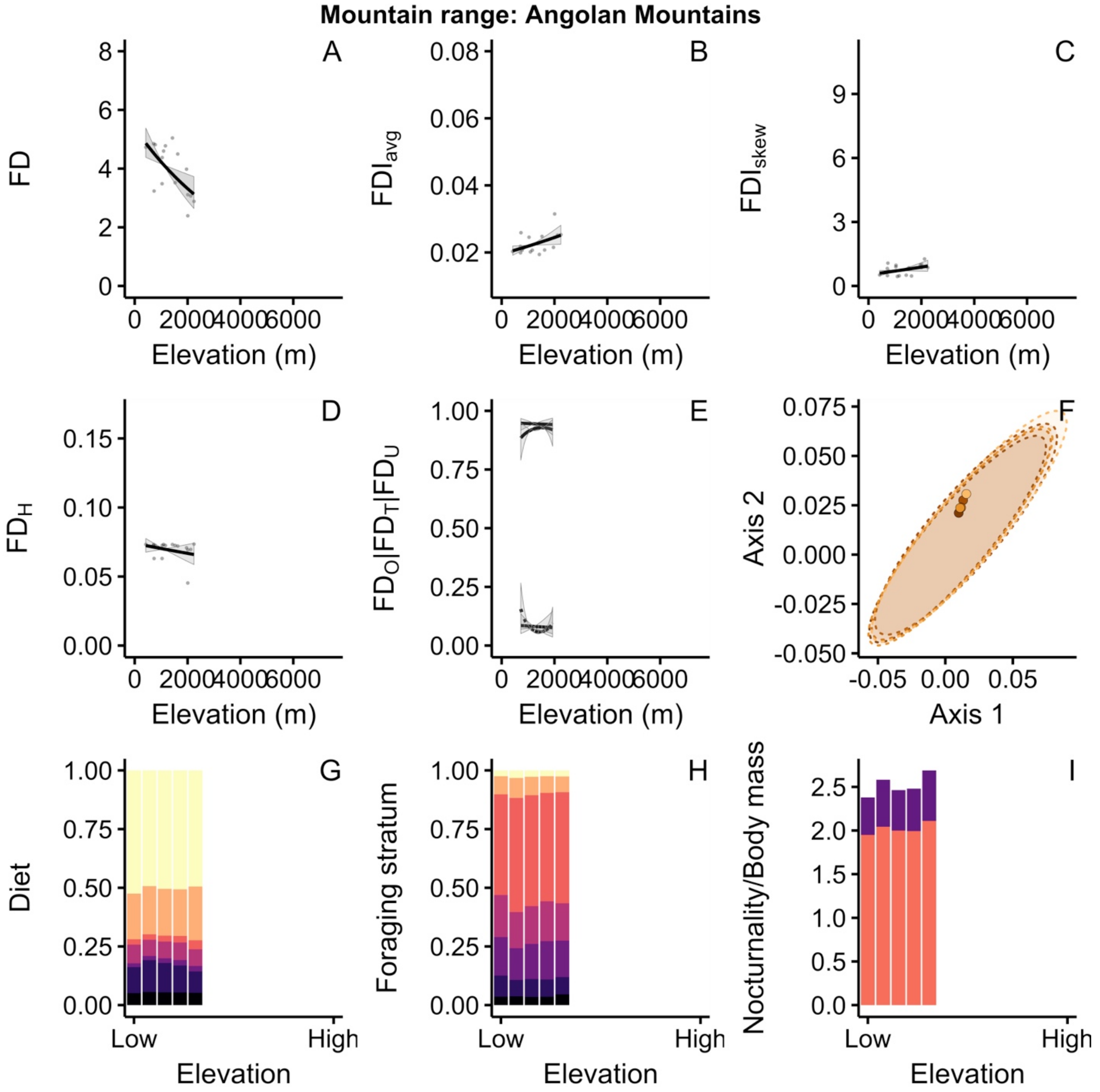

Mountain range: Angolan Mountains
A
B
C
D
E
F
G
H
I
FD
FDI_avg
FDI_skew
FD_H
FD_O|FD_T|FD_U
Axis 2
Axis 1
Diet
Foraging stratum
Nocturnality/Body mass
Elevation (m)
Elevation
Low
High
0
2000
4000
6000

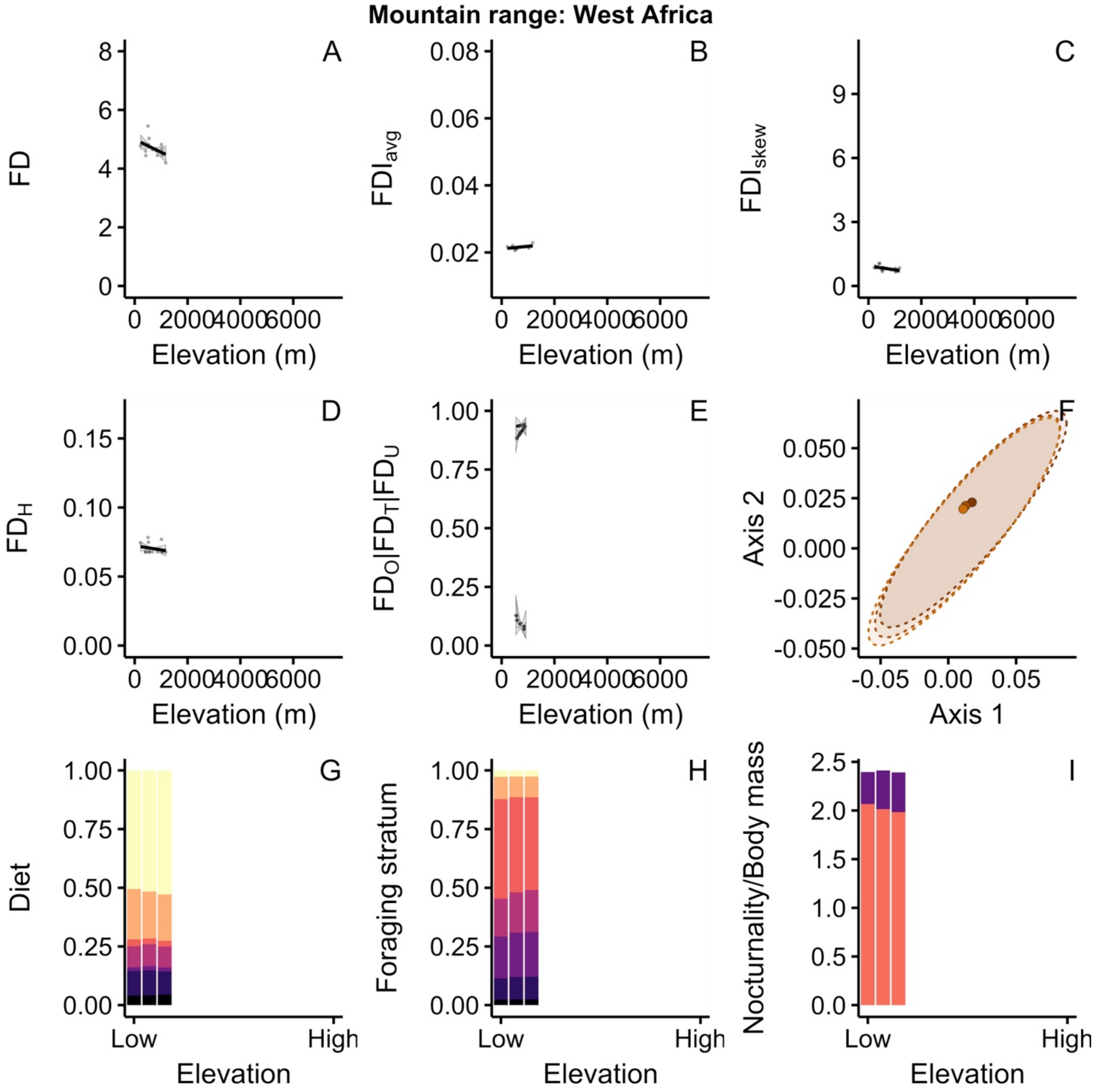

Mountain range: West Africa
A
B
C
D
E
F
G
H
I
FD
FDI$_{avg}$
FDI$_{skew}$
FD$_H$
FD$_O$|FD$_T$|FD$_U$
Axis 2
Axis 1
Diet
Foraging stratum
Nocturnality/Body mass
Elevation (m)
Elevation
Low
High

**Mountain range: Scandinavia**

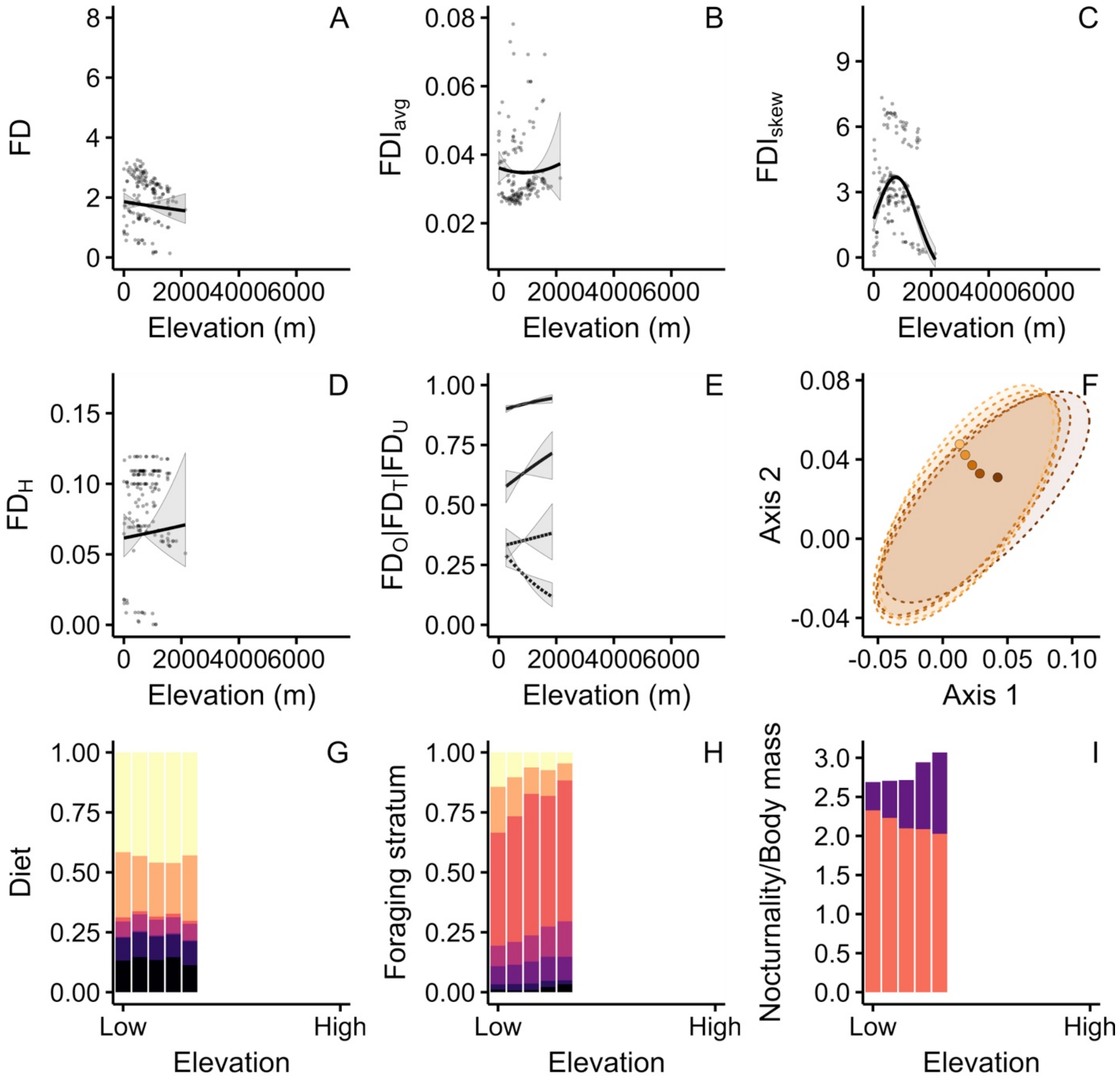

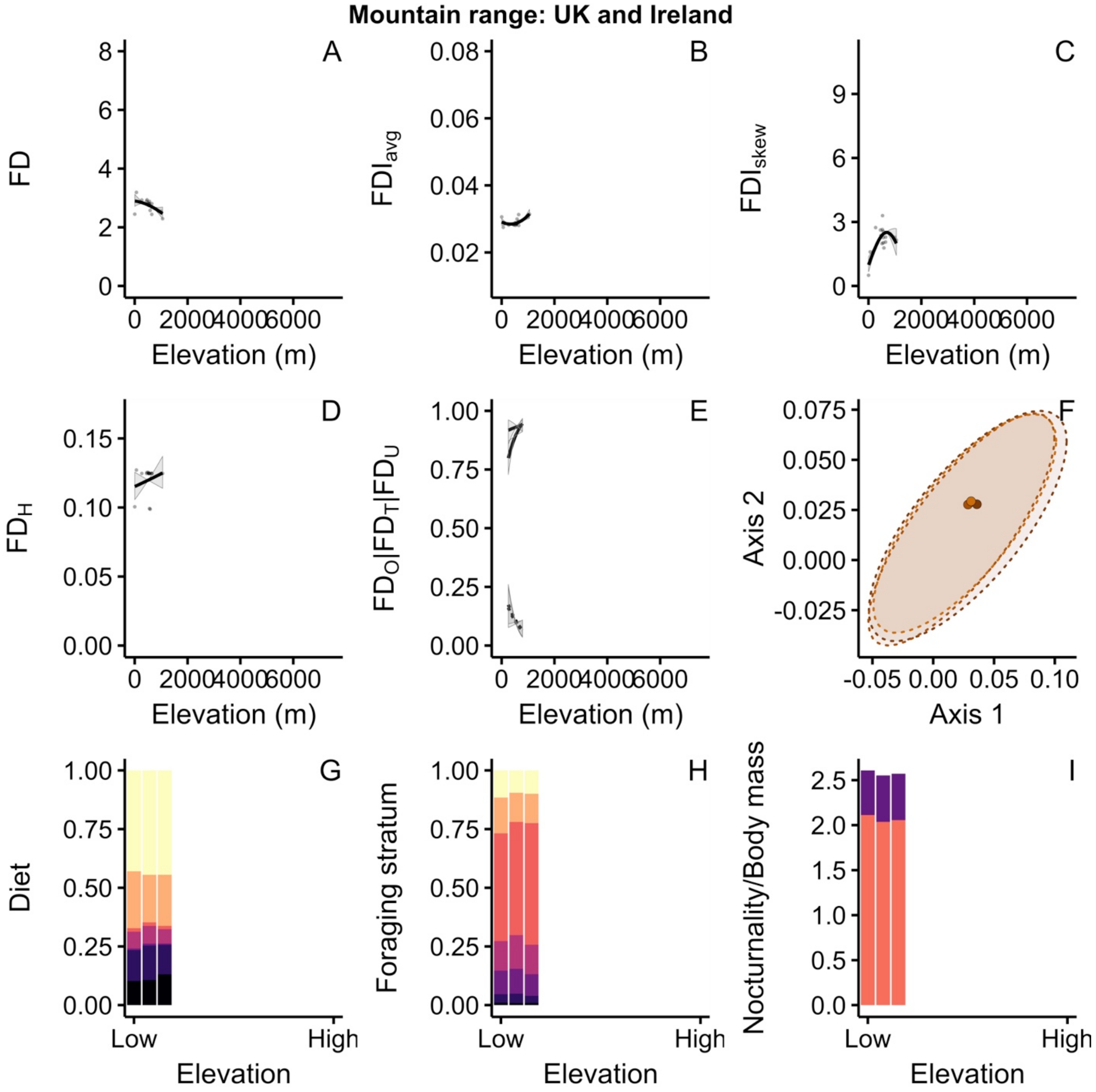
Mountain range: UK and Ireland
A
FD
Elevation (m)
B
FDI_avg
Elevation (m)
C
FDI_skew
Elevation (m)
D
FD_H
Elevation (m)
E
FD_O|FD_T|FD_U
Elevation (m)
F
Axis 2
Axis 1
G
Diet
Low
High
Elevation
H
Foraging stratum
Low
High
Elevation
I
Nocturnality/Body mass
Low
High
Elevation

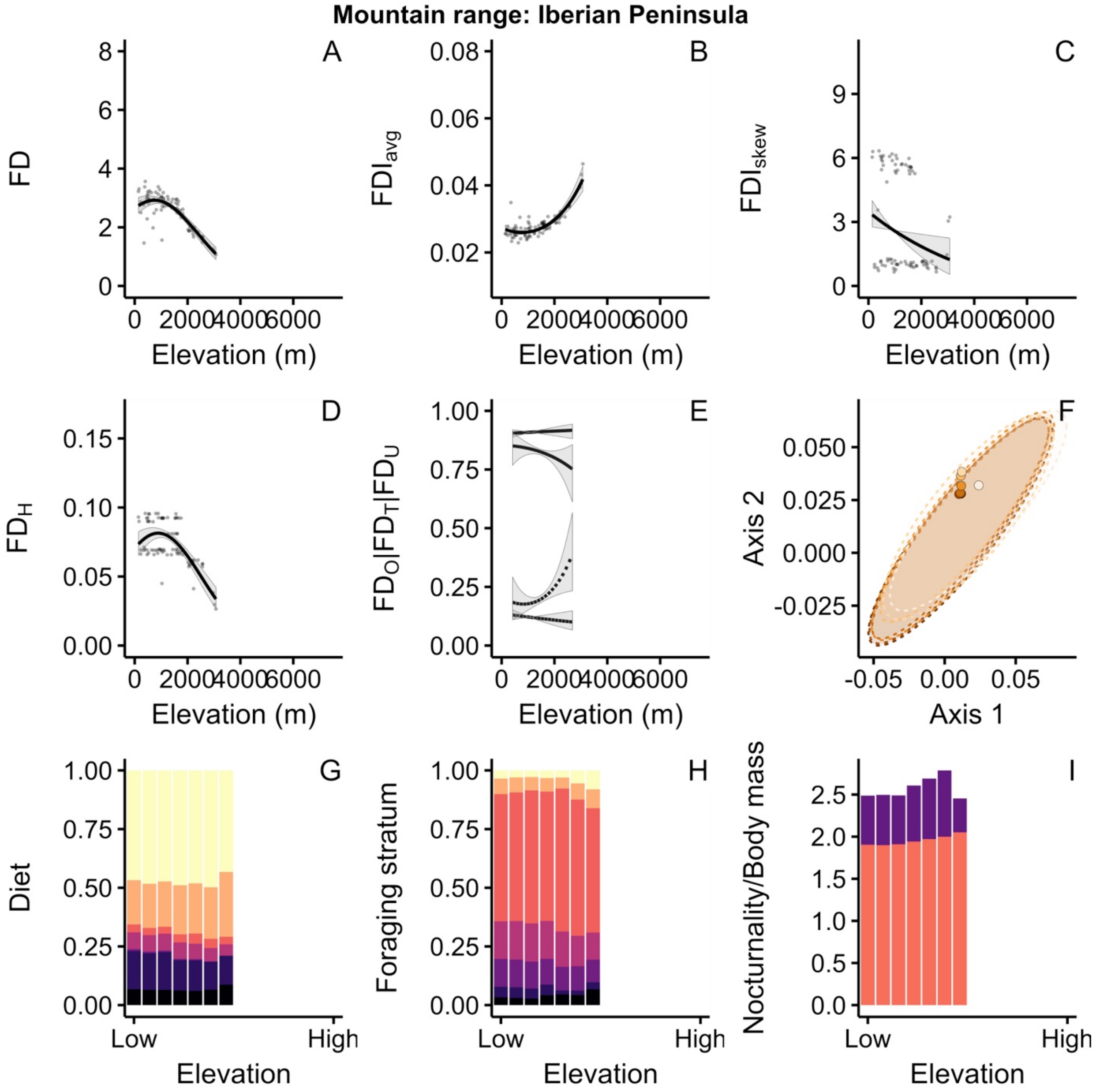

Mountain range: Iberian Peninsula
A
B
C
D
E
F
G
H
I
FD
FDI$_{avg}$
FDI$_{skew}$
FD$_H$
FD$_O$|FD$_T$|FD$_U$
Axis 2
Axis 1
Diet
Foraging stratum
Nocturnality/Body mass
Elevation (m)
Elevation
Low
High

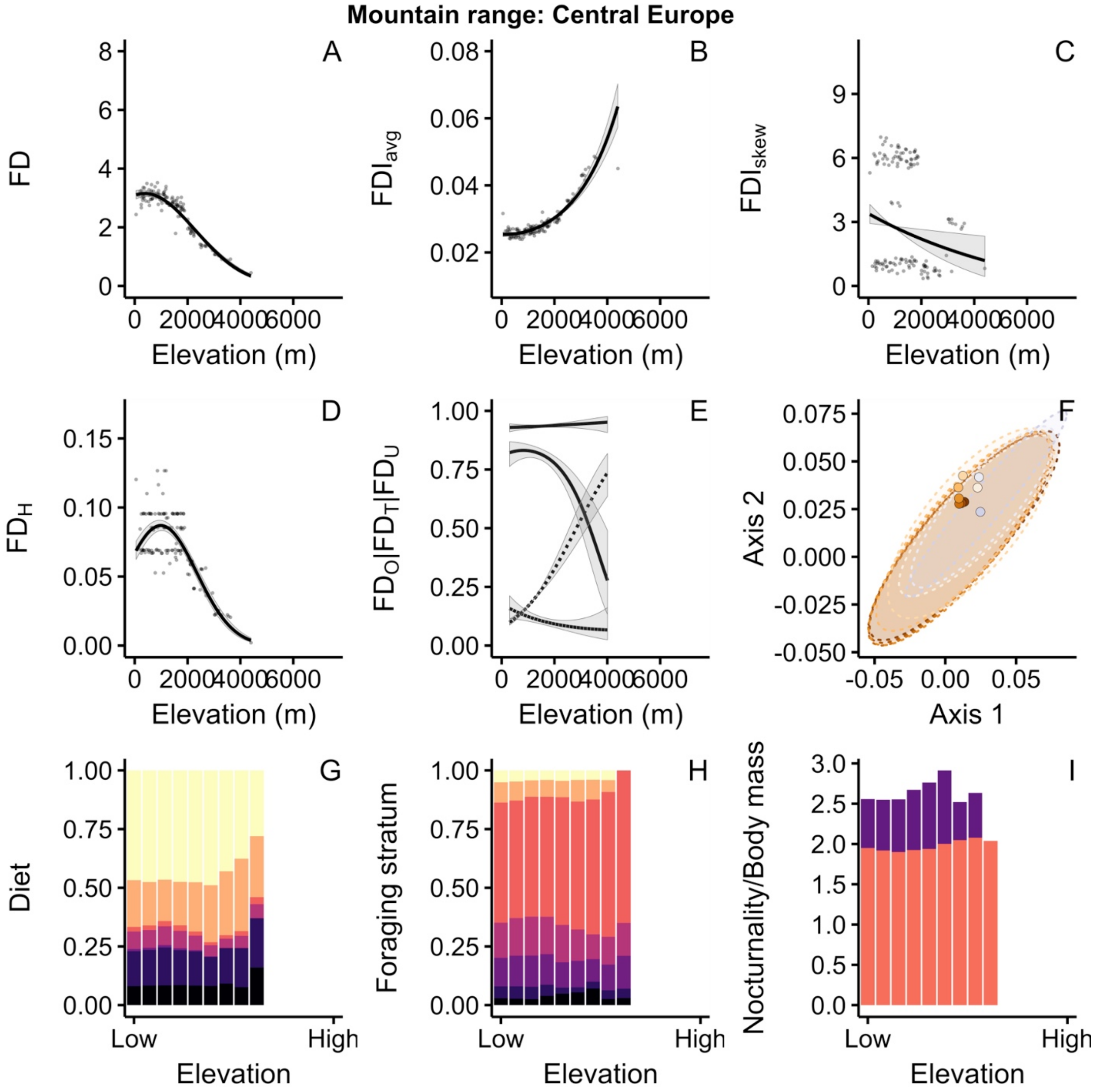
Mountain range: Central Europe
A
FD
Elevation (m)
B
FDI_avg
Elevation (m)
C
FDI_skew
Elevation (m)
D
FD_H
Elevation (m)
E
FD_O|FD_T|FD_U
Elevation (m)
F
Axis 2
Axis 1
G
Diet
Low
High
Elevation
H
Foraging stratum
Low
High
Elevation
I
Nocturnality/Body mass
Low
High
Elevation

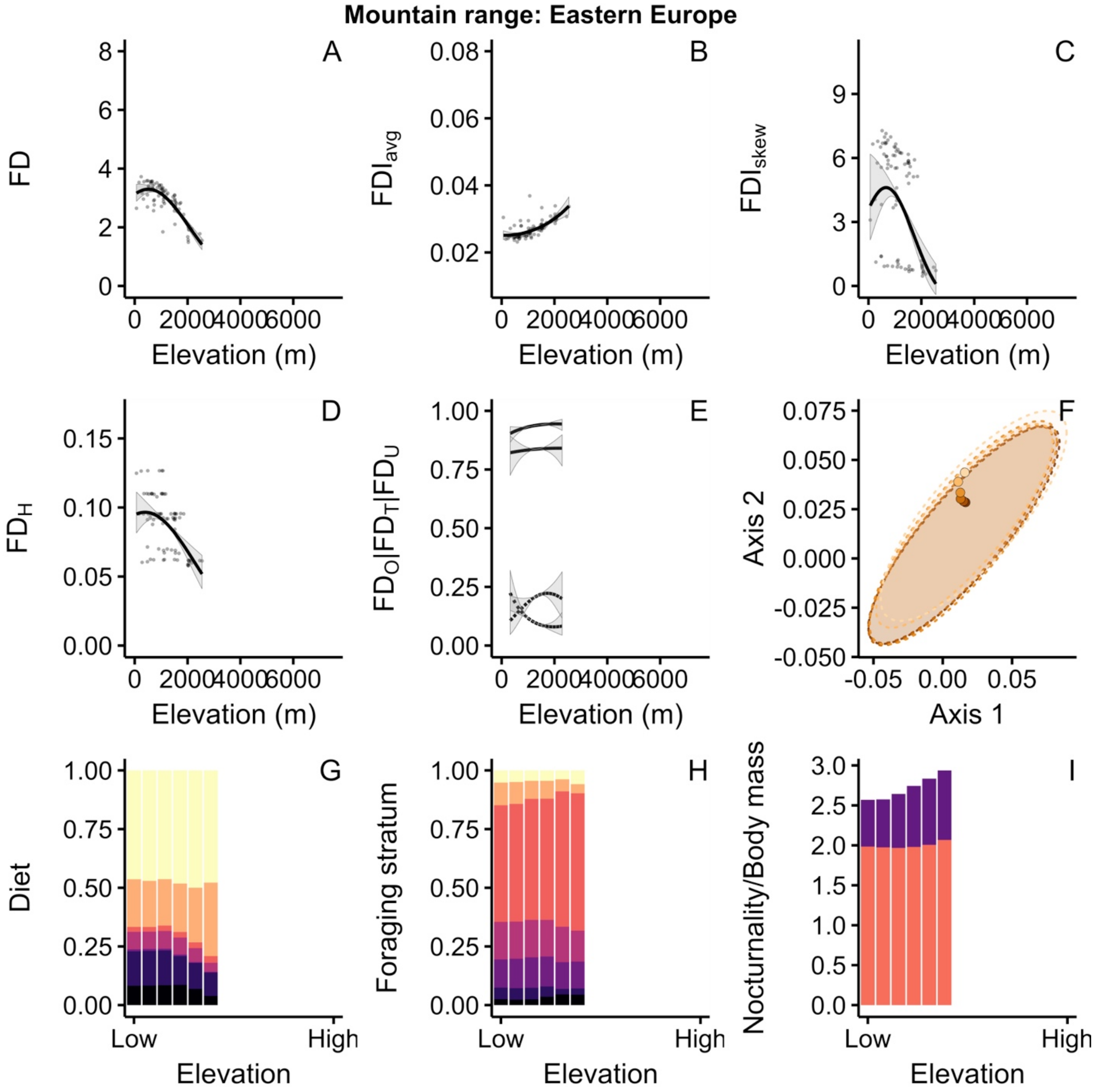

Mountain range: Eastern Europe
A
B
C
D
E
F
G
H
I
FD
FDI_avg
FDI_skew
FD_H
FD_O|FD_T|FD_U
Axis 2
Axis 1
Diet
Foraging stratum
Nocturnality/Body mass
Elevation (m)
Elevation
Low
High

**Mountain range: Italian Mountains**

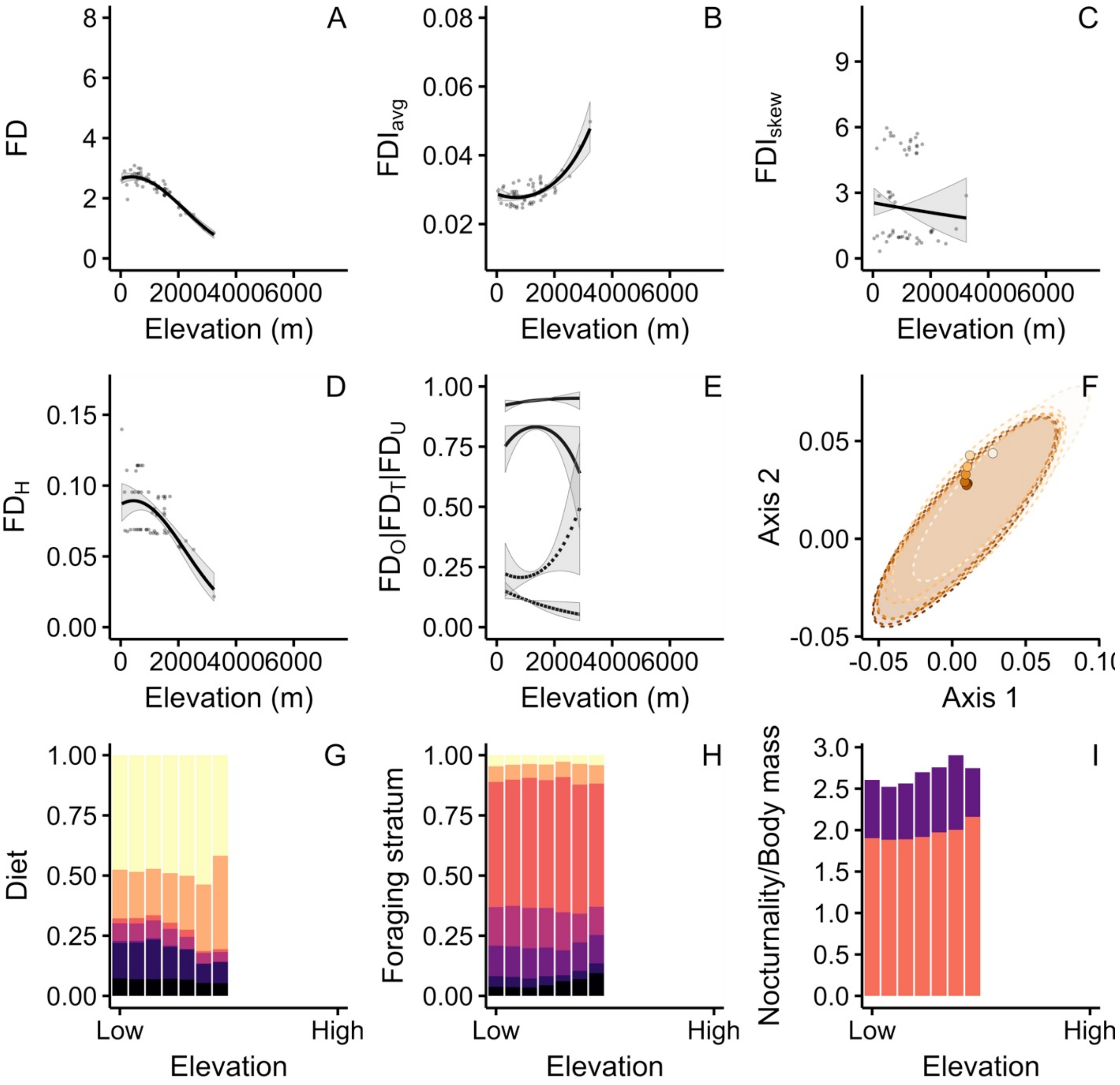

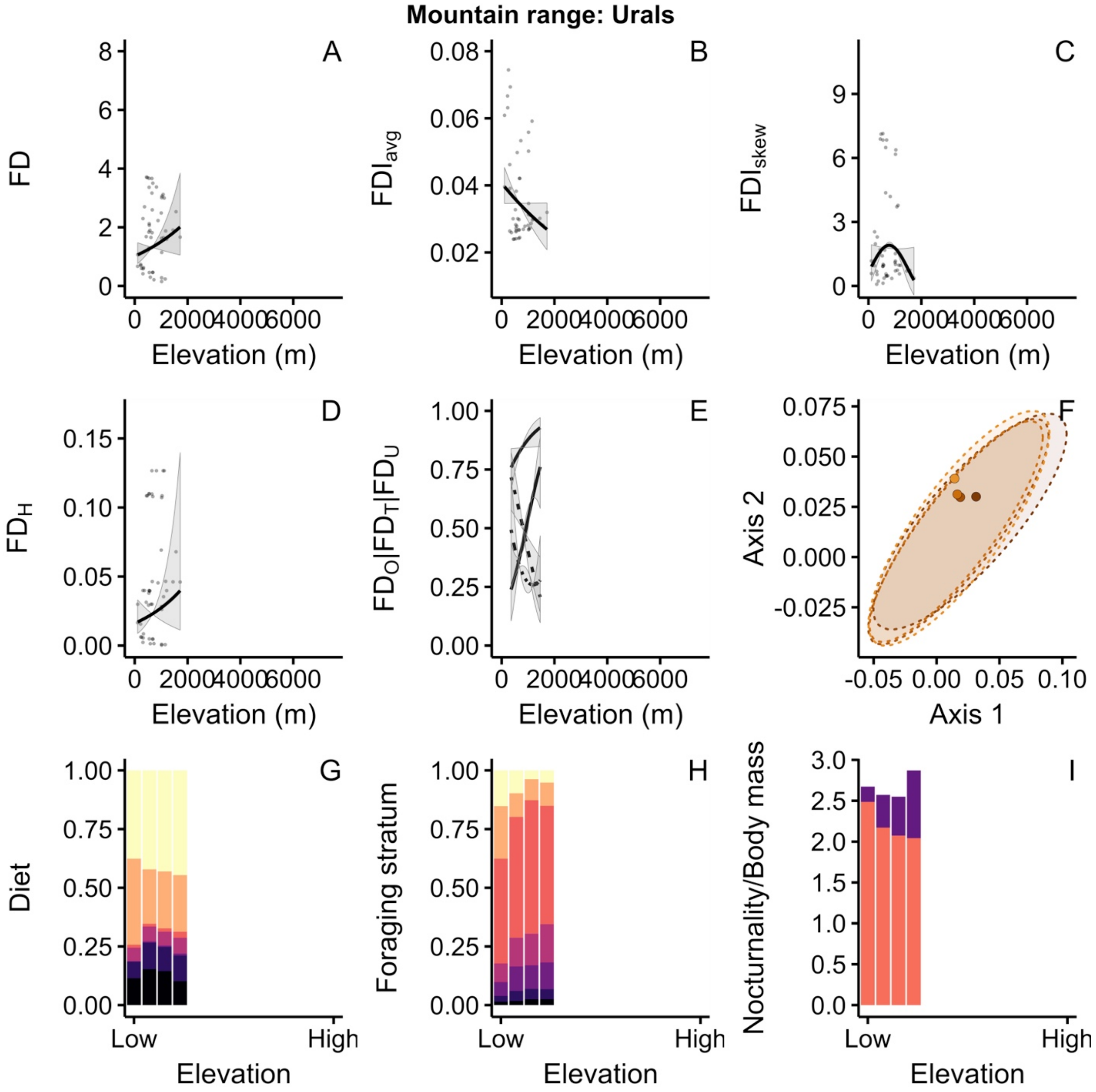

Mountain range: Urals
A
B
C
D
E
F
G
H
I
FD
FDI_avg
FDI_skew
FD_H
FD_O|FD_T|FD_U
Axis 2
Axis 1
Diet
Foraging stratum
Nocturnality/Body mass
Elevation (m)
Elevation
Low
High

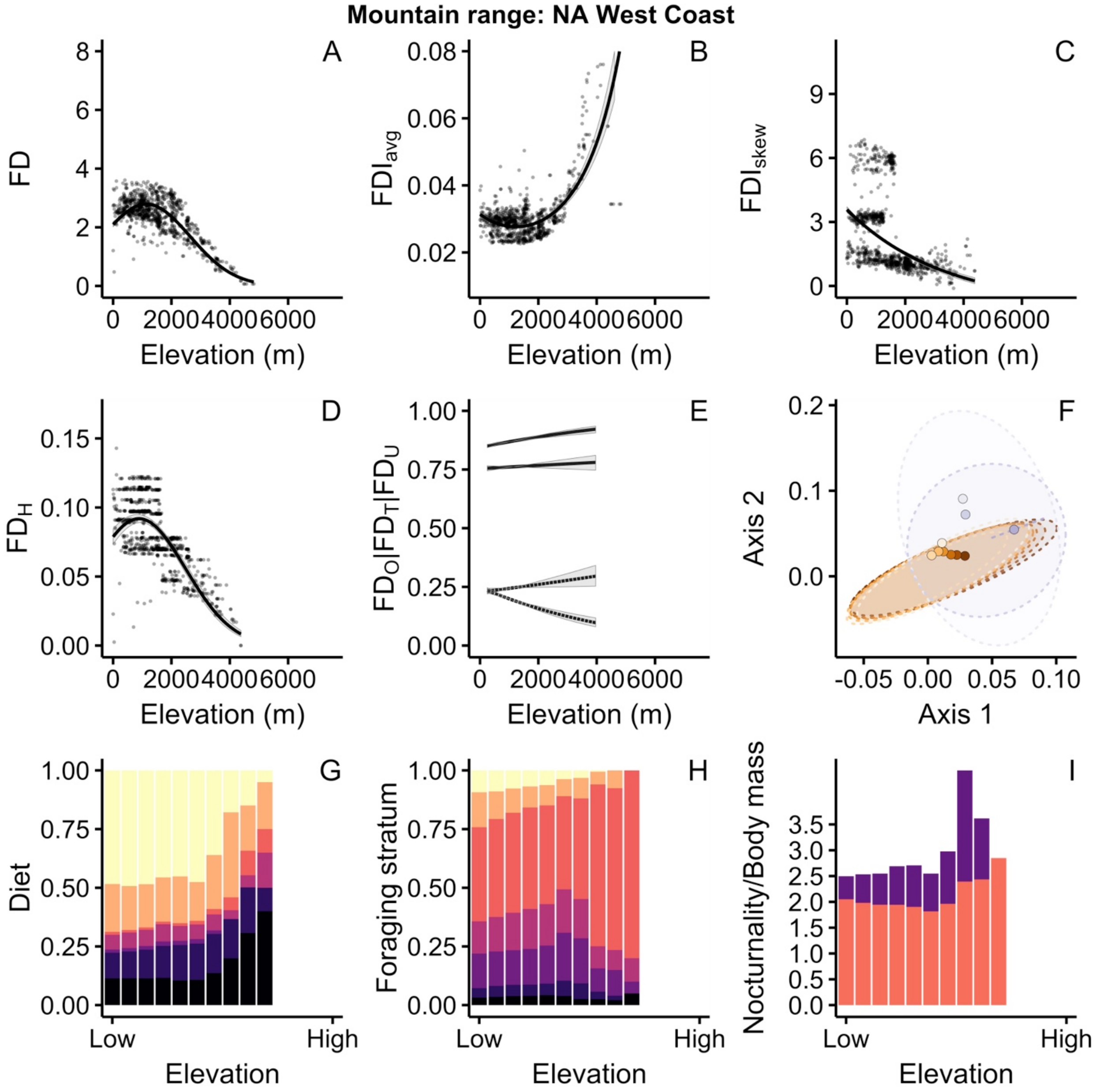

Mountain range: NA West Coast
A
B
C
D
E
F
G
H
I
FD
FDI_avg
FDI_skew
FD_H
FD_O|FD_T|FD_U
Axis 1
Axis 2
Diet
Foraging stratum
Nocturnality/Body mass
Elevation (m)
Elevation
Low
High

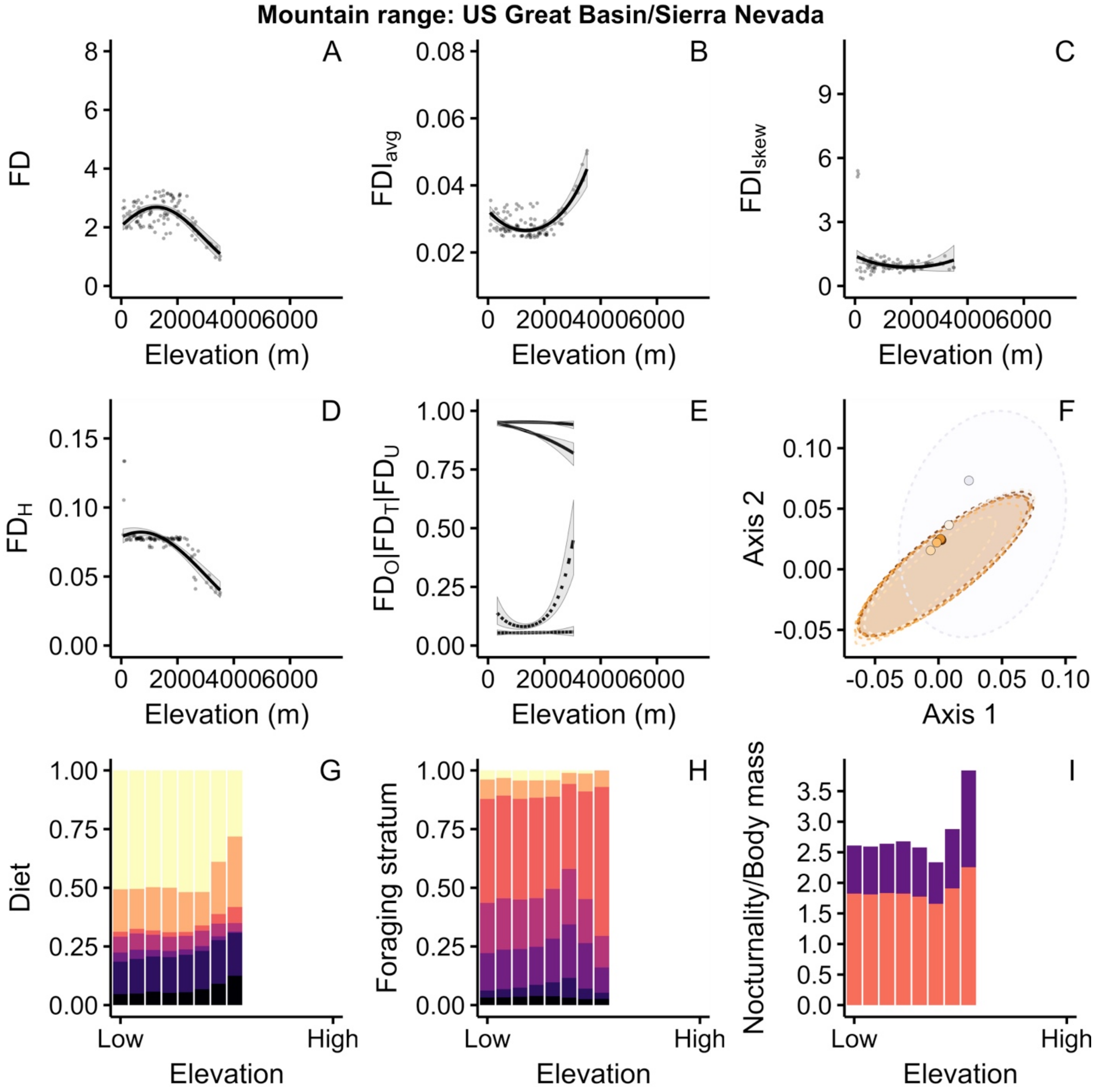

Mountain range: US Great Basin/Sierra Nevada
A
B
C
D
E
F
G
H
I
FD
$FDI_{avg}$
$FDI_{skew}$
$FD_H$
$FD_O|FD_T|FD_U$
Axis 2
Axis 1
Diet
Foraging stratum
Nocturnality/Body mass
Elevation (m)
Elevation
Low
High

**Mountain range: NA East Coast**

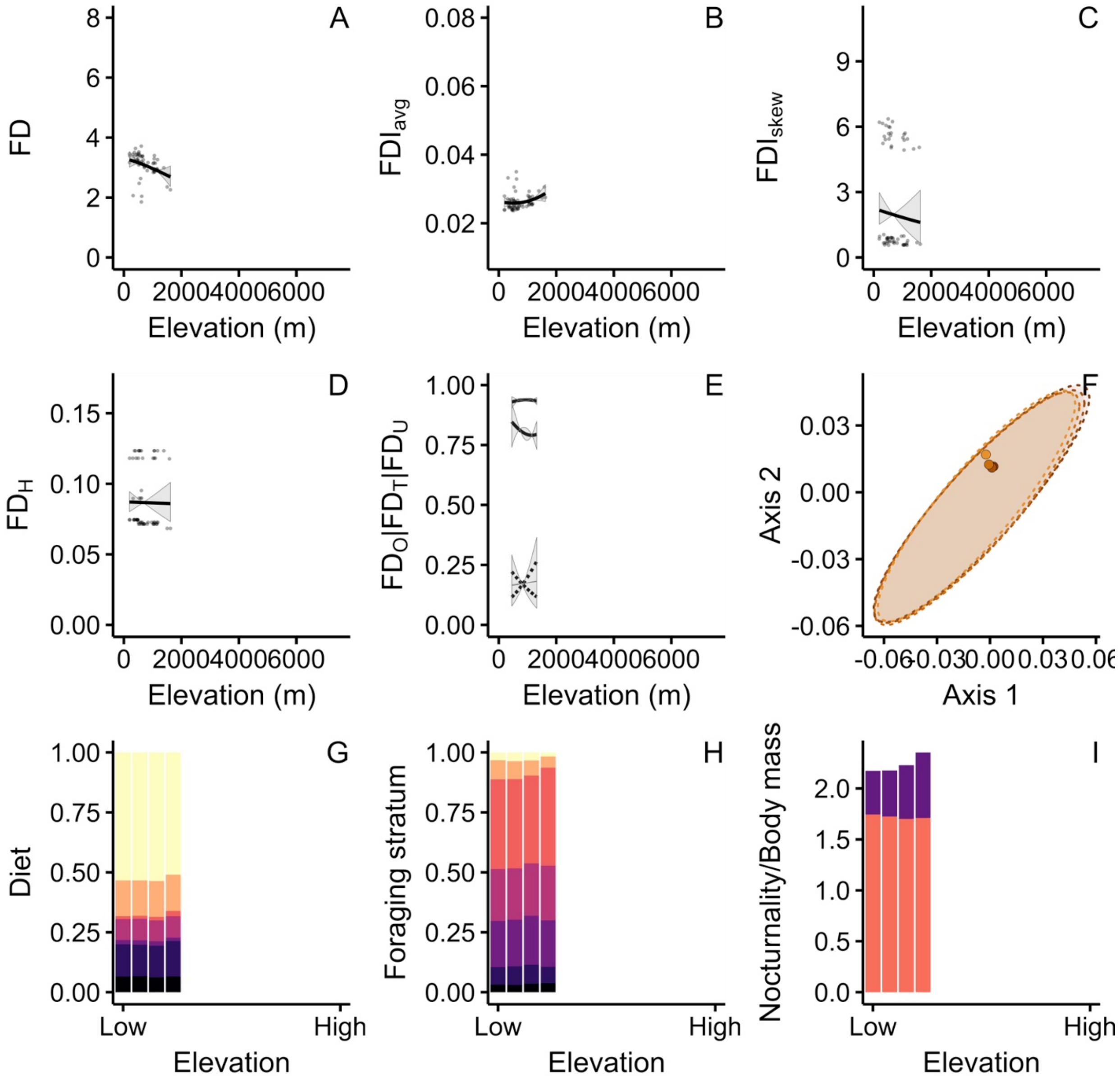

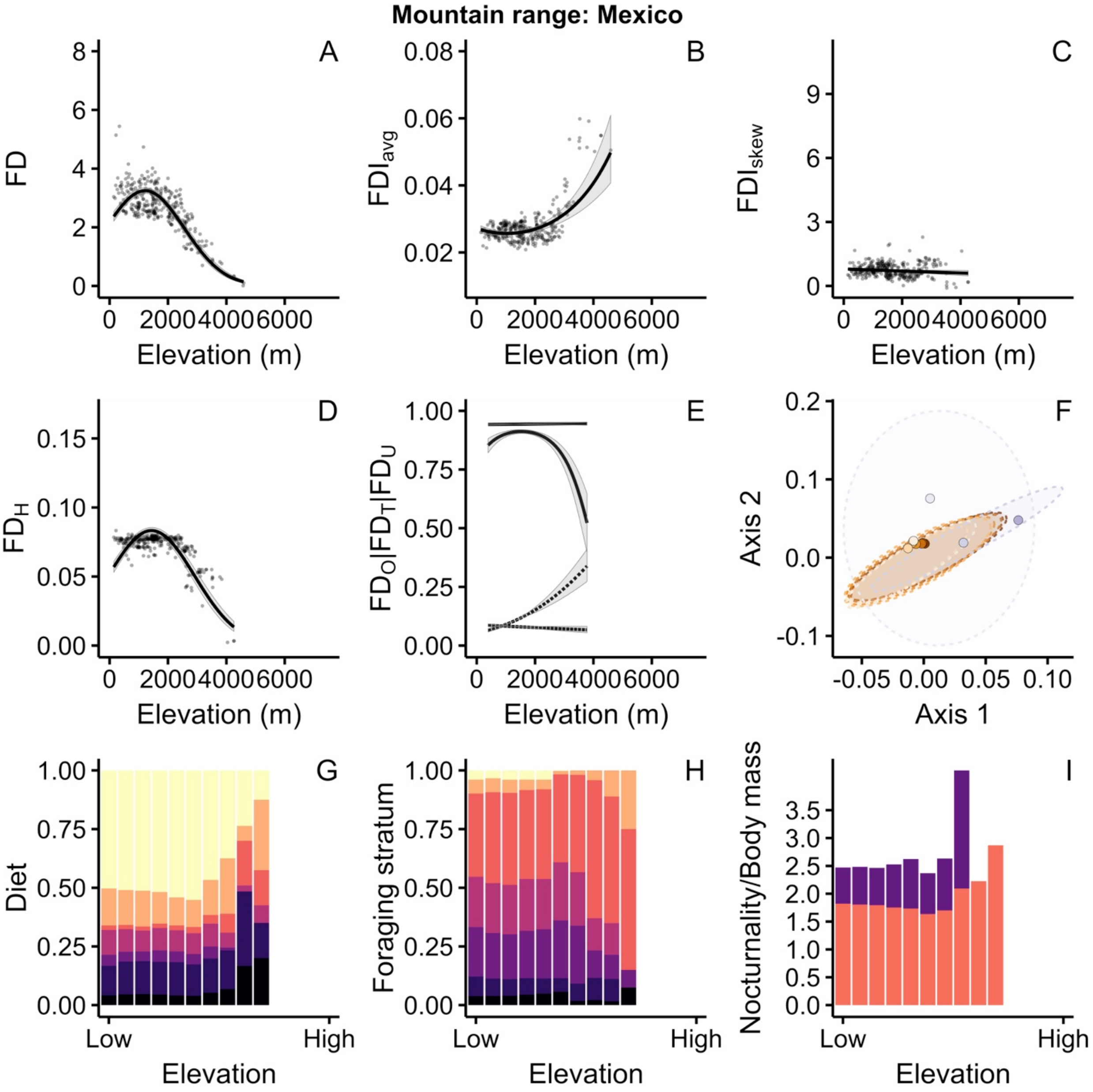
Mountain range: Mexico
A
B
C
D
E
F
G
H
I
FD
FDI_avg
FDI_skew
FD_H
FD_O|FD_T|FD_U
Axis 2
Axis 1
Diet
Foraging stratum
Nocturnality/Body mass
Elevation (m)
Elevation
Low
High

**Mountain range: Sierra Madre de Chiapas**

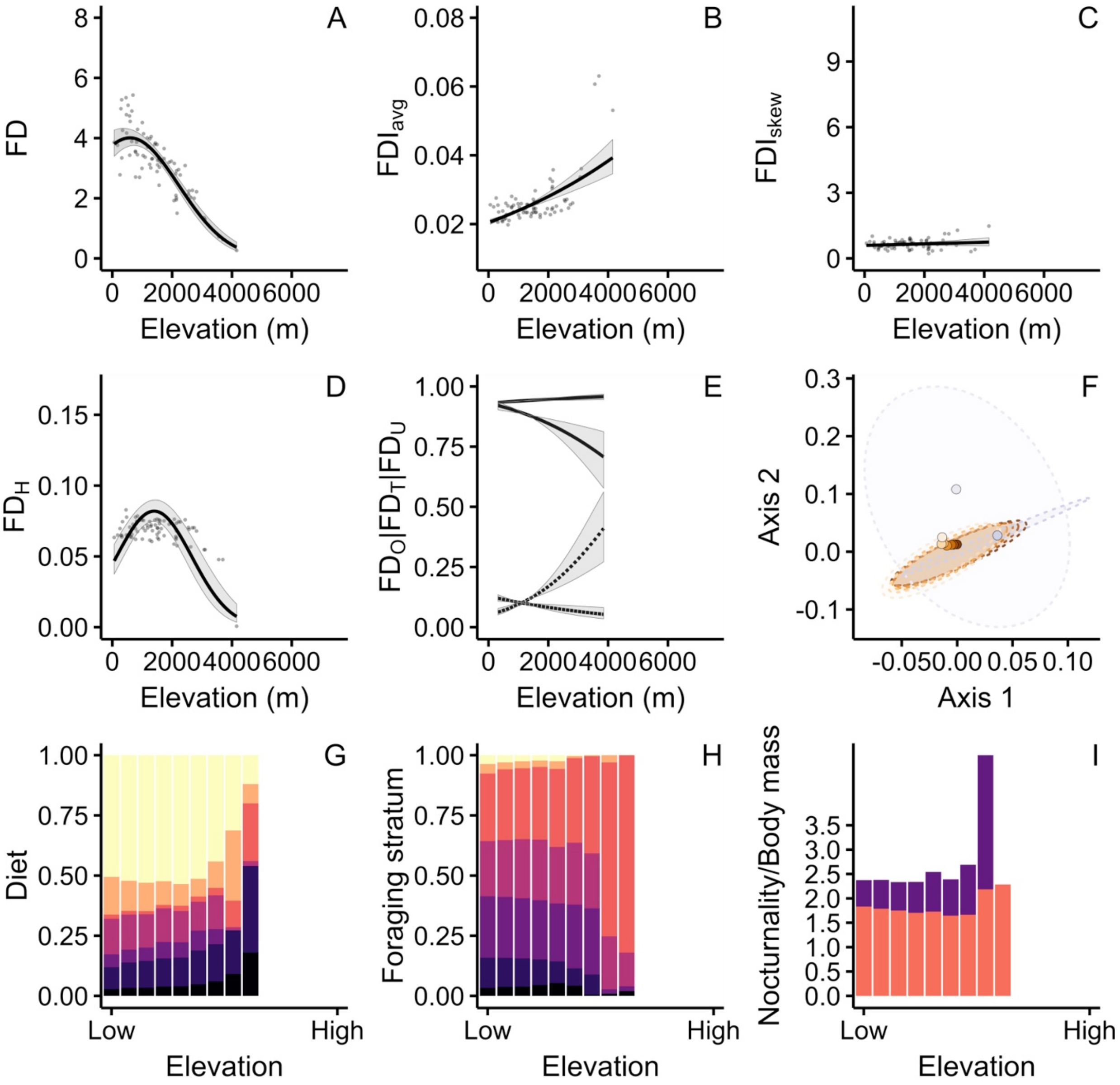

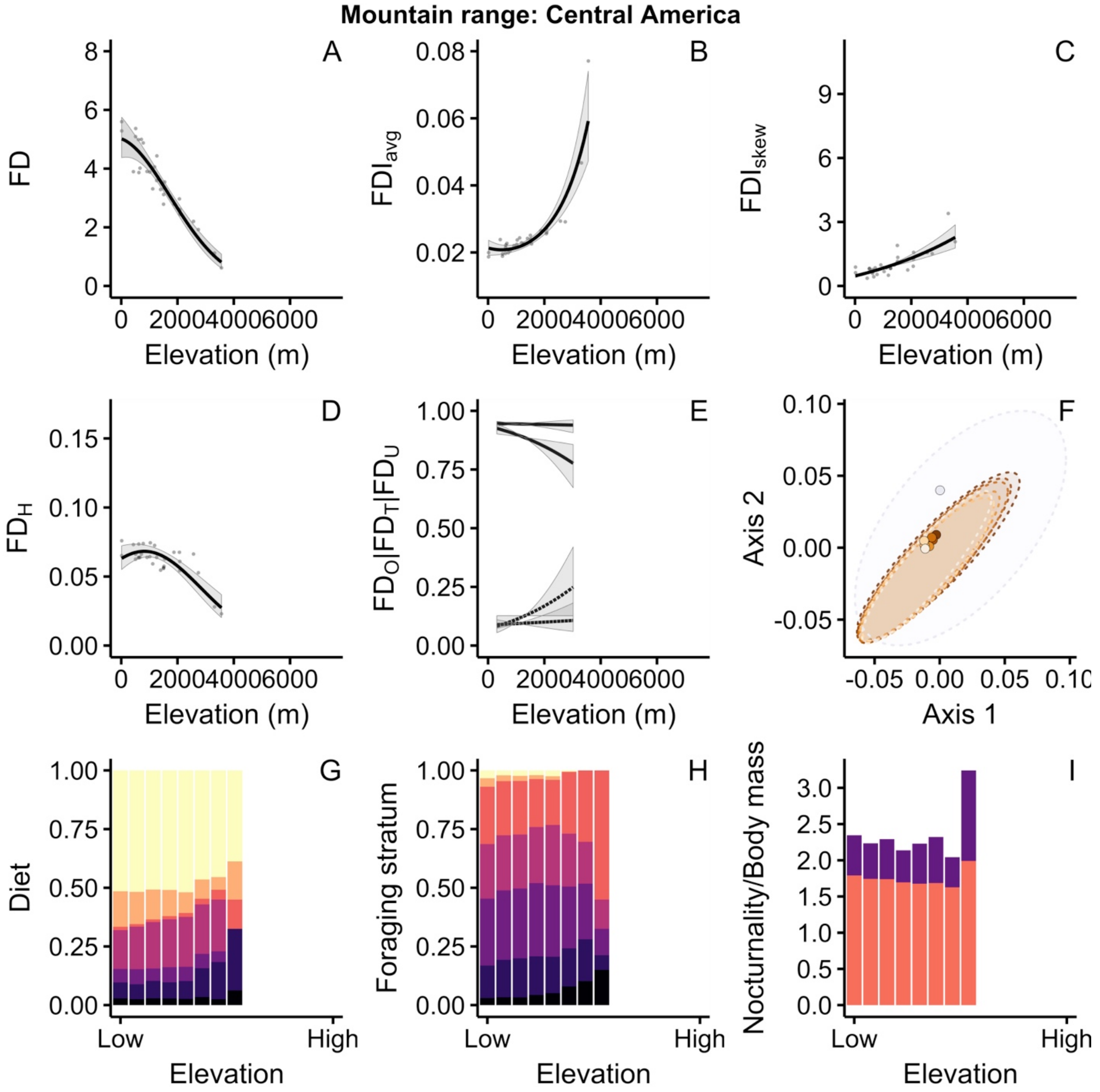

Mountain range: Central America
A
B
C
D
E
F
G
H
I
FD
FDI_avg
FDI_skew
FD_H
FD_O|FD_T|FD_U
Axis 2
Axis 1
Diet
Foraging stratum
Nocturnality/Body mass
Elevation (m)
Elevation
Low
High

**Mountain range: Caribbean Islands**

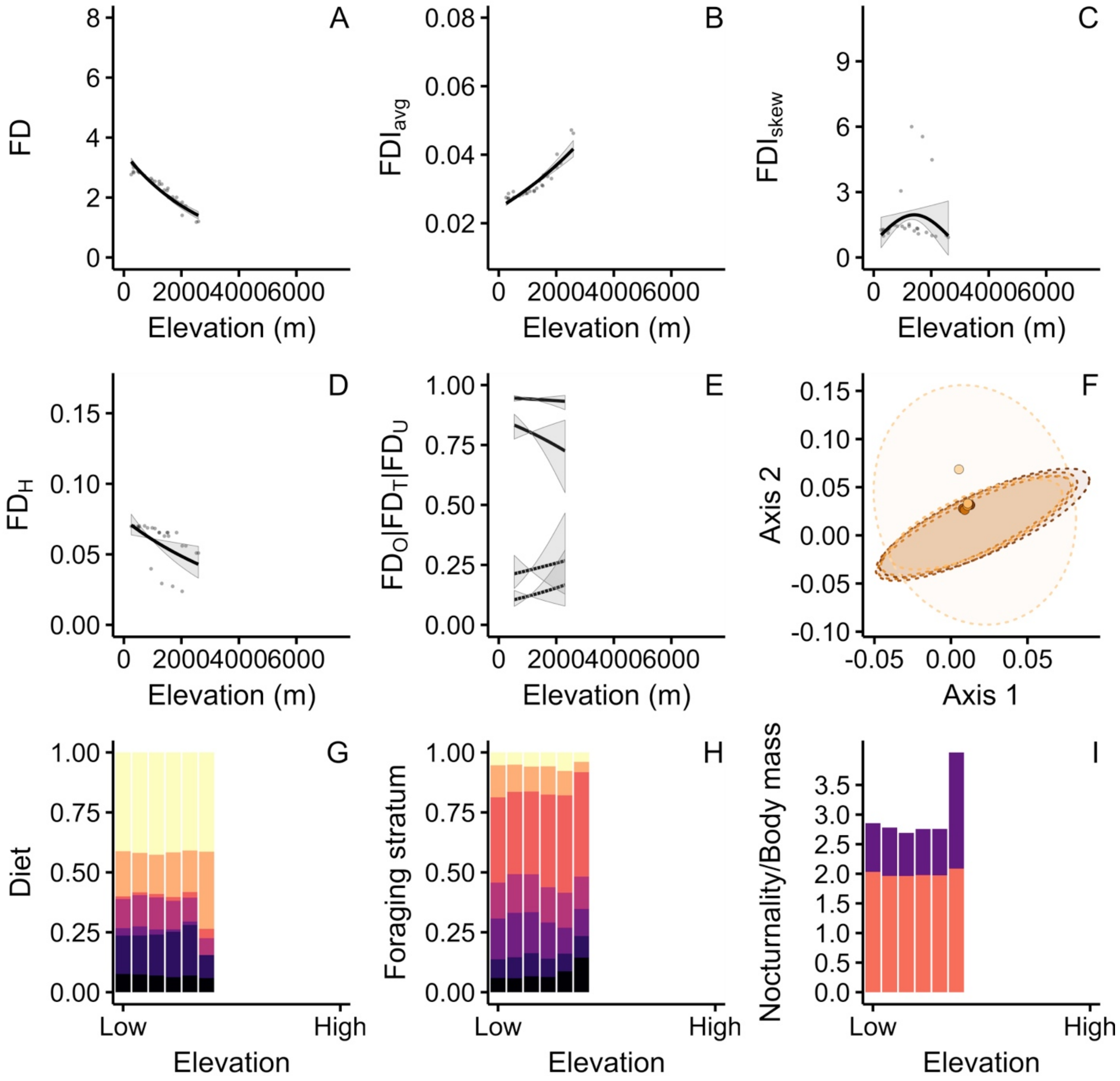

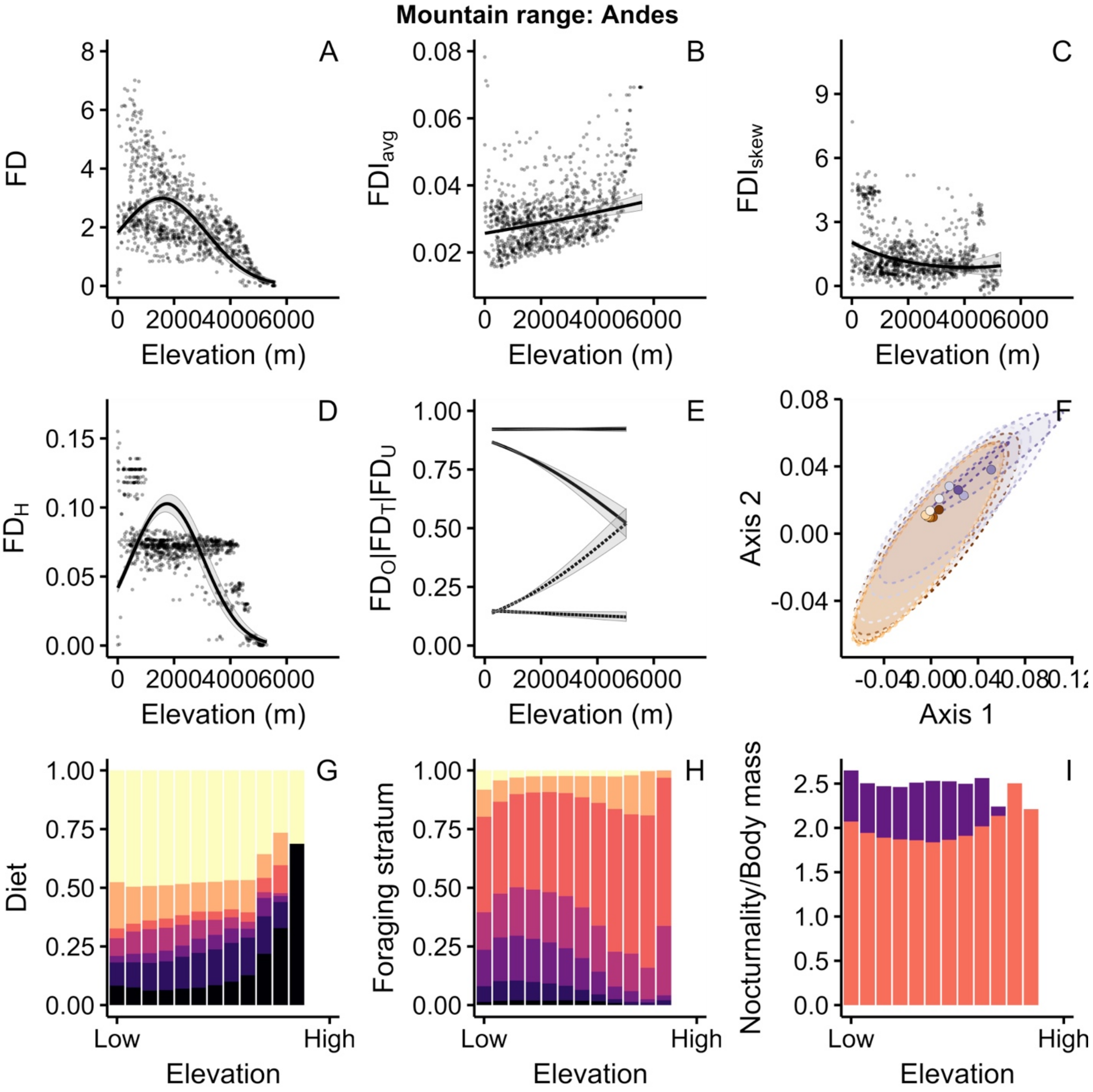

Mountain range: Andes
A
B
C
D
E
F
G
H
I
FD
FDI_avg
FDI_skew
FD_H
FD_O|FD_T|FD_U
Axis 2
Axis 1
Elevation (m)
Diet
Foraging stratum
Nocturnality/Body mass
Elevation
Low
High

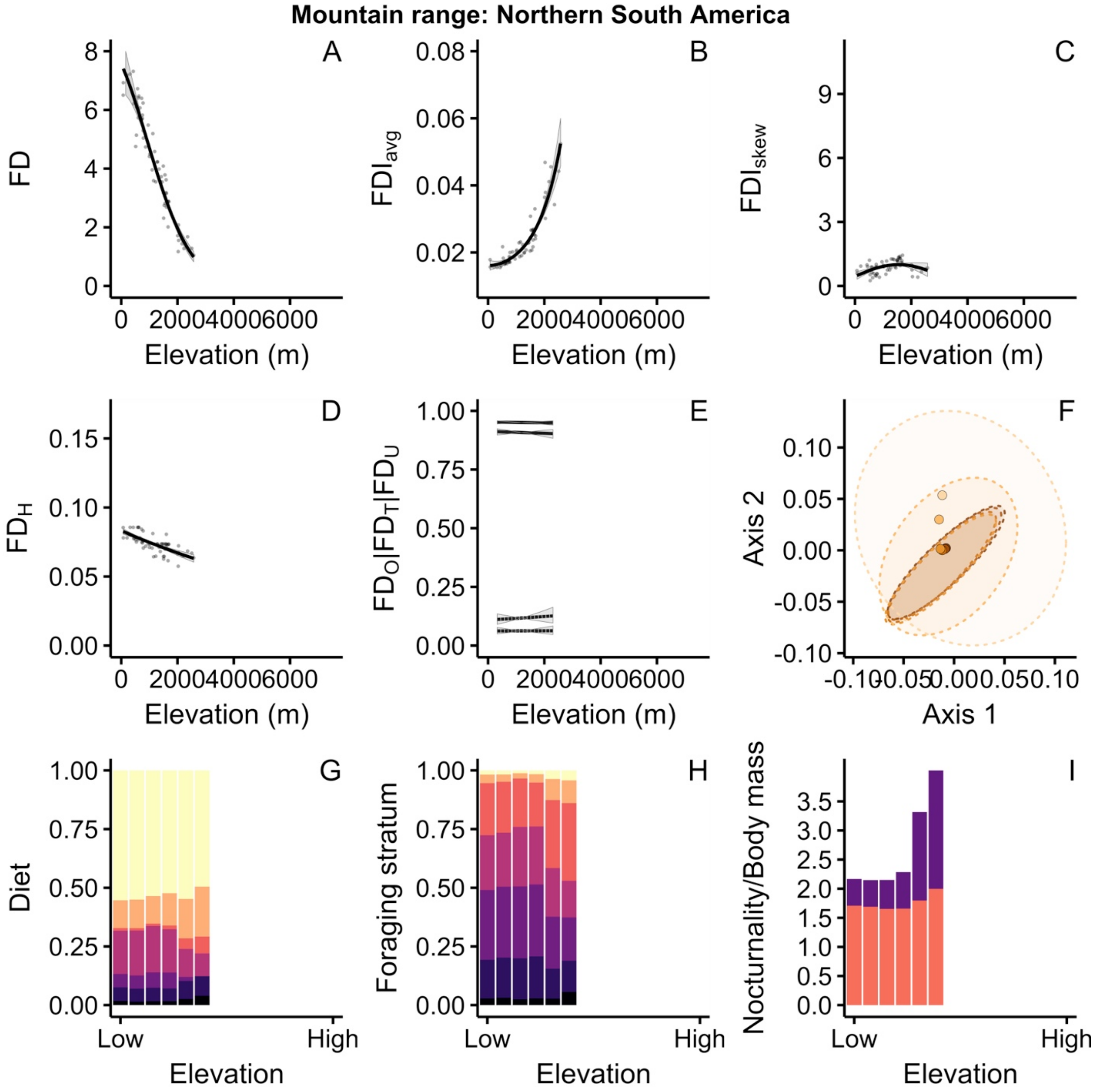

Mountain range: Northern South America
A
B
C
D
E
F
G
H
I
FD
$FDI_{avg}$
$FDI_{skew}$
$FD_H$
$FD_O|FD_T|FD_U$
Axis 2
Axis 1
Diet
Foraging stratum
Nocturnality/Body mass
Elevation (m)
Elevation
Low
High

**Mountain range: Eastern South America**

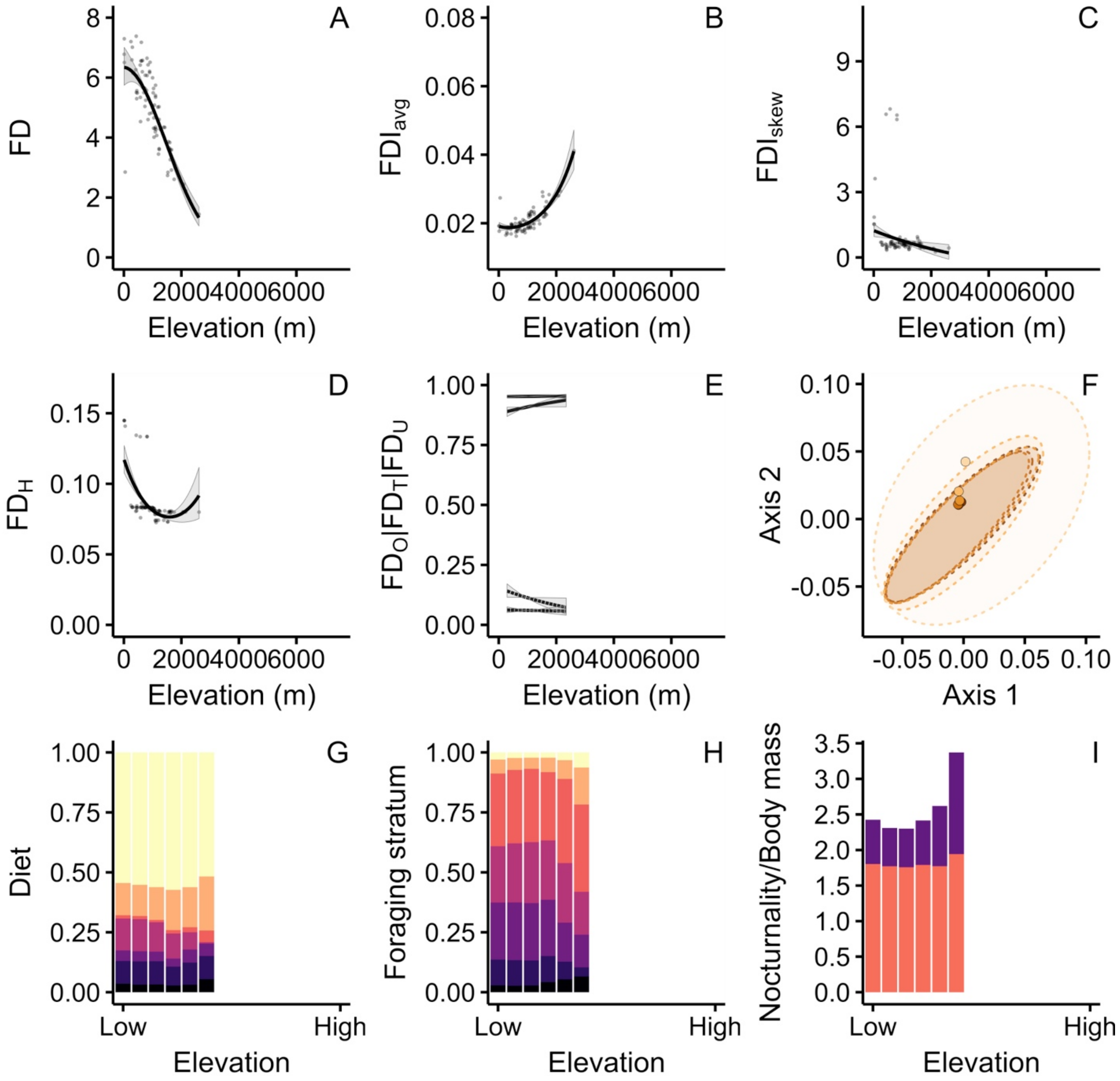

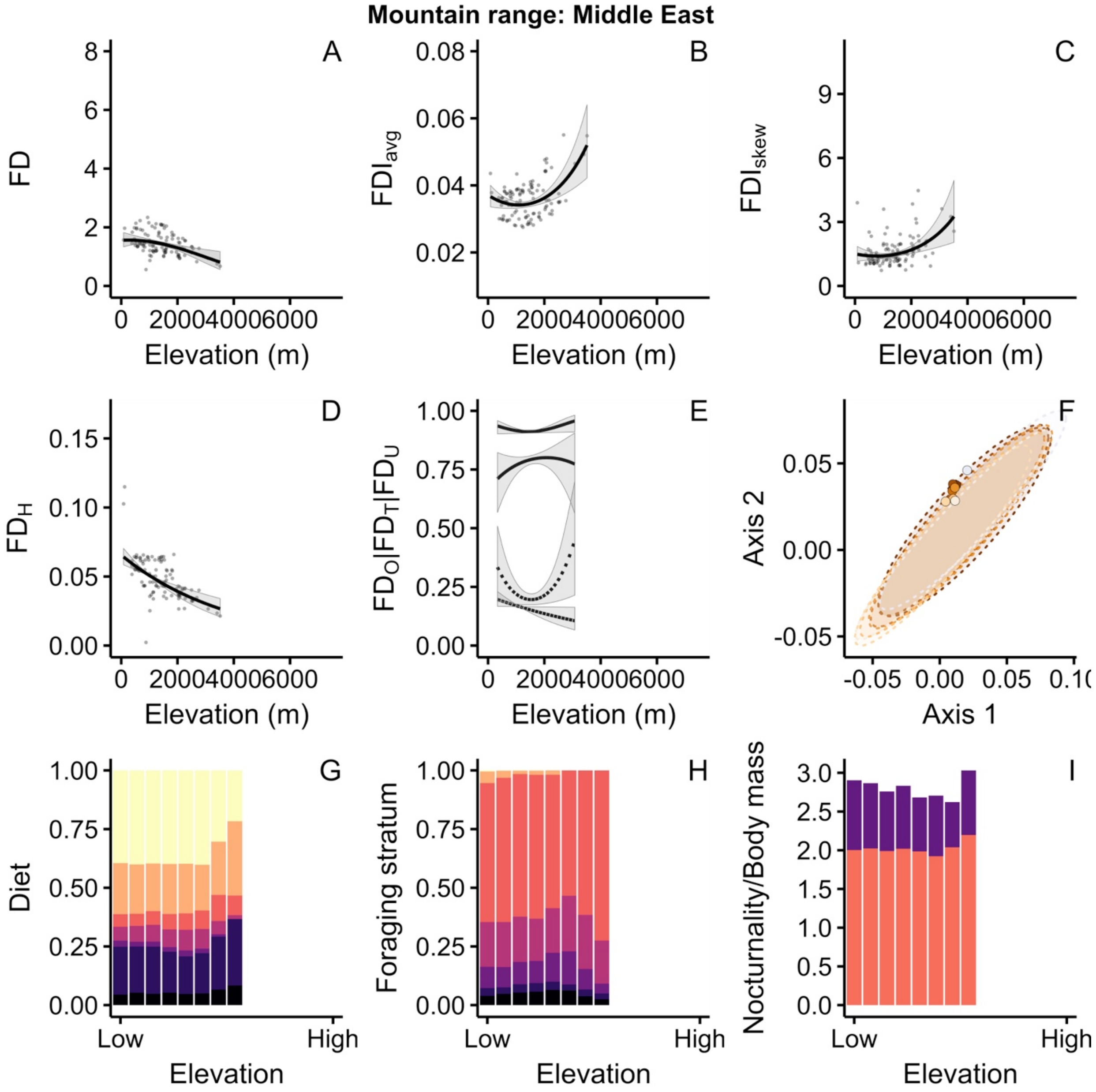

Mountain range: Middle East
A
B
C
D
E
F
G
H
I
FD
FDI$_{avg}$
FDI$_{skew}$
FD$_H$
FD$_O$|FD$_T$|FD$_U$
Axis 2
Axis 1
Diet
Foraging stratum
Nocturnality/Body mass
Elevation (m)
Elevation
Low
High

**Mountain range: Hindukush-Himalaya**

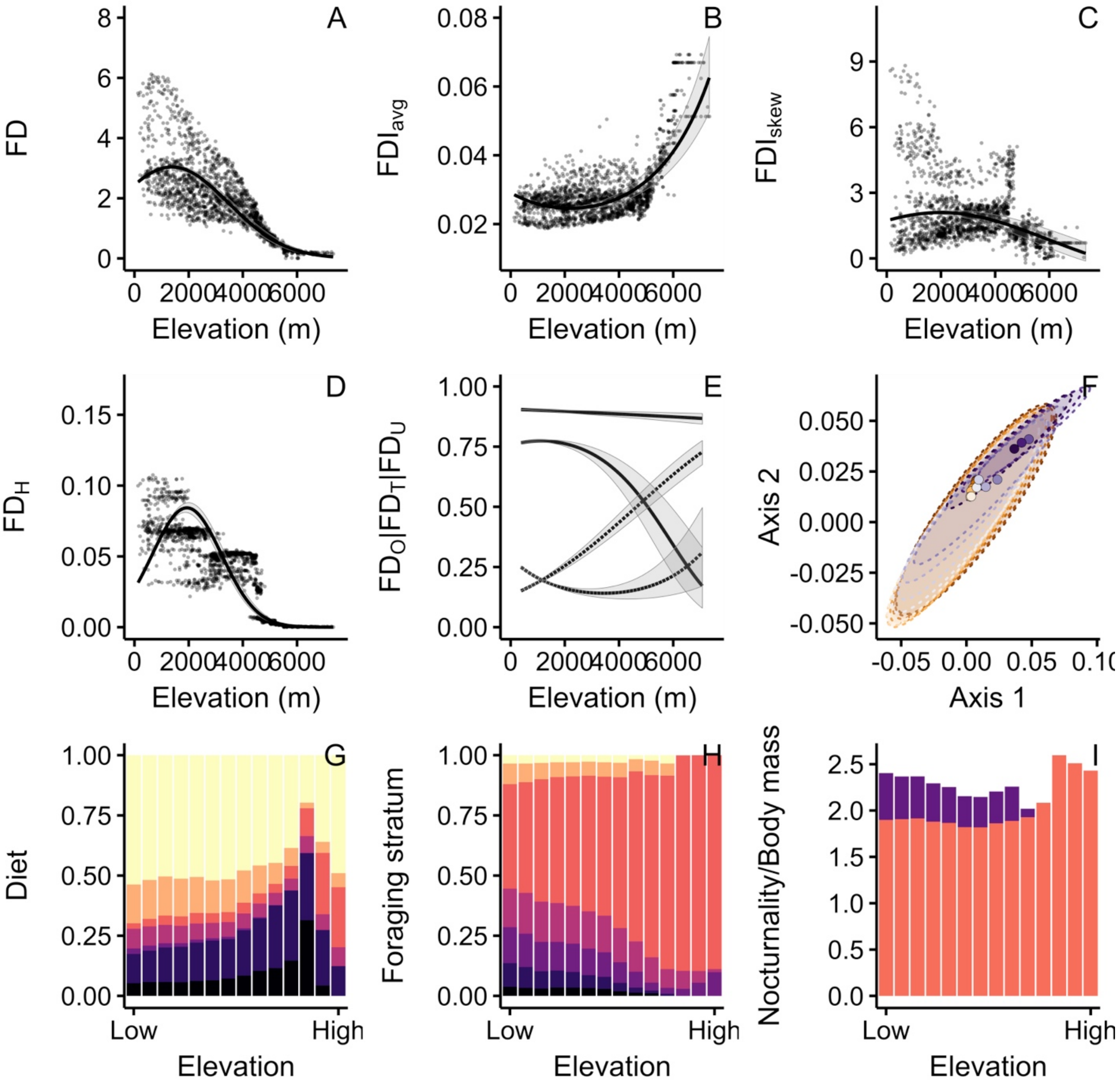

**Mountain range: India**

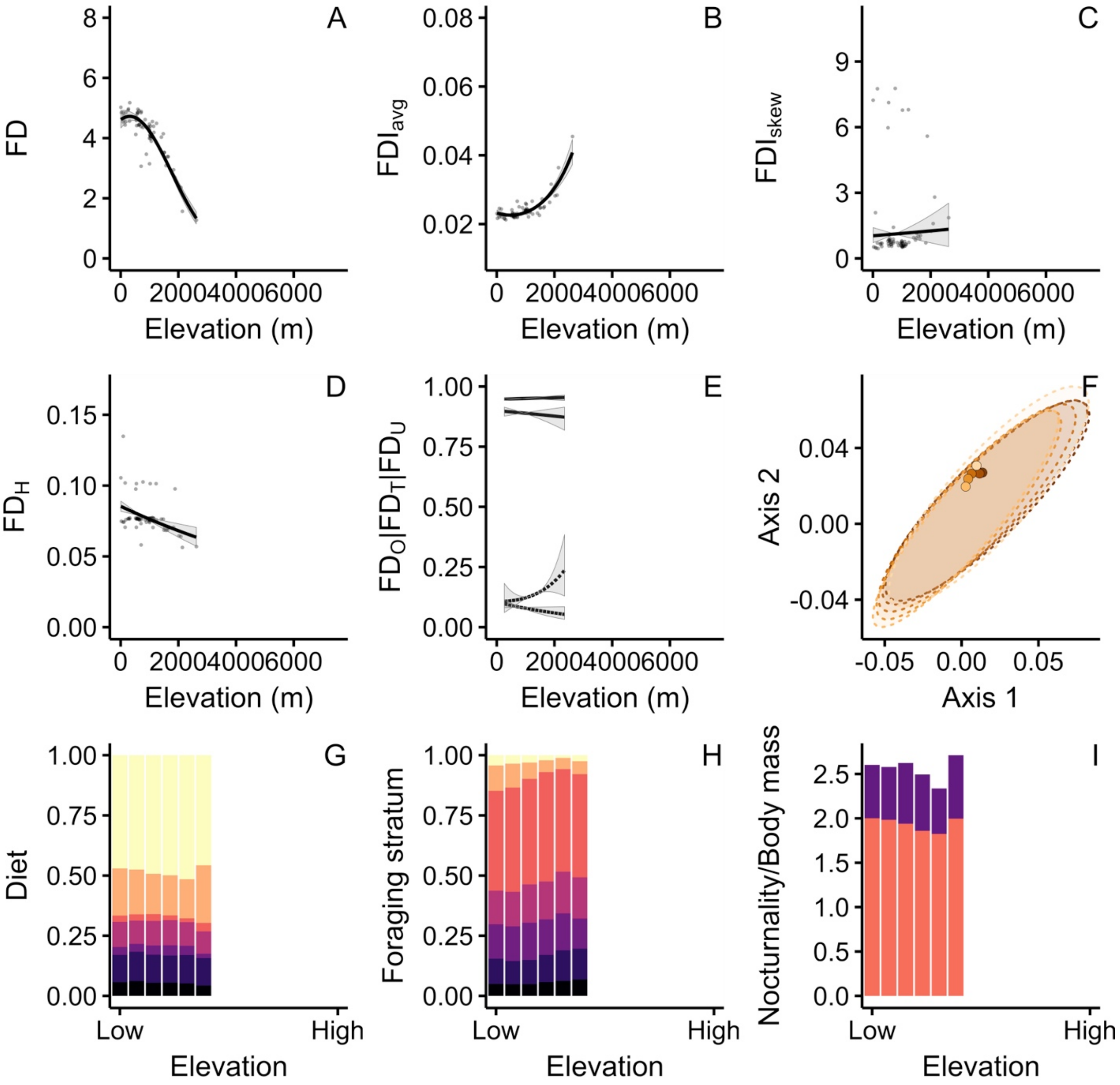

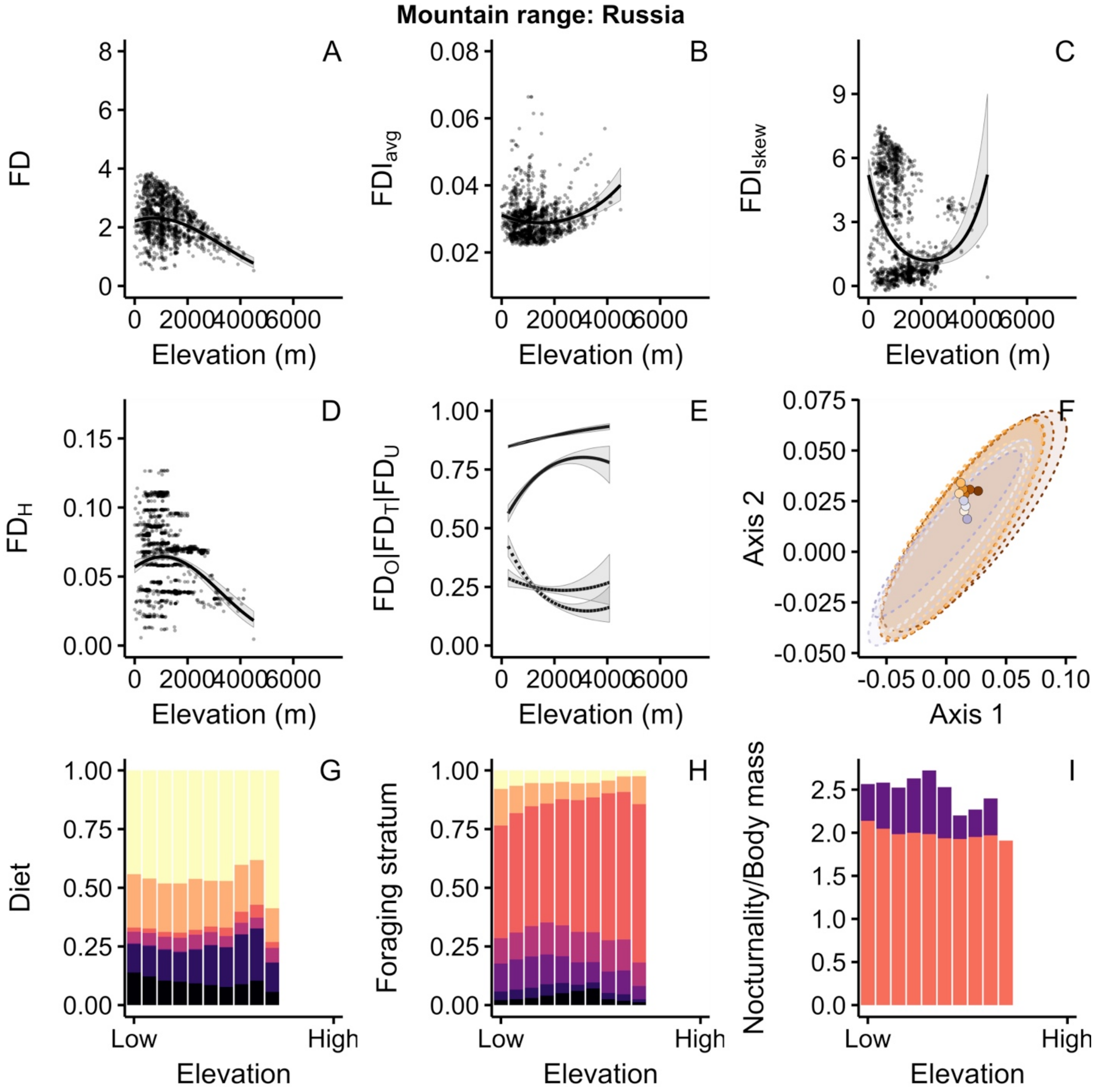

Mountain range: Russia
A
B
C
D
E
F
G
H
I
FD
FDI$_{avg}$
FDI$_{skew}$
FD$_H$
FD$_O$|FD$_T$|FD$_U$
Axis 2
Axis 1
Diet
Foraging stratum
Nocturnality/Body mass
Elevation (m)
Elevation
Low
High

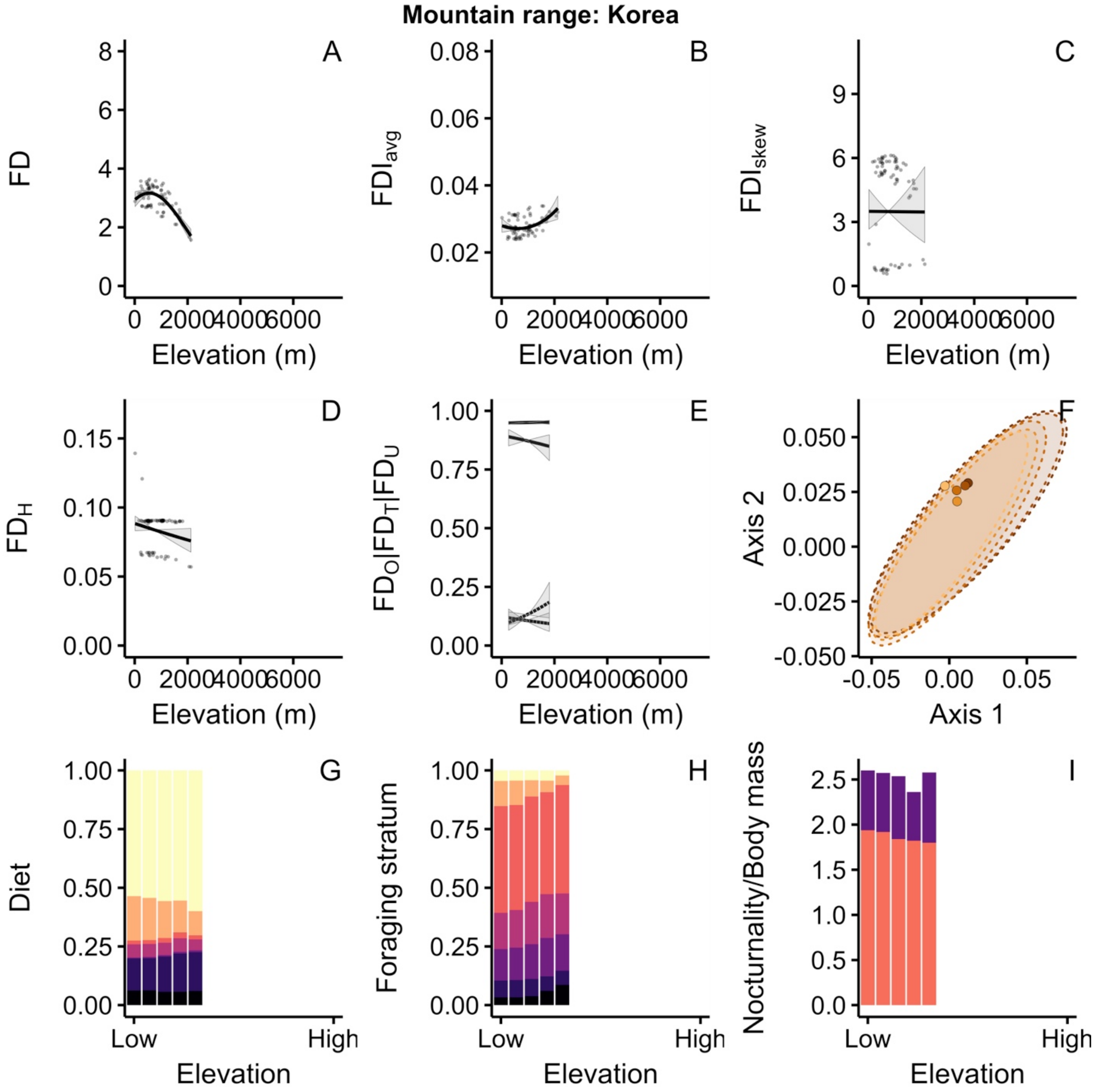
Mountain range: Korea
A
B
C
D
E
F
G
H
I
FD
FDI$_{avg}$
FDI$_{skew}$
FD$_H$
FD$_O$|FD$_T$|FD$_U$
Axis 2
Axis 1
Diet
Foraging stratum
Nocturnality/Body mass
Elevation (m)
Elevation
Low
High

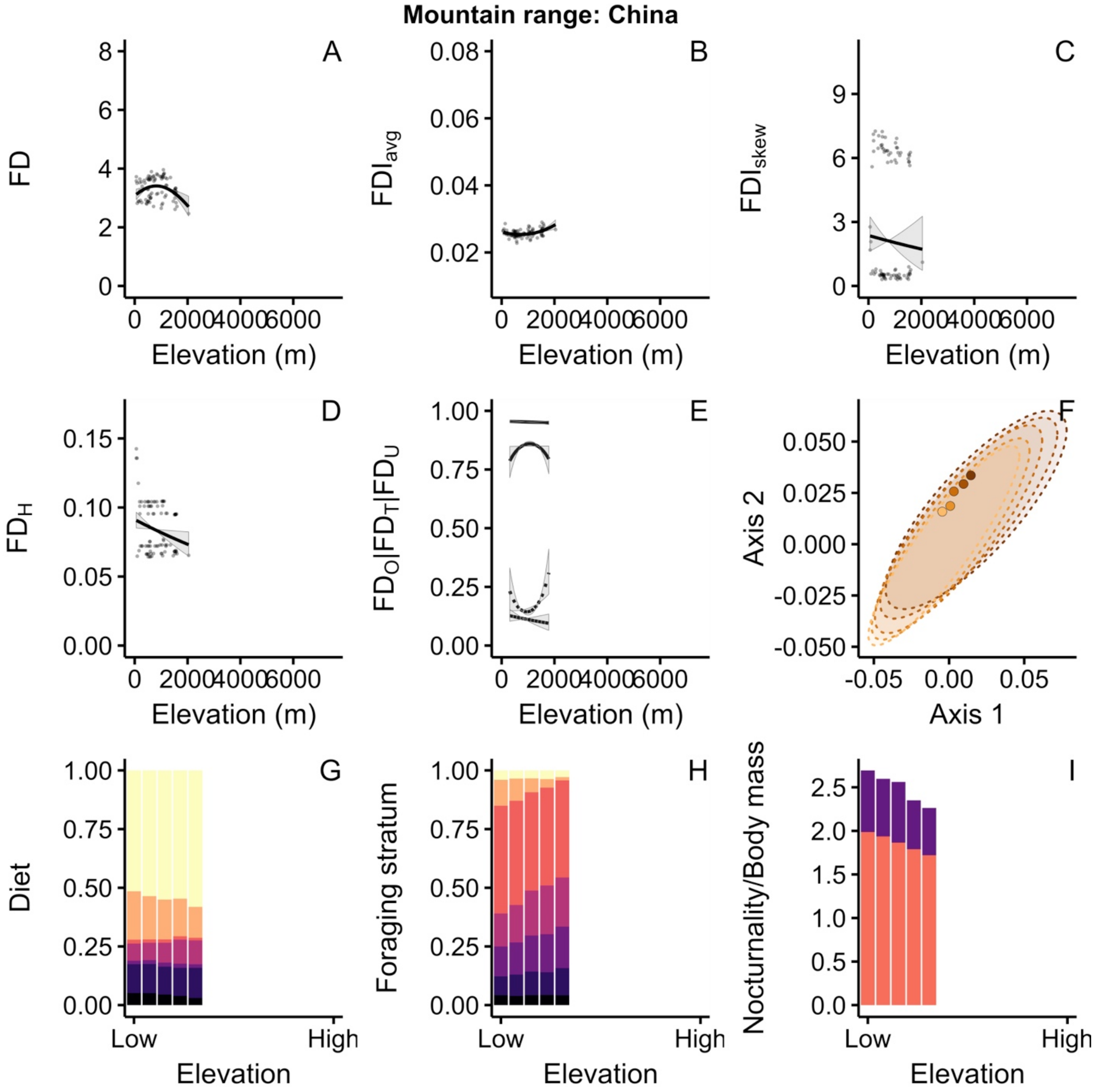

Mountain range: China
A
FD
Elevation (m)
B
$FDI_{avg}$
Elevation (m)
C
$FDI_{skew}$
Elevation (m)
D
$FD_H$
Elevation (m)
E
$FD_O|FD_T|FD_U$
Elevation (m)
F
Axis 2
Axis 1
G
Diet
Low
High
Elevation
H
Foraging stratum
Low
High
Elevation
I
Nocturnality/Body mass
Low
High
Elevation

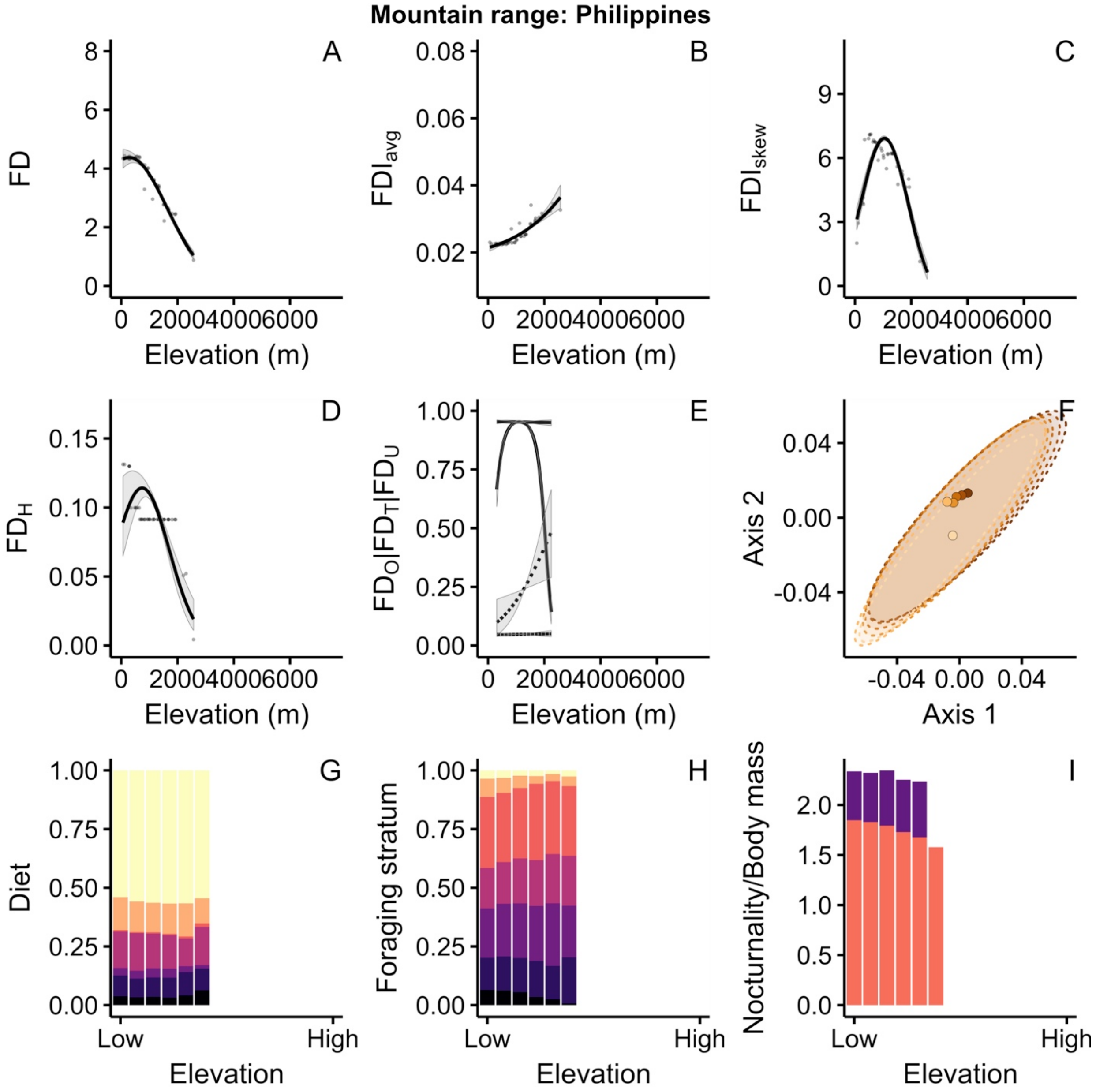

Mountain range: Philippines
A
B
C
D
E
F
G
H
I
FD
FDI$_{avg}$
FDI$_{skew}$
FD$_H$
FD$_O$|FD$_T$|FD$_U$
Axis 2
Axis 1
Diet
Foraging stratum
Nocturnality/Body mass
Elevation (m)
Elevation
Low
High

**Mountain range: Taiwan**

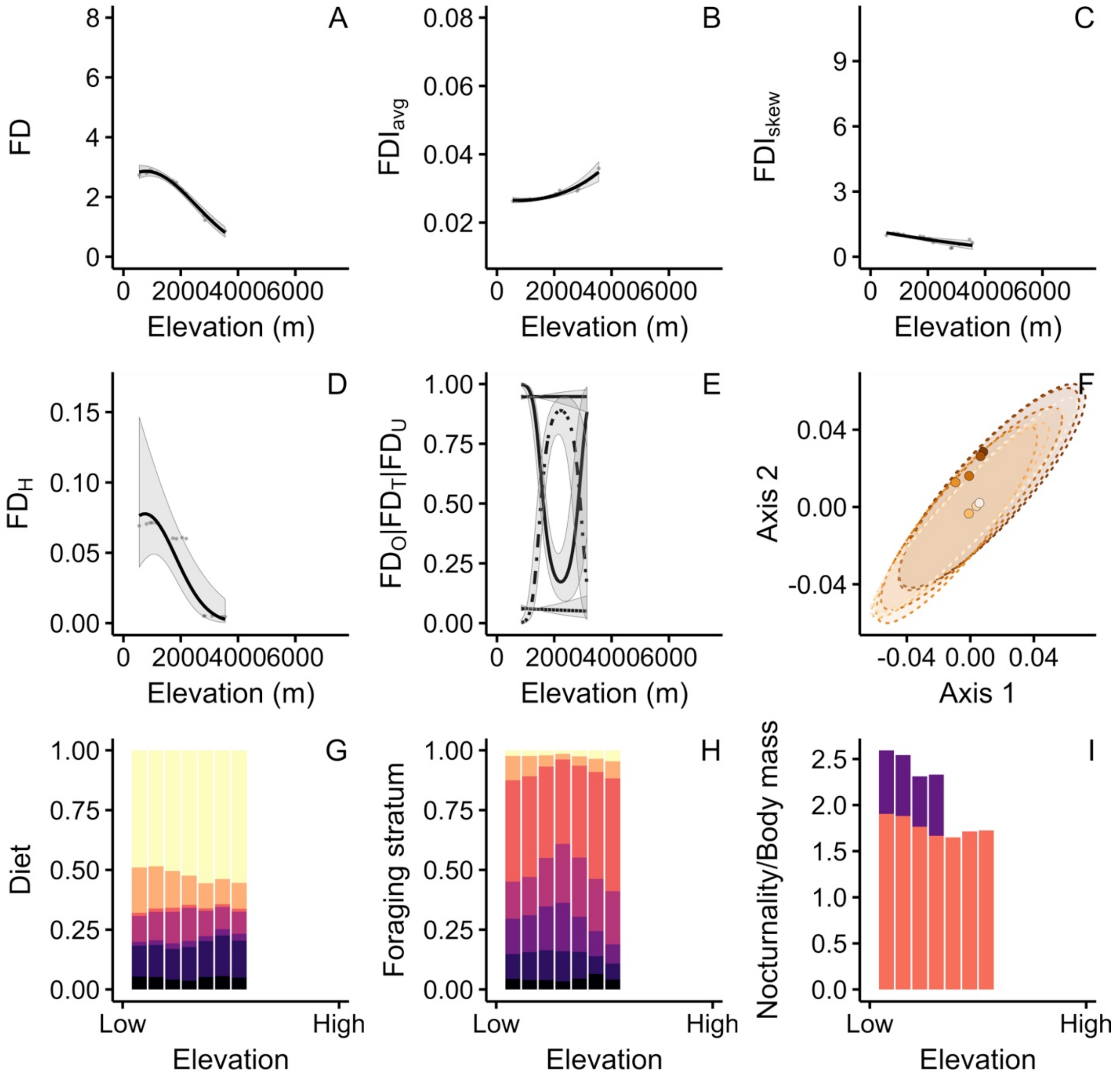

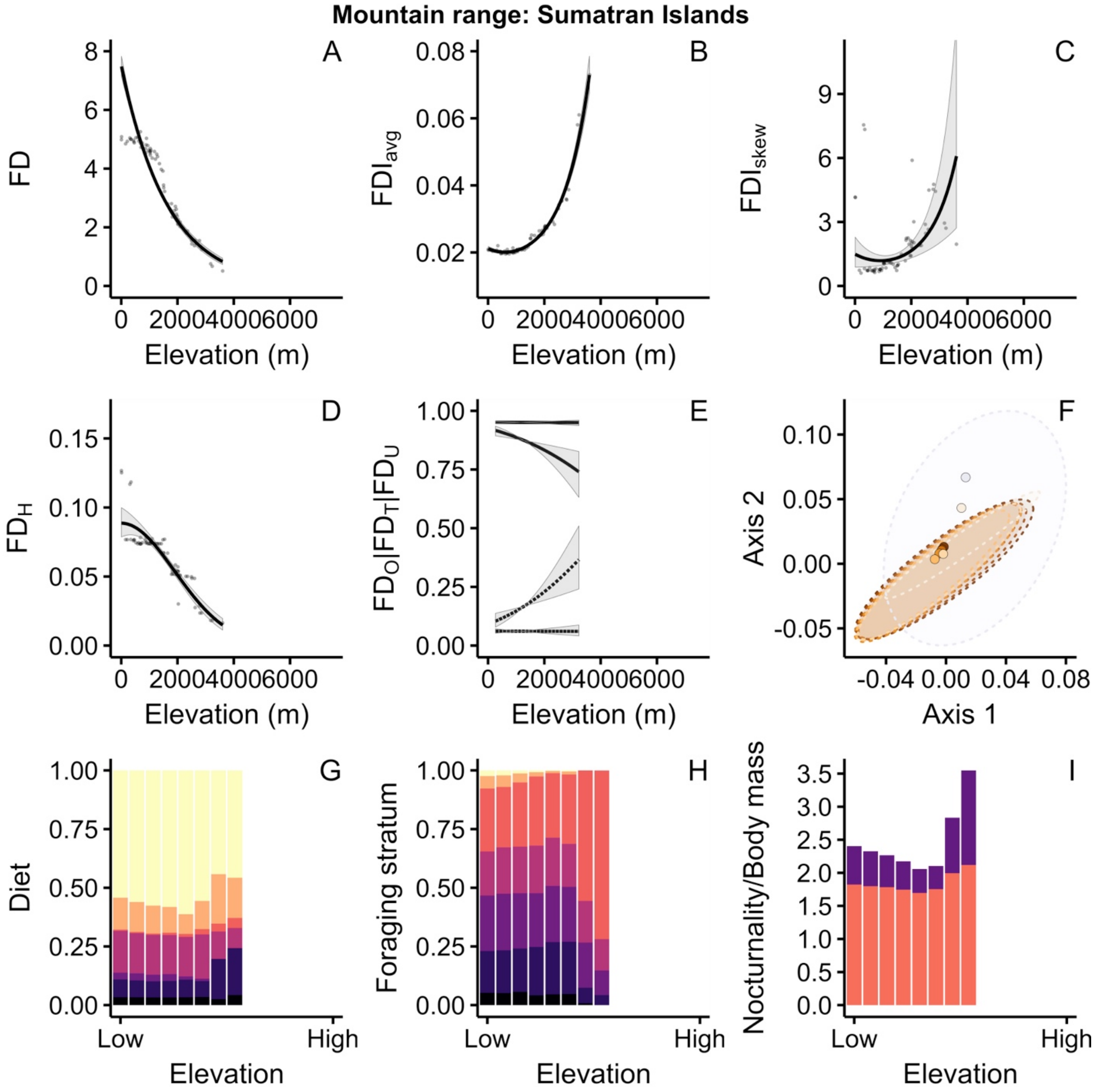
Mountain range: Sumatran Islands
A
B
C
D
E
F
G
H
I
FD
FDI$_{avg}$
FDI$_{skew}$
FD$_H$
FD$_O$|FD$_T$|FD$_U$
Axis 2
Axis 1
Diet
Foraging stratum
Nocturnality/Body mass
Elevation (m)
Elevation
Low
High

**Mountain range: Papua New Guinea**

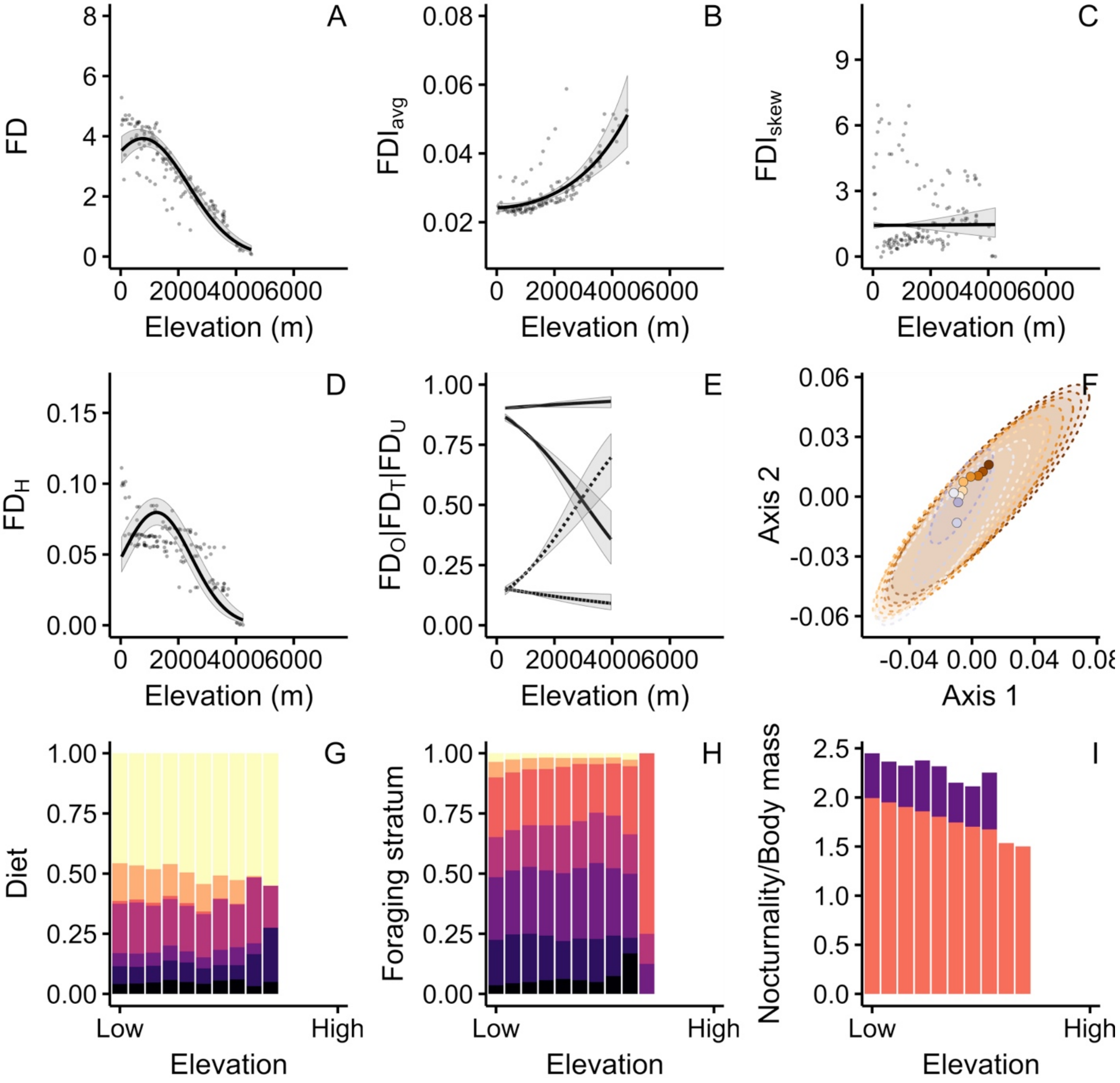

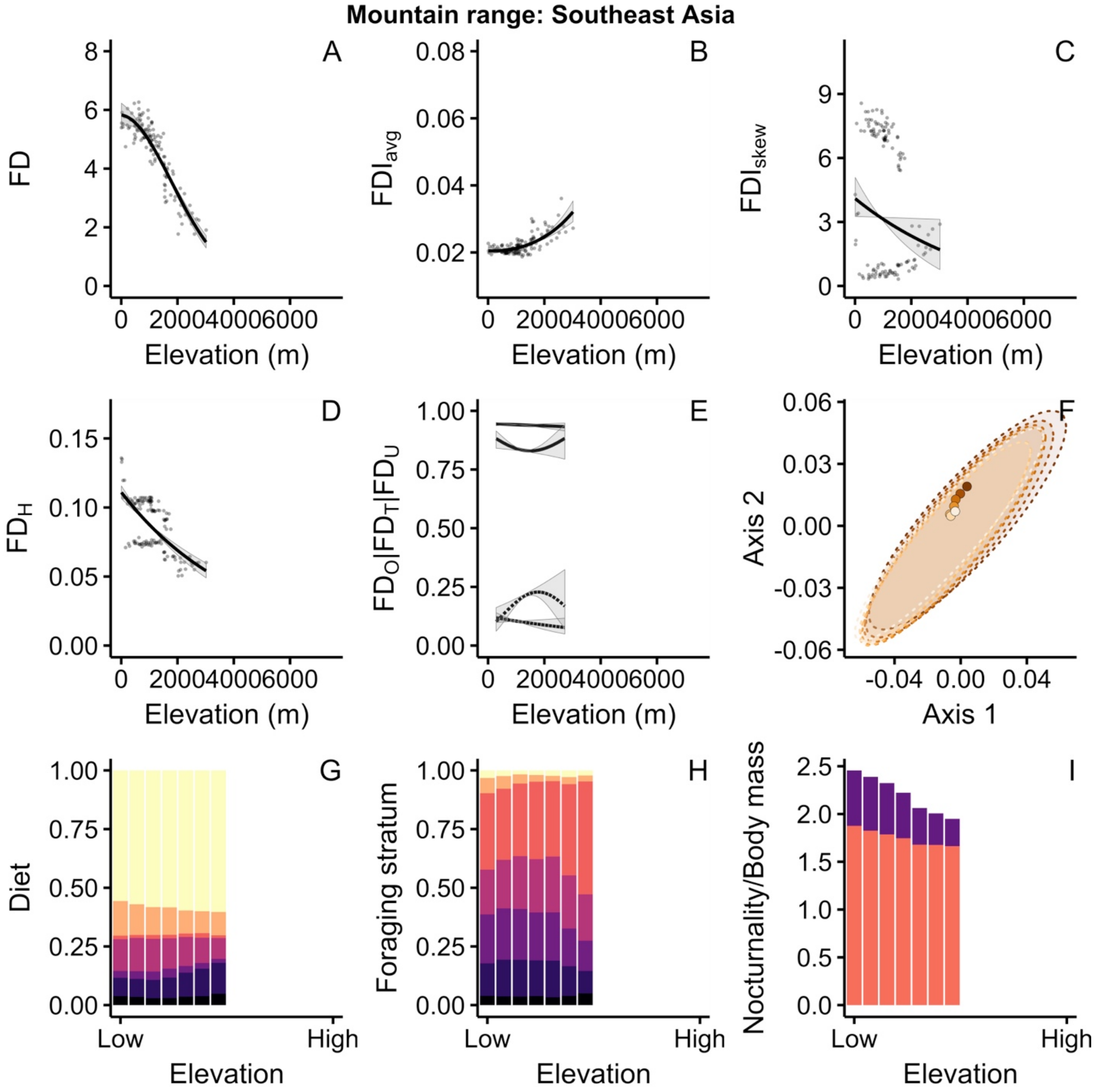

Mountain range: Southeast Asia
A
B
C
D
E
F
G
H
I
FD
FDI$_{avg}$
FDI$_{skew}$
FD$_H$
FD$_O$|FD$_T$|FD$_U$
Axis 2
Axis 1
Diet
Foraging stratum
Nocturnality/Body mass
Elevation (m)
Elevation
Low
High

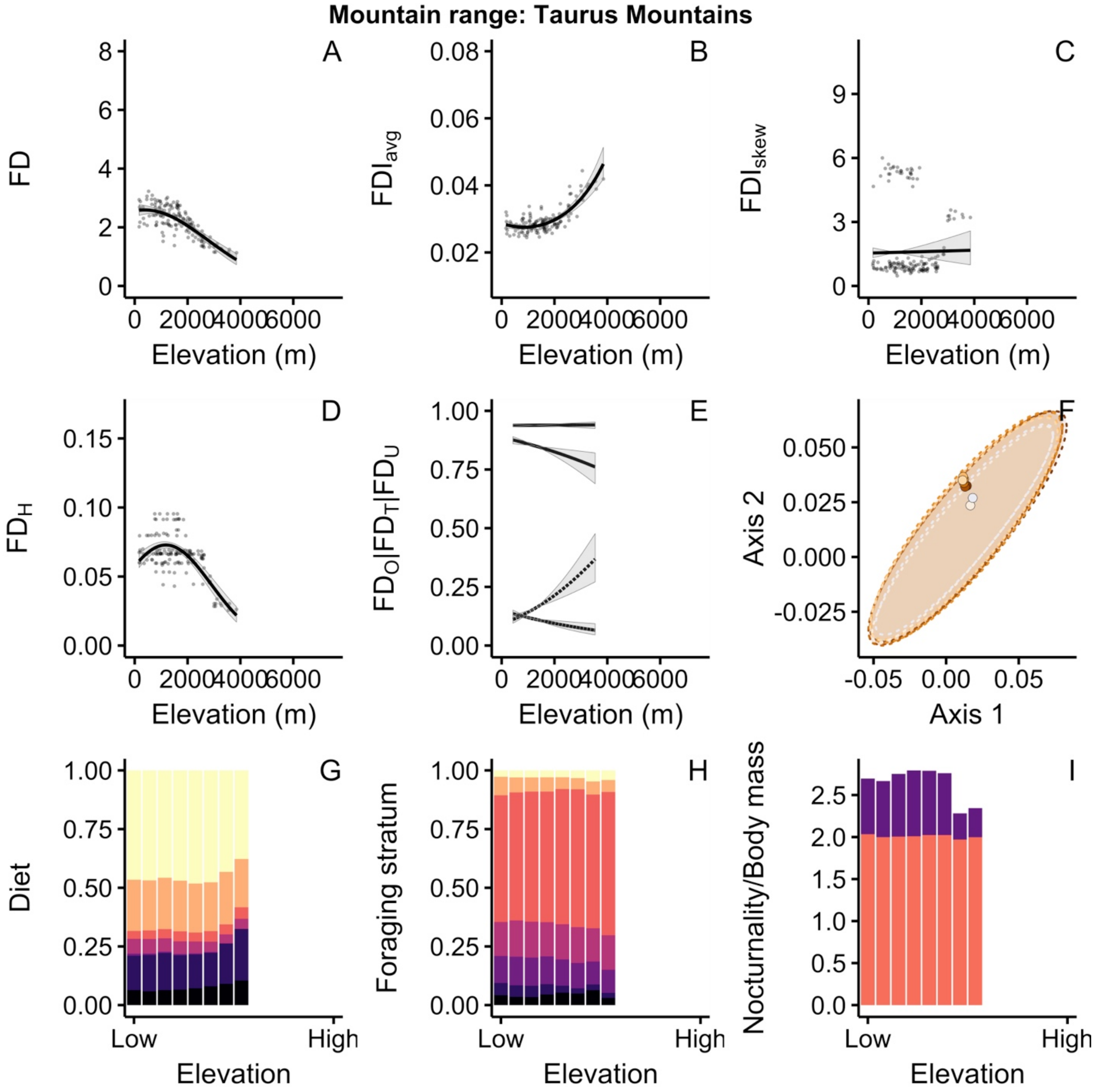
Mountain range: Taurus Mountains
A
B
C
D
E
F
G
H
I
FD
FDI$_{avg}$
FDI$_{skew}$
FD$_H$
FD$_O$|FD$_T$|FD$_U$
Axis 2
Axis 1
Diet
Foraging stratum
Nocturnality/Body mass
Elevation (m)
Elevation
Low
High

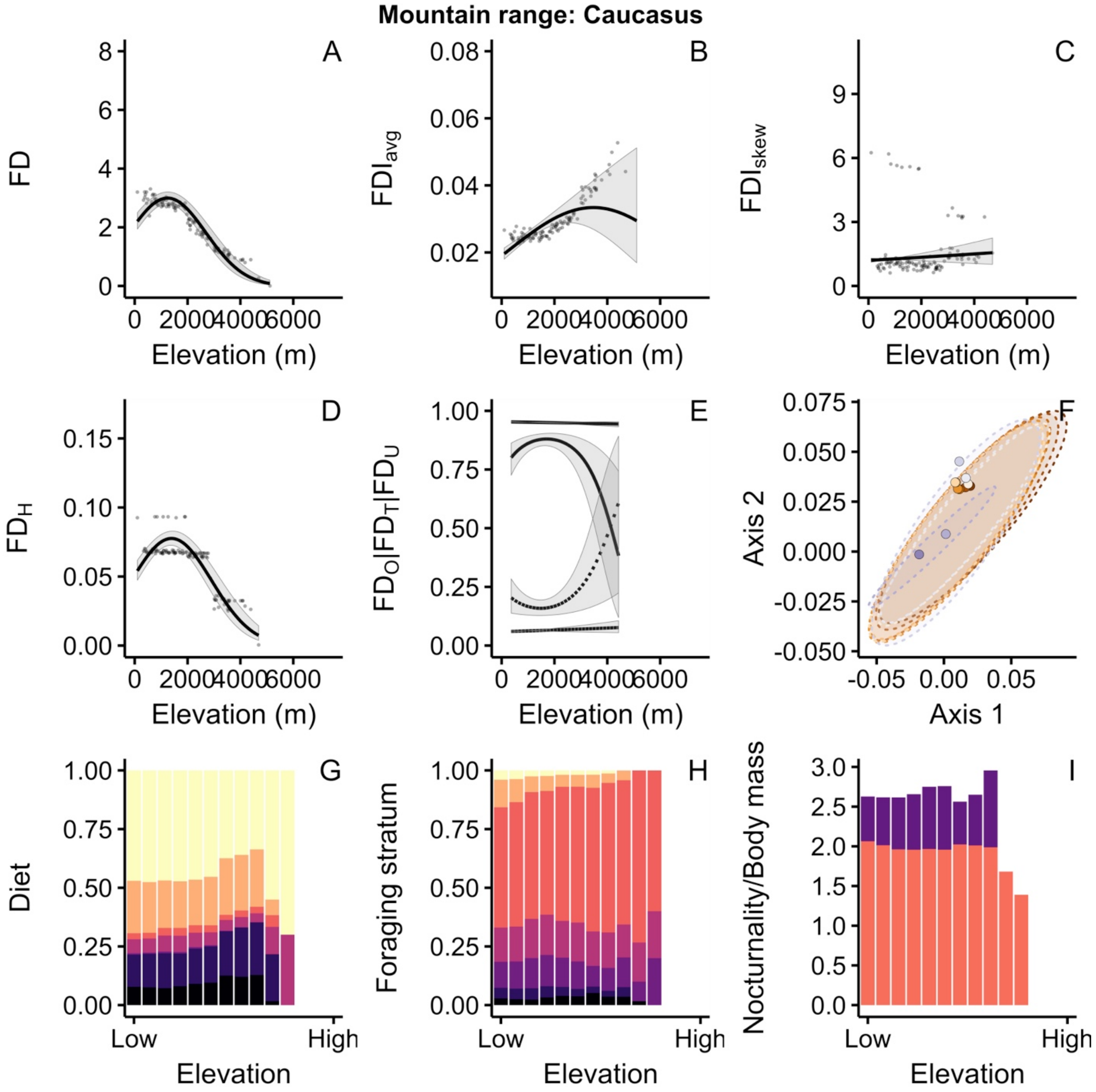
Mountain range: Caucasus
A
B
C
D
E
F
G
H
I
FD
FDI$_{avg}$
FDI$_{skew}$
FD$_H$
FD$_O$|FD$_T$|FD$_U$
Axis 2
Axis 1
Diet
Foraging stratum
Nocturnality/Body mass
Elevation (m)
Elevation
Low
High

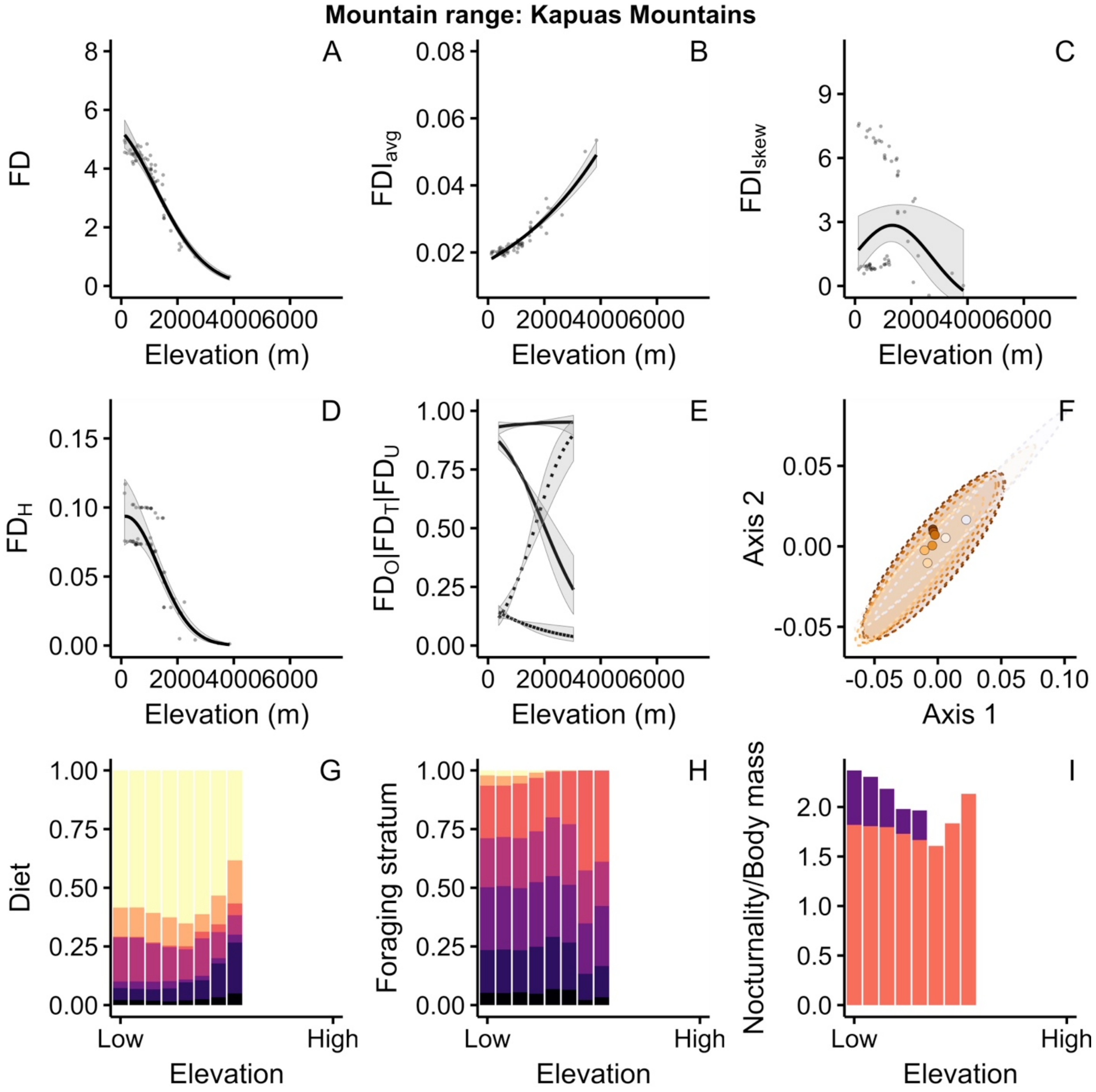

Mountain range: Kapuas Mountains
A
FD
Elevation (m)
B
$FDI_{avg}$
Elevation (m)
C
$FDI_{skew}$
Elevation (m)
D
$FD_H$
Elevation (m)
E
$FD_O|FD_T|FD_U$
Elevation (m)
F
Axis 2
Axis 1
G
Diet
Low
High
Elevation
H
Foraging stratum
Low
High
Elevation
I
Nocturnality/Body mass
Low
High
Elevation

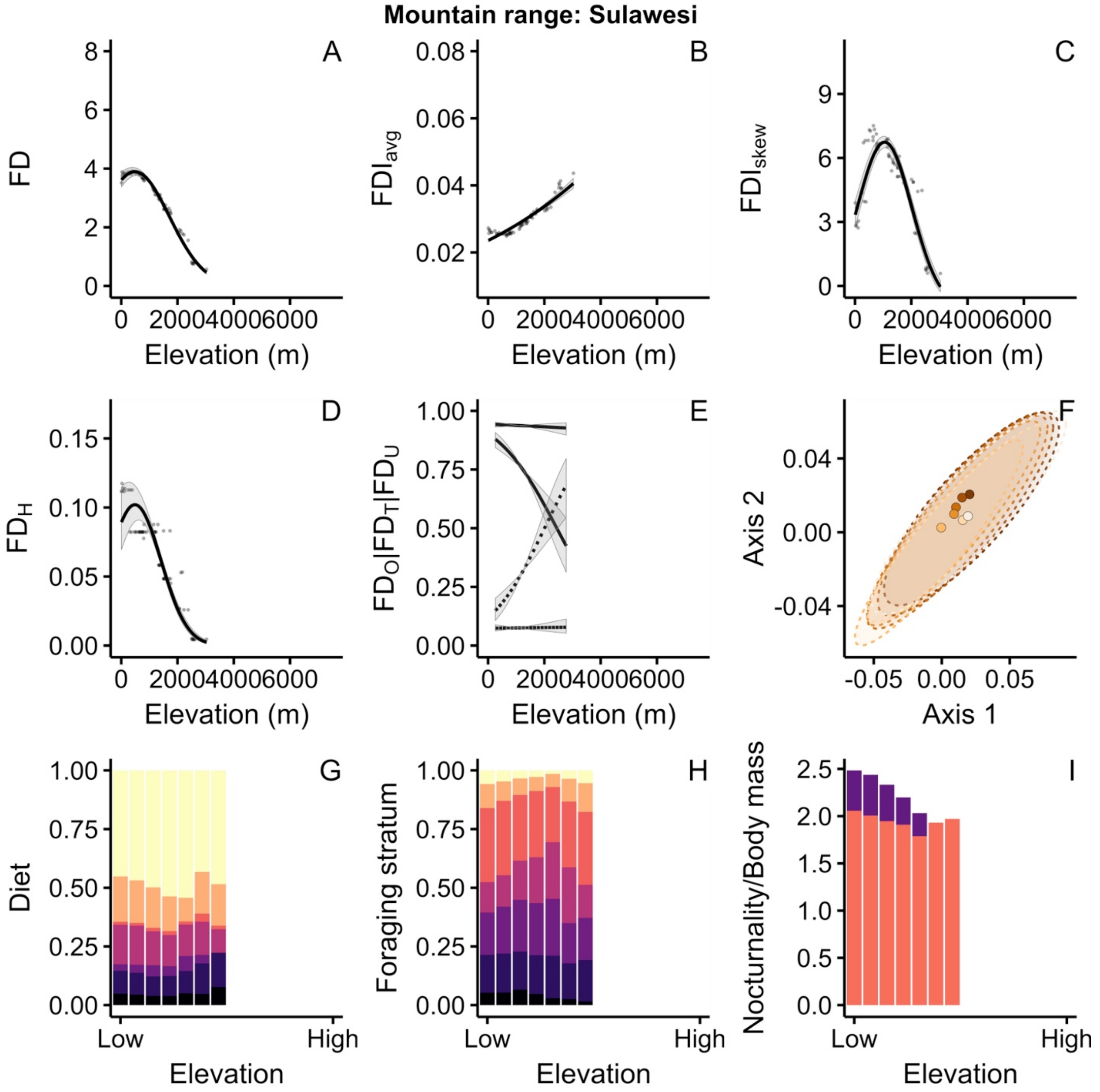
Mountain range: Sulawesi
A
B
C
D
E
F
G
H
I
FD
FDI$_{avg}$
FDI$_{skew}$
FD$_H$
FD$_O$|FD$_T$|FD$_U$
Axis 2
Axis 1
Diet
Foraging stratum
Nocturnality/Body mass
Elevation (m)
Elevation
Low
High

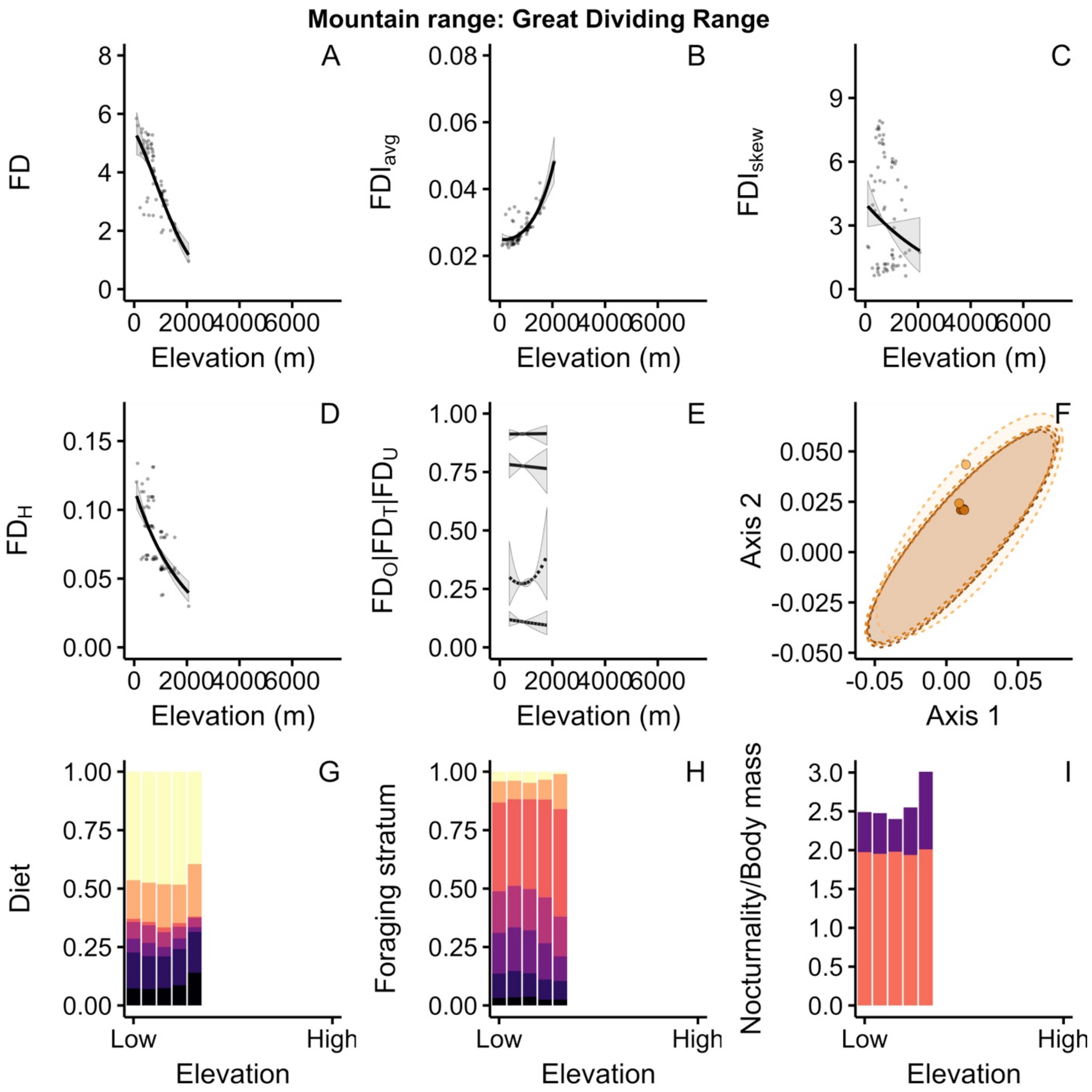

Mountain range: Great Dividing Range
A
B
C
D
E
F
G
H
I
FD
$FDI_{avg}$
$FDI_{skew}$
$FD_H$
$FD_O|FD_T|FD_U$
Axis 2
Axis 1
Diet
Foraging stratum
Nocturnality/Body mass
Elevation (m)
Elevation
Low
High

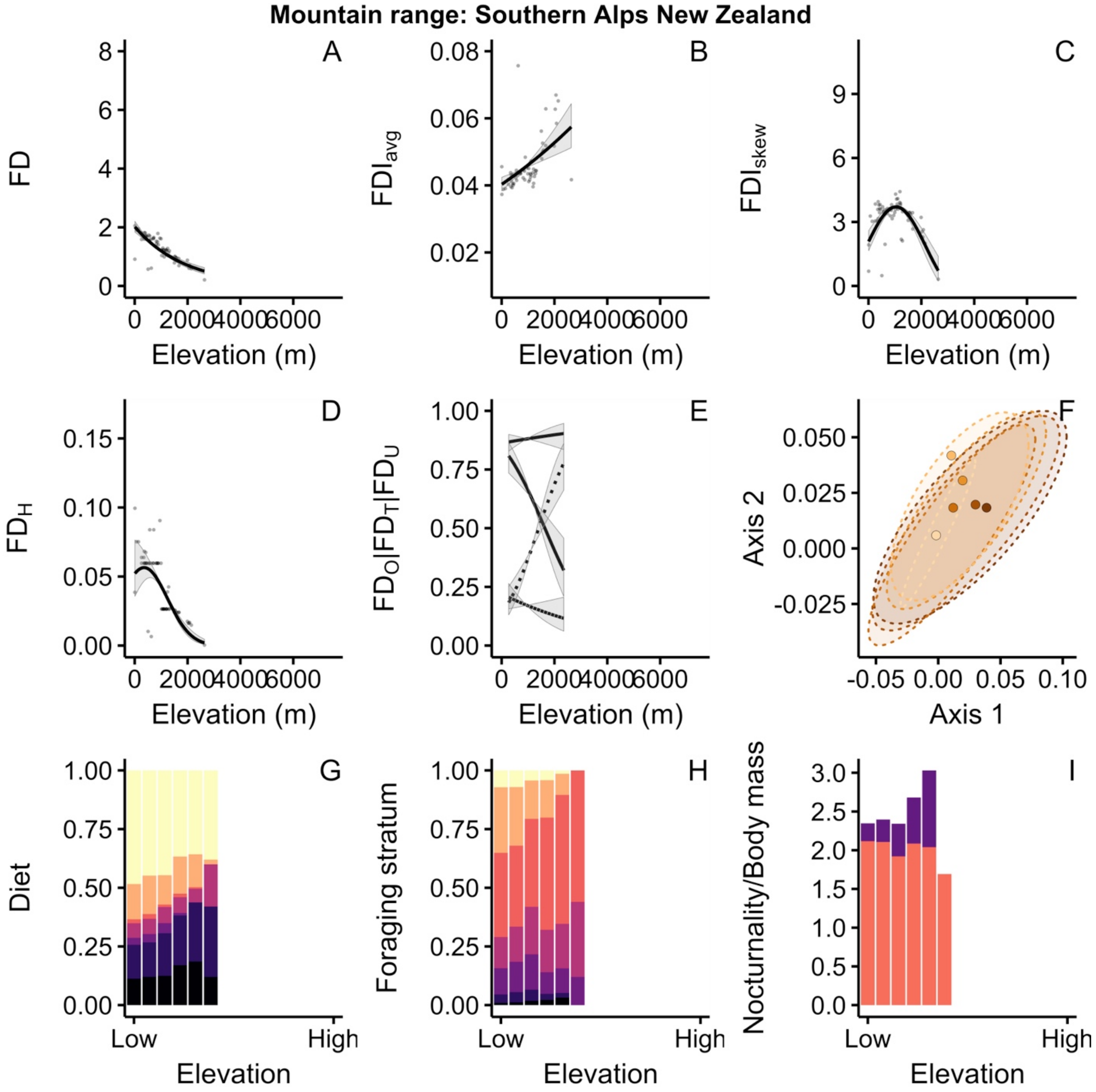
Mountain range: Southern Alps New Zealand
A
B
C
D
E
F
G
H
I
FD
FDI_avg
FDI_skew
FD_H
FD_O|FD_T|FD_U
Axis 2
Axis 1
Diet
Foraging stratum
Nocturnality/Body mass
Elevation (m)
Elevation
Low
High

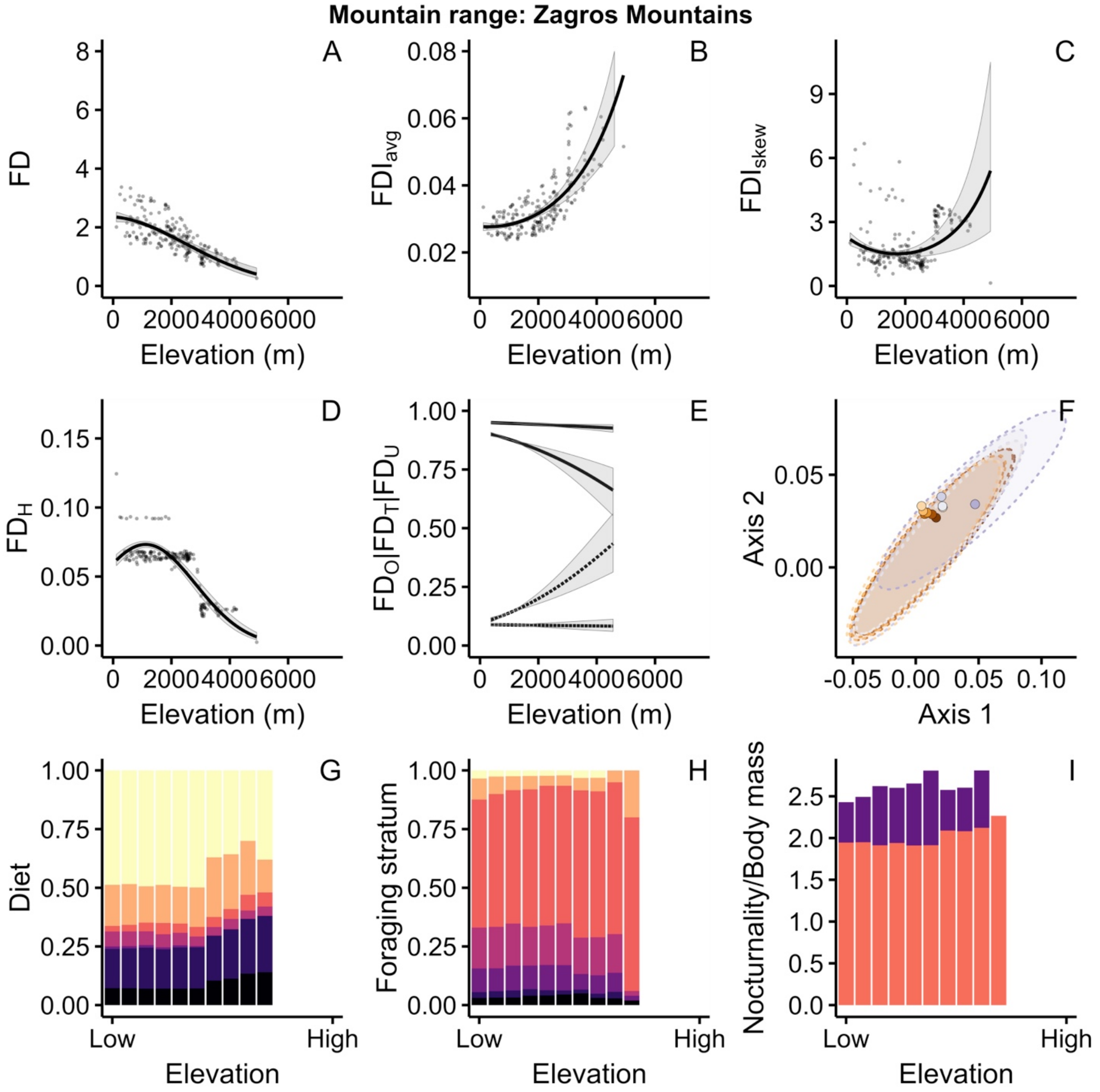
Mountain range: Zagros Mountains
A
B
C
D
E
F
G
H
I
FD
FDI_avg
FDI_skew
FD_H
FD_O|FD_T|FD_U
Axis 1
Axis 2
Diet
Foraging stratum
Nocturnality/Body mass
Elevation (m)
Elevation
Low
High

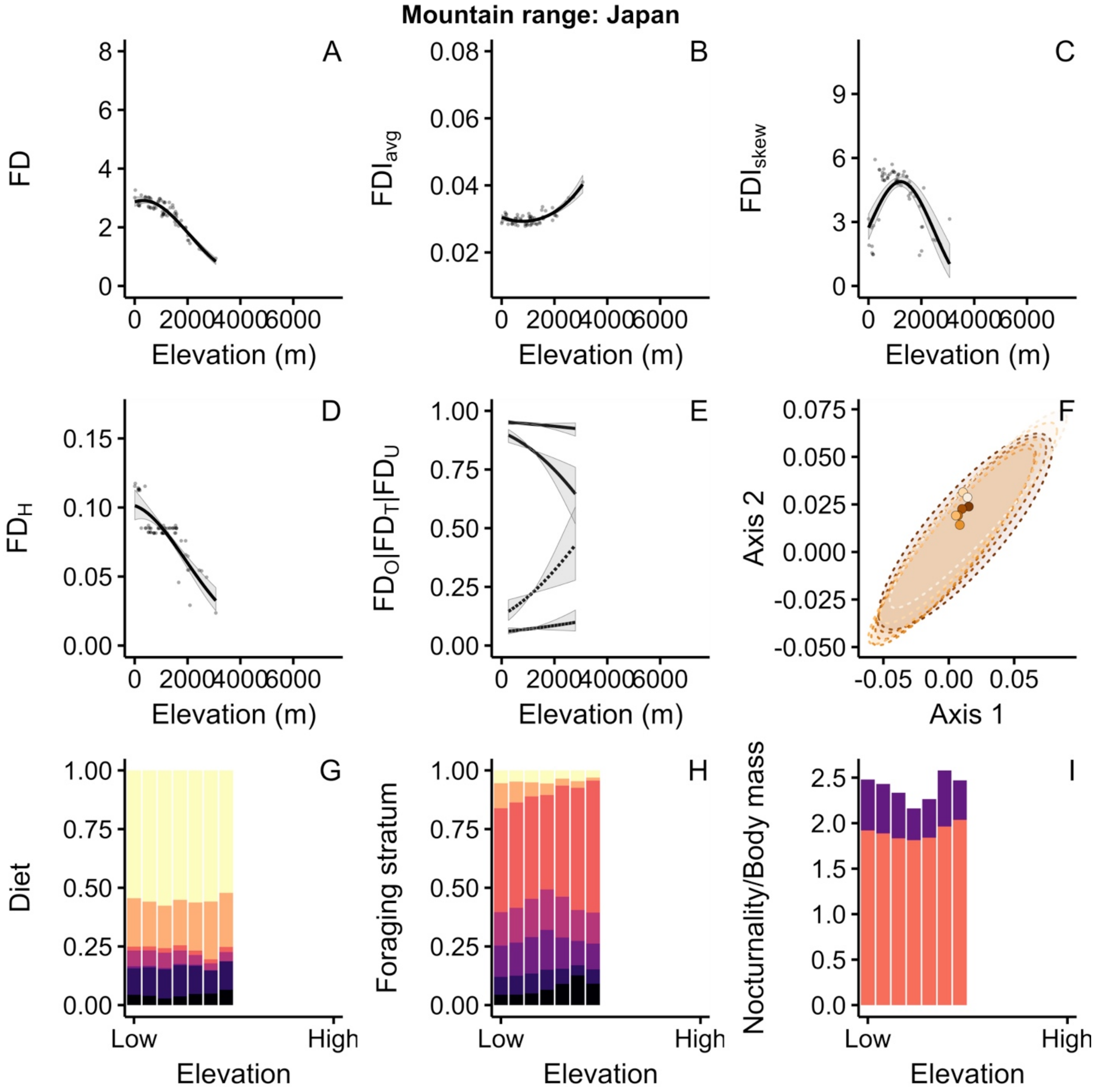
Mountain range: Japan
A
B
C
D
E
F
G
H
I
FD
FDI_avg
FDI_skew
FD_H
FD_O|FD_T|FD_U
Axis 2
Axis 1
Diet
Foraging stratum
Nocturnality/Body mass
Elevation (m)
Elevation
Low
High

Fig. S4. The interaction of central latitude of mountain regions with elevation on gradients of avian functional and phylogenetic diversity, given by predictions from the multi-level models. Specifically, we show predictions for dendrogram-based avian assemblage functional diversity (FD; A), the assemblage mean ($FDI_{avg}$; B) and skewness ($FDI_{skew}$; C) of species' local functional distinctness underpinning it, and phylogenetic diversity (PD; D). Grey areas are 95% credible intervals. For details see on multi-level models Materials and Methods.

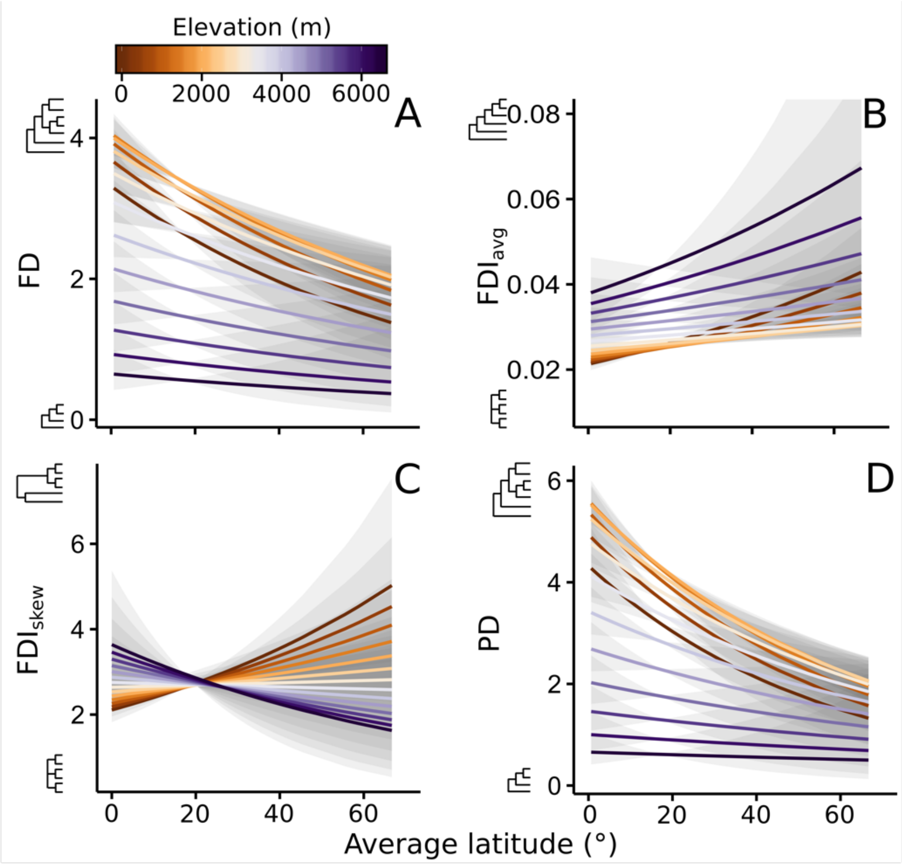

Fig. S5. The frequency with which assemblage phylogenetic structure accurately predicts functional structure across the latitudinal gradient. Frequency of successful predictions is given by the proportion of phylogenetically overdispersed (clustered) assemblages that are also functionally overdispersed (clustered). Clustering and overdispersion are given by p-values of 0.025 and 0.975, respectively, estimated from the quantile scores for the observed values of FD and PD (see text for details). To account for the effect of separate mountain ranges, the frequency of accurate predictions was averaged per mountain region; mountain region's central latitude was used to evaluate the latitudinal gradient in the frequency of accurate predictions.

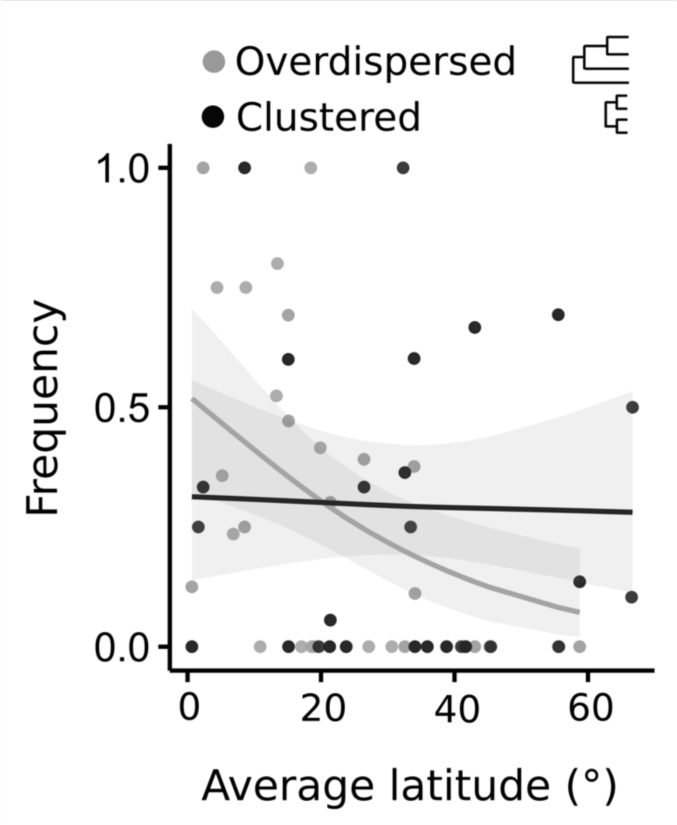

Fig. S6. Null-model expectation of the elevational gradients of the avian trait space for (A) all components of the dietary and (B) foraging stratum axes, (C) nocturnality, and (D) assemblage mean body mass.

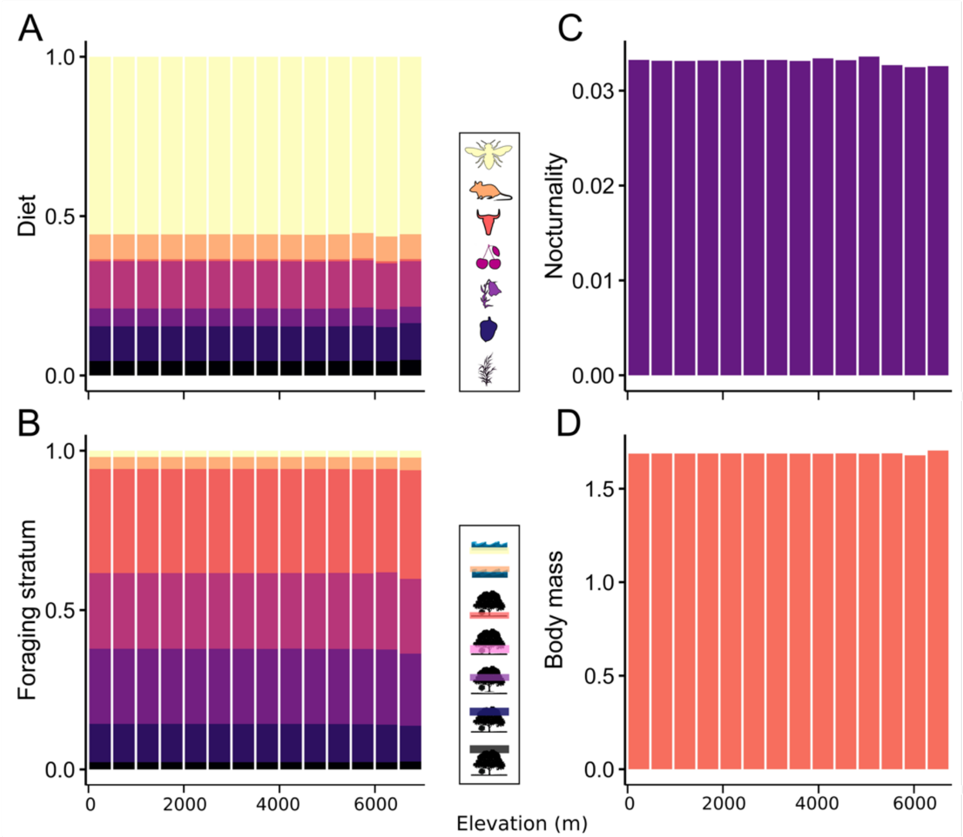

Fig. S7. The interaction of central latitude of mountain regions with elevational gradients of dietary traits. Shown are proportions of insectivorous (A), vertebrate (B), scavenging (C), frugivorous (D), nectarivorous (E), granivorous (F), and plant matter (G) diets. Fitted aggregate global pattern combining all 8,410 assemblages, with mountain ranges included as random effects in the model, are shown in red dashed line. Grey areas are 95% credible intervals. For five highlighted regions, see Fig. 1. For details on the models, see Materials and Methods.

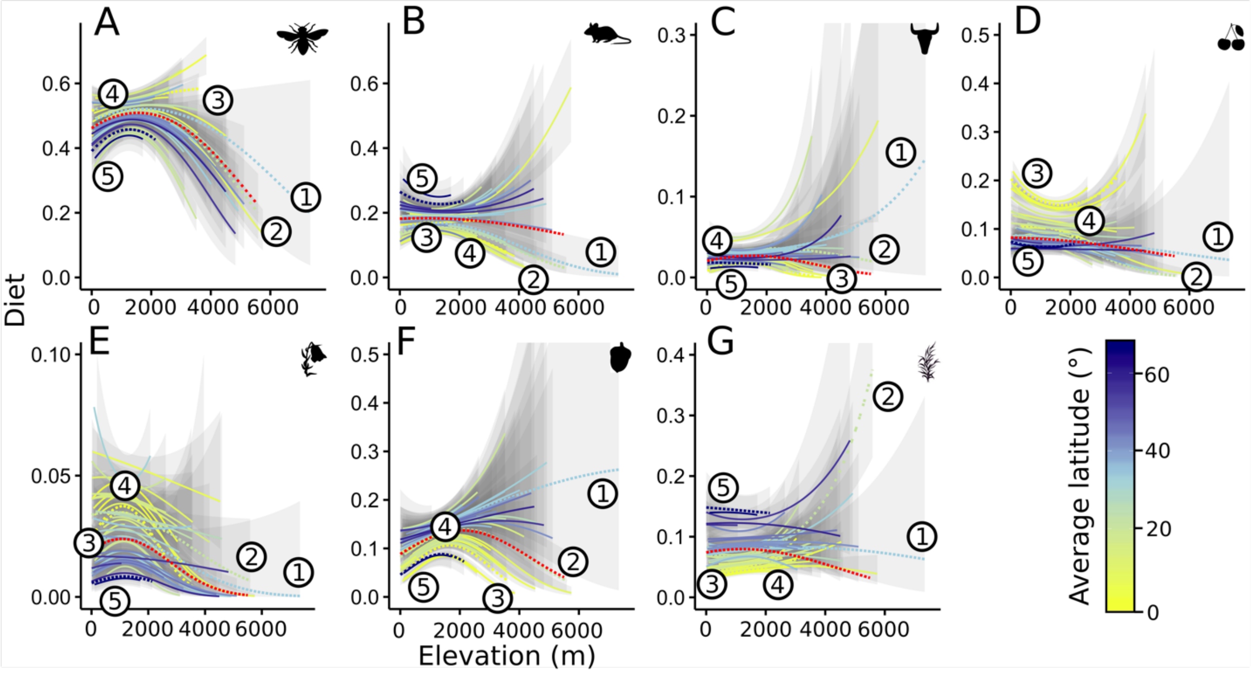

Fig. S8. The interaction of central latitude of mountain regions with elevational gradients of foraging stratum traits. Shown are proportions of water below surface (A), water around surface (B), graound (C), understory (D), mid canopy (E), upper canopy (F), and aerial (G) foraging strata. Fitted aggregate global pattern combining all 8,410 assemblages, with mountain ranges included as random effects in the model, are shown in red dashed line. Grey areas are 95% credible intervals. For five highlighted regions, see Fig. 1. For details on the models, see Materials and Methods.

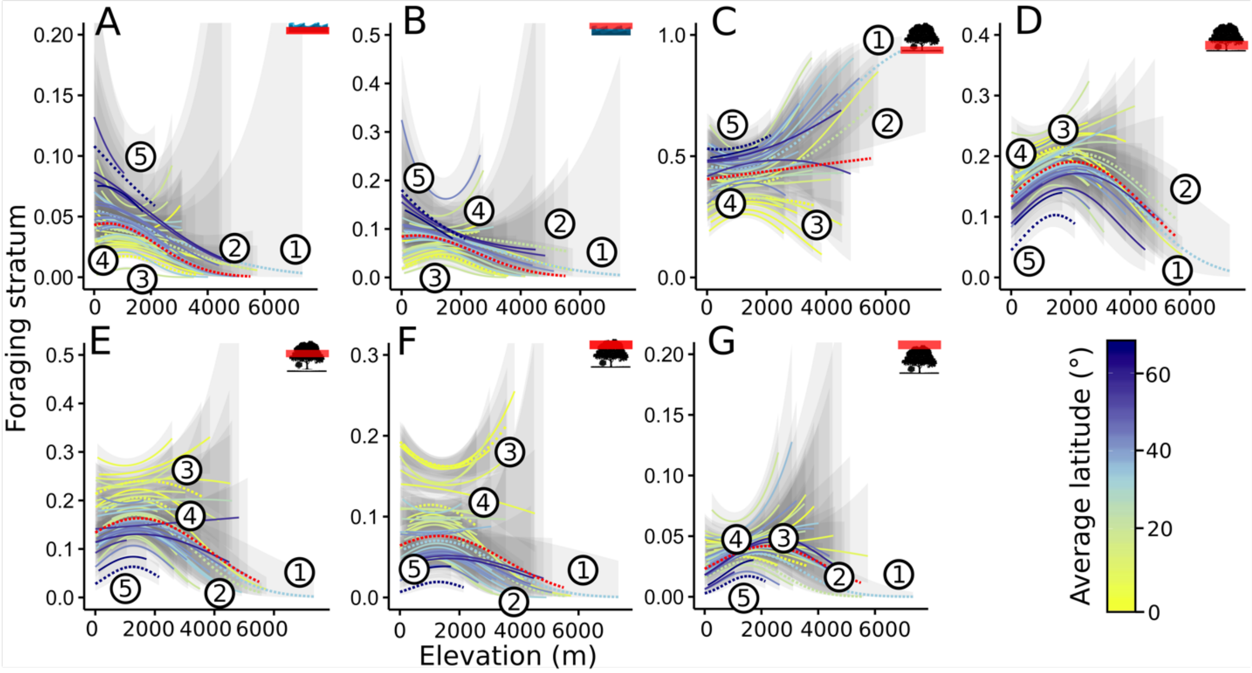

Fig. S9. Elevational gradients of global patterns combining all 8,410 assemblages of the overlap in trait space ($FD_O$) measured as Sørensen similarity, true turnover ($FD_T$) measured as Simpson's similarity, and fractions of trait space unique to assemblages located at lower ($FD_{Ul}$) and higher ($FD_{Uh}$) elevations. $FD_O$, $FD_T$, $FD_{Ul}$, and $FD_{Uh}$ were calculated for pairs of points located at an elevation *a* (point 1) and the geographically closest point (point 2) located within (A) (*a*+250m, *a*+750m) and (B) (*a*+750m, *a*+1250m) elevational region.

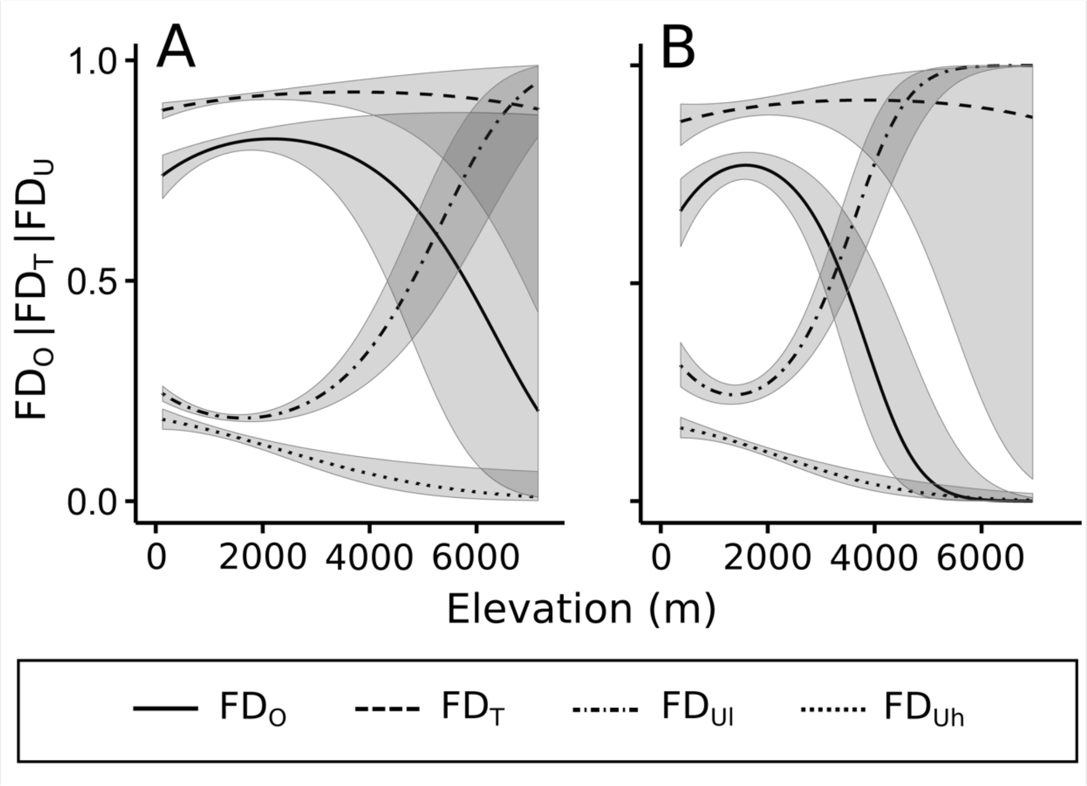
